# CoVAMPnet: Comparative Markov State Analysis for Studying Effects of Drug Candidates on Disordered Biomolecules

**DOI:** 10.1101/2023.01.06.523007

**Authors:** Sérgio M. Marques, Petr Kouba, Anthony Legrand, Jiri Sedlar, Lucas Disson, Joan Planas-Iglesias, Zainab Sanusi, Antonin Kunka, Jiri Damborsky, Tomas Pajdla, Zbynek Prokop, Stanislav Mazurenko, Josef Sivic, David Bednar

## Abstract

Computational study of the effect of drug candidates on intrinsically disordered biomolecules is challenging due to their vast and complex conformational space. Here we developed a Comparative Markov State Analysis (CoVAMPnet) framework to quantify changes in the conformational distribution and dynamics of a disordered biomolecule in the presence and absence of small organic drug candidate molecules. First, molecular dynamics trajectories are generated using enhanced sampling, in the presence and absence of small molecule drug candidates, and ensembles of soft Markov state models (MSMs) are learned for each system using unsupervised machine learning. Second, these ensembles of learned MSMs are aligned across different systems based on a solution to an optimal transport problem. Third, the directional importance of inter-residue distances for the assignment to different conformational states is assessed by a discriminative analysis of aggregated neural network gradients. This final step provides interpretability and biophysical context to the learned MSMs. We applied this novel computational framework to assess the effects of ongoing phase 3 therapeutics tramiprosate (TMP) and its metabolite 3-sulfopropanoic acid (SPA) on the disordered Aβ42 peptide involved in Alzheimer’s disease. Based on adaptive sampling molecular dynamics and CoVAMPnet analysis, we observed that both TMP and SPA preserved more structured conformations of Aβ42 by interacting non-specifically with charged residues. SPA impacted Aβ42 more than TMP, protecting α-helices and suppressing the formation of aggregation-prone β-strands. Experimental biophysical analyses showed only mild effects of TMP/SPA on Aβ42, and activity enhancement by the endogenous metabolization of TMP into SPA. Our data suggest that TMP/SPA may also target other biomolecules than Aβ peptides. The CoVAMPnet method is broadly applicable to study the effects of drug candidates on the conformational behavior of intrinsically disordered biomolecules.

**TOC Graphic:** 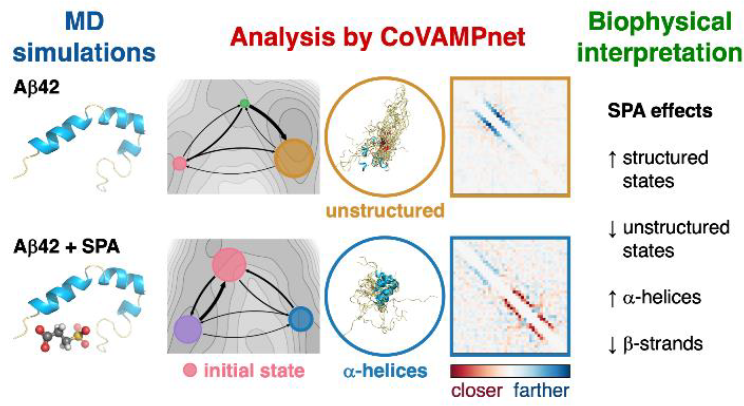

## INTRODUCTION

Alzheimer’s disease (AD) is globally the fifth leading cause of death and fourth cause of disability in people aged 75 years and above, and thus represents an enormous societal burden^1^. Amyloid-beta (Aβ) peptides play a major role in the development of AD, although the mechanism behind their toxicity is still debated^2,3^. A model of toxicity known as the oligomer hypothesis states that Aβ oligomerizes into toxic pore-forming oligomers at the neuronal plasma membrane, which ultimately leads to cell death. Among the different Aβ peptides, the 42-residue long (Aβ42; **Figure 1A**) is the most aggregation-prone isoform^4,5^.

**Figure 1.**
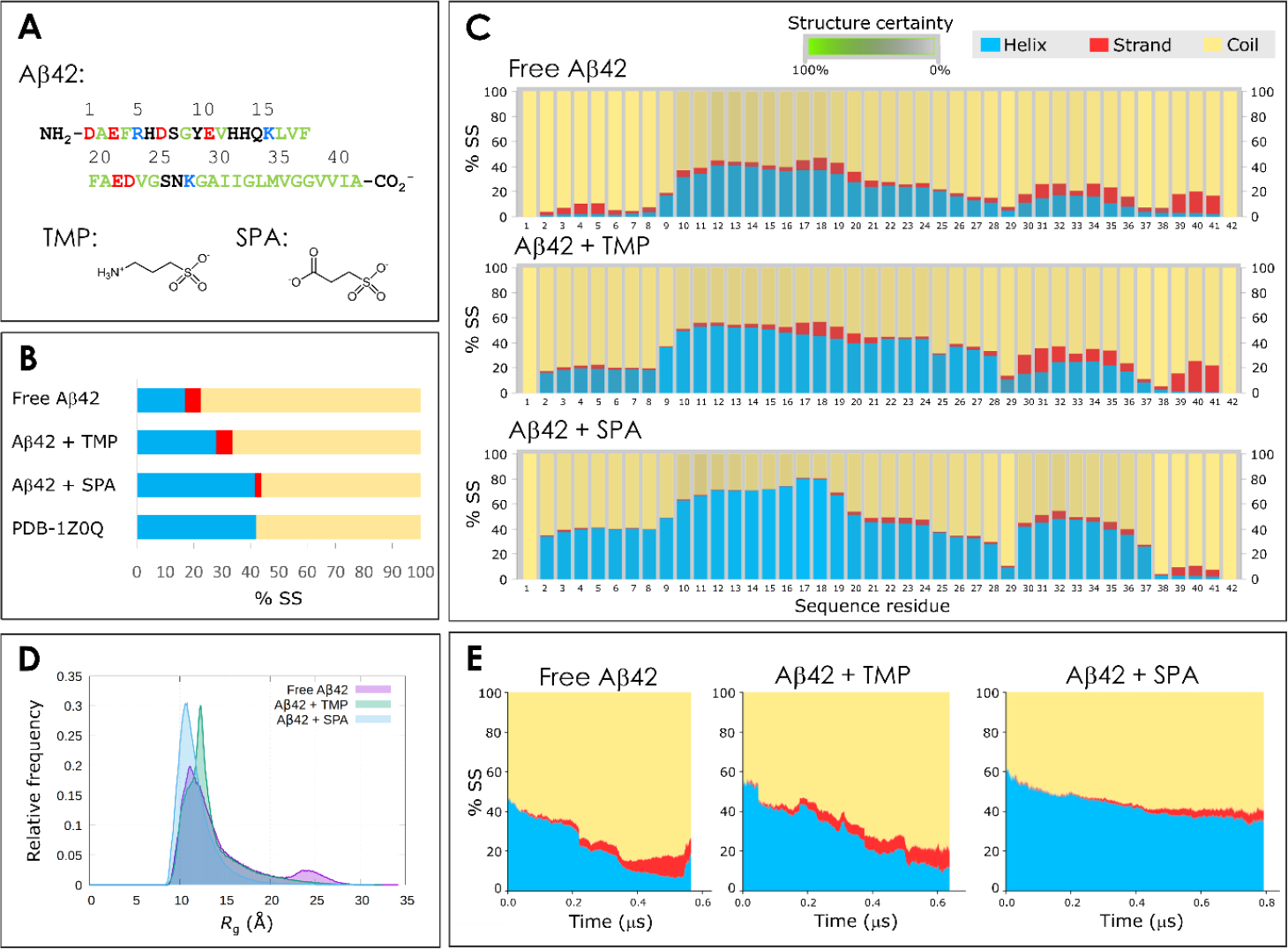
Structures of Aβ42 peptide and the studied small molecules, and properties of the ensembles from the adaptive simulations for the free Aβ42, Aβ42 + TMP and Aβ42 + SPA. A) Sequence of the Aβ42 peptide and chemical structures of tramiprosate (TMP) and 3-sulfopropanoic acid (SPA) in the dominant protonation states at the physiological pH 7.4. The sequence residues are color-coded as follows: *red* for negatively charged; *blue* for positively charged; *green* for hydrophobic; and *black* for polar neutral residues. B) Total secondary structural propensity (% SS) of Aβ42 during the adaptive MDs, in the original NMR ensemble (PDB 1Z0Q with 30 structures), and from the experimental measurements of free Aβ42 in aqueous solution. C) Secondary structure propensity of Aβ42 by residue, obtained for the global ensembles from the adaptive simulations. The certainty of the secondary structure assignment was obtained by the statistical variance among ten randomized bins of frames, and is represented by the saturation of the secondary structure color (the more saturated the color, the more certain the assignment, as indicated by the legend). D) Distribution of the radius of gyration (*R*_g_) of the ensembles from the same adaptive simulations. E) Time evolution of the secondary structure of Aβ42 during the time-based aligned adaptive sampling MD simulations. The secondary elements are aggregated across all 42 residues, averaged at each time over all the trajectories parallel in time according to the time-based alignment. Only the timespan covering at least 20 parallel trajectories is plotted.

The Aβ peptides are intrinsically disordered, which makes them difficult to study both experimentally and computationally. Intrinsically disordered proteins do not adopt a single well-defined structure, but rather exist as ensembles of conformations with similar energies. These ensembles are best characterized by their population distributions and probabilities of several properties or descriptors (e.g., radius of gyration and secondary structure)^6,7^. The disordered nature of Aβ42 significantly complicates the analysis of its molecular dynamics (MD) trajectories, namely the definition of conformational states, which is an important step towards a deeper understanding of the system and its slowest transitions^8^. A popular approach for identifying notable conformational states in MD simulations involves building so-called Markov state models (MSMs). Under the assumption of the dynamics being Markovian (memoryless), these models cluster the conformational space into states preserving the Markovianity of the transitions and estimate the equilibrium distribution and transition rates between the states. The conventional methods for building MSMs typically rely on a selection of collective variables, compressing the high-dimensional MD data and simplifying the clustering. Recent progress in variational approaches for conformational dynamics has further allowed scoring different MSMs, e.g., based on their ability to approximate the slowest modes of the dynamics, thus facilitating the development of automatic frameworks for the identification of Markov states^9^. Although some of these procedures are quite advanced and enable, e.g., an accurate estimation of transition rates even from biased simulation data^10^, the manual selection of the collective variables is typically laborious and can often cause the resulting models to fail the tests for Markovianity. While MSMs are extremely valuable tools, they possess certain limitations, such as the assumption of Markovianity, constraints on state representation granularity, reliance on extensive sampling, and relatively rapid relaxation dynamics^11–14^. Several alternative methodologies exist to address these shortcomings. These include hidden Markov models (HMMs) to relax the Markovian assumption^12^, approaches incorporating memory effects such as the generalized master equation (GME) and the generalized Langevin equation (GLE) for more effective dynamic property assessment^11^, and methods rooted in deep learning^15^.

A powerful framework based on deep learning is VAMPnet, a neural network that learns a probabilistic assignment of each simulation frame to individual states in an unsupervised manner by maximizing a variational score^16^. In contrast to the other methods, the VAMPnet approach does not relax the Markovianity assumption but rather combines the search of collective variables with the optimization of a cost function to efficiently identify the slowest modes of the system. The application of VAMPnets to the analysis of Aβ42 trajectories has already shown great potential in producing robust MSMs for quantification of the Aβ42 kinetics and equilibrium properties^17^. Several recent methods build on the VAMPnet approach to address the efficiency of protein representation^18,19^, scalability to multi-domain protein systems^20^, stability of the training process^21^, sampling of rare conformations^22^, or the importance of residues based on the attention mechanism^18,23^. However, to the best of our knowledge, a method for aligning and comparing ensembles of learned MSMs across different systems that would simplify the biophysical interpretation of the conformational states by identifying their distinctive features is still missing. In this work, we develop such a method to help understand and quantify the effects of drug candidates on the conformational space of the analyzed system.

This problem is important in many fields of research, particularly in AD. Due to the prevalence and severity of the disease, there is a growing interest in pharmaceuticals capable of preventing the early stages of the Aβ42 oligomerization and stopping the pathogenic amyloid cascade^3,4,24^. Tramiprosate (TMP), also known as homotaurine or 3-amino-1-propane sulfonic acid, is a naturally occurring aminosulfonate. Even at high concentrations, it is well tolerated in the human brain, where it is metabolized into 3-sulfopropanoic acid (SPA) (**Figure 1A**). TMP has been reported to prevent the formation of fibrillar forms of Aβ, reduce the Aβ-induced death rate of neuronal cell cultures, and lower the amyloid plaque deposition in the brain^25–27^. Clinical trials have shown its ability to slow down the cognitive decline in patients with homozygous expression of the apolipoprotein E gene *APOE4*, similarly to FDA-approved aducanumab^24,28^. TMP can act not only on Aβ, but also on other pathways that contribute to cognitive impairment in AD and other neurologic disorders^29,30^. ALZ-801 is a valine-conjugated prodrug of TMP that is currently in phase 3 of clinical trials for early-stage AD patients bearing the *APOE4/4* genotype (NCT04770220)^31,32^. Preliminary *in vitro* and *in silico* studies suggested that both TMP and SPA can lock the Aβ peptides in monomeric conformations that are less prone to oligomerization, thus inhibiting the first step in the pathological pathway of Aβ^33–35^. However, these studies do not provide sufficient insights to fully explain the mechanism of action of these molecules on Aβ. At the moment, it is still unclear whether TMP or its metabolite SPA can exert a stronger biological effect, and this was one of our motivations to carry out this study.

To analyze the effect of TMP and SPA on Aβ and understand how these small molecules may prevent the formation of Aβ oligomers and fibrils, we developed a new computational framework called Comparative Markov State Analysis (CoVAMPnet). The CoVAMPnet framework reveals the impact of a small molecule (in our case, TMP or SPA) on the conformational space and dynamics of an intrinsically disordered biomolecule (in our case, Aβ) in three steps. First, molecular dynamic trajectories are generated using enhanced sampling, and an ensemble of soft MSMs is computed for each system by training VAMPnet neural networks^17^. In particular, we simulated the monomeric Aβ42 peptide in its free form and in the presence of drug candidates TMP or SPA. Second, using our novel alignment method, these ensembles are aligned to identify similar conformational states across the different systems based on a solution to an optimal transport problem. This proved useful in quantifying the similarities and differences in Aβ42 conformations in response to the presence or absence of the small molecules. Finally, our new approach based on analyzing gradients of the trained neural networks is used to elucidate the patterns underlying the learned MSMs and to understand the biophysical relevance of the molecular features, namely the directional inter-residue distances, for the classification into each state. To our knowledge, this is the first time that such a biomolecular relevance analysis has been used to compare and interpret MSMs built by unsupervised machine learning methods and quantify the effects of drug candidates on the conformational space of a disordered protein. Experimental comparison of Aβ42 in its free form and in the presence of TMP or SPA by circular dichroism (CD), Fourier-transform infrared spectroscopy (FTIR), nuclear magnetic resonance (NMR), and fluorometry has further shown the effects of the small molecules on longer time scales, complementing our computational findings.

## MATERIALS AND METHODS

Here, we present only a concise description of the methods used, focusing mainly on the novel methodology. A complete and detailed description is provided in **Supplementary Materials and Methods**.

### Molecular dynamics (MD) simulations

#### System preparation

The structures of tramiprosate (TMP) and 3-sulfopropanoic acid (SPA) were constructed and minimized using Avogadro 2^36^. During the calculation of partial charges, the structures were further optimized by Gaussian 09^37^, and the *antechamber* module of AmberTools 16^38^ was then used to prepare the force field-compatible parameters. The three-dimensional structural data of the Aβ42 peptide was obtained from the RCSB Protein Data Bank^39^ (PDB entry 1Z0Q). It resulted from NMR experiments and contains 30 structures, which were saved separately. The Aβ42 peptide was protonated using PROPKA^40^ at physiological pH 7.4, the small molecules were embedded (when appropriate), the systems solvated, and their topologies built using High-Throughput Molecular Dynamics (HTMD)^41^ in combination with the CHARMM36m^42^ (C36m) force field. We used a stoichiometry of 100 molecules of TMP or SPA per molecule of Aβ42. This ratio approximates the experimental conditions (1000:1) without compromising the computational costs of the simulations.

#### MD simulation protocols

All the systems were equilibrated using HTMD.^41^ The endpoint of the equilibration cycle was taken as a starting point for subsequent MD simulations, either classic or adaptive sampling ones. The simulations employed the same settings as the last step of the equilibration, and their trajectories were saved every 0.1 ns. HTMD was used to perform adaptive sampling of the Aβ42 conformations. Due to the conformational complexity of Aβ42, three protocols (namely A, B, and C) were assessed. Each protocol differed from the others in the starting structure set, the adaptive metric, the number of adaptive epochs and replicas, and the total cumulative MD time (**Supplementary Table 1**). Protocols A and B were only applied to free Aβ42, while protocol C was applied to free Aβ42, Aβ42 + TMP, and Aβ42 + SPA.

Classical MD simulations were also performed using HTMD, where only the structure of the first model of the PDB entry 1Z0Q was used as the starting point. The free Aβ42, Aβ42 + TMP and Aβ42 + SPA systems were prepared and equilibrated as described above. These MDs were performed using only the CHARMM36m force field. Each MD was run in sequential batches of 200 ns each, for a total of 5 μs, and 10 independent replicates were performed for each system.

#### Analyses of properties in combined MD ensembles

In order to analyze the produced MD simulations, their topologies were converted from CHARM to AMBER using ParmEd^43^ when required. Water molecules and ions were filtered out from the resulting MDs, which were then compiled into a simulation list using HTMD. The *cpptraj*^44^ module of AmberTools 16^38^ was used to compute several properties in the combined ensembles: root-mean square deviation (RMSD), radius of gyration (*R*_g_), and linear interaction energy (LIE)^45^ between Aβ42 and TMP or SPA. DSSP 3.0^46^ was used to assign a secondary structure to every residue in every snapshot of the combined trajectories, and the default DSSP seven-letter alphabet was converted to the three main secondary elements (α-helix, β-strand, and coil, see MD analysis section in **Supplementary Materials and Methods**). Accounting for all the residues of each secondary structure type in the peptide for all the analyzed snapshots resulted in the total secondary structure content of the ensemble. Mechanics/generalized Born solvent accessible surface area (MM/GBSA)^47,48^ calculations were performed with the MMPBSA.py.MPI^47^ module of AmberTools 14 to obtain the free energy of the peptide for every frame of the ensemble, from which the peptide intramolecular interactions were derived.

### Comparative Markov State Model Analysis (CoVAMPnet)

This section describes our Comparative Markov State Analysis (CoVAMPnet) of adaptive sampling MD simulations of the free Aβ42, Aβ42 + TMP, and Aβ42 + SPA systems. CoVAMPnet builds on the Variational Approach to Markov Processes by VAMPnet neural networks, followed by two new analyses: (i) alignment of the learned MSM ensembles across different systems based on a solution to an optimal transport problem, and (ii) characterization of the learned states by the inter-residue distances based on the neural network gradients.

#### Learning Markov state models using neural networks

The Variational Approach to Markov Processes (VAMP)^49^ was used to learn Markov state models (MSMs) via unsupervised training of VAMP neural networks (VAMPnets)^16^ with physical constraints^50^. VAMPnet learns a nonlinear function that maps the peptide tertiary structure to a vector of state probabilities. The physical constraints ensure that the learned MSM is reversible and that the elements of the matrix representing the governing Koopman operator^16^ (a linear operator propagating the state probabilities in time) are non-negative. In this work, we used the VAMPnet implementation by Löhr et al.^17^, including the self-normalizing set-up^51^.

The VAMPnet architecture consists of two parallel weight-sharing lobes: one for a frame at time *t* and the other for a frame at time *t* + τ in the same trajectory, where τ is a fixed lag time. Each frame was represented on the input as a vector (780 elements) of the upper triangular part of the peptide inter-residue heavy atom distance matrix without the diagonal and the first two subdiagonals (i.e., without the distances to the first and second neighboring residues). The output nodes in each lobe measure the probabilities of the constructed MSM states for the input frame. The network was trained on pairs of MD simulation frames separated by a selected lag time τ. To obtain the probabilities of the learned states, the frames were run through one of the lobes. For each system, an ensemble of 20 models was built. The pairs of frames were divided into 20 random splits (90% training and 10% validation) and for each split, three VAMPnet models were trained with different initialization and the one with the highest VAMP-E score^49^ was selected for the MSM ensemble. The soft assignment of a frame was defined as the average of its state probabilities across the ensemble, whereas the hard assignment was defined as the state with the highest probability in the soft assignment of the frame. Throughout this work, the soft assignments were used everywhere unless it was necessary to select example frames from a particular state (such as the example structures in **Figure 2A** or the frames representing the states for the columns of the matrix in **Supplementary Figures S22**). Further details on our VAMPnet setup are described in **Supplementary Materials and Methods**.

**Figure 2.**
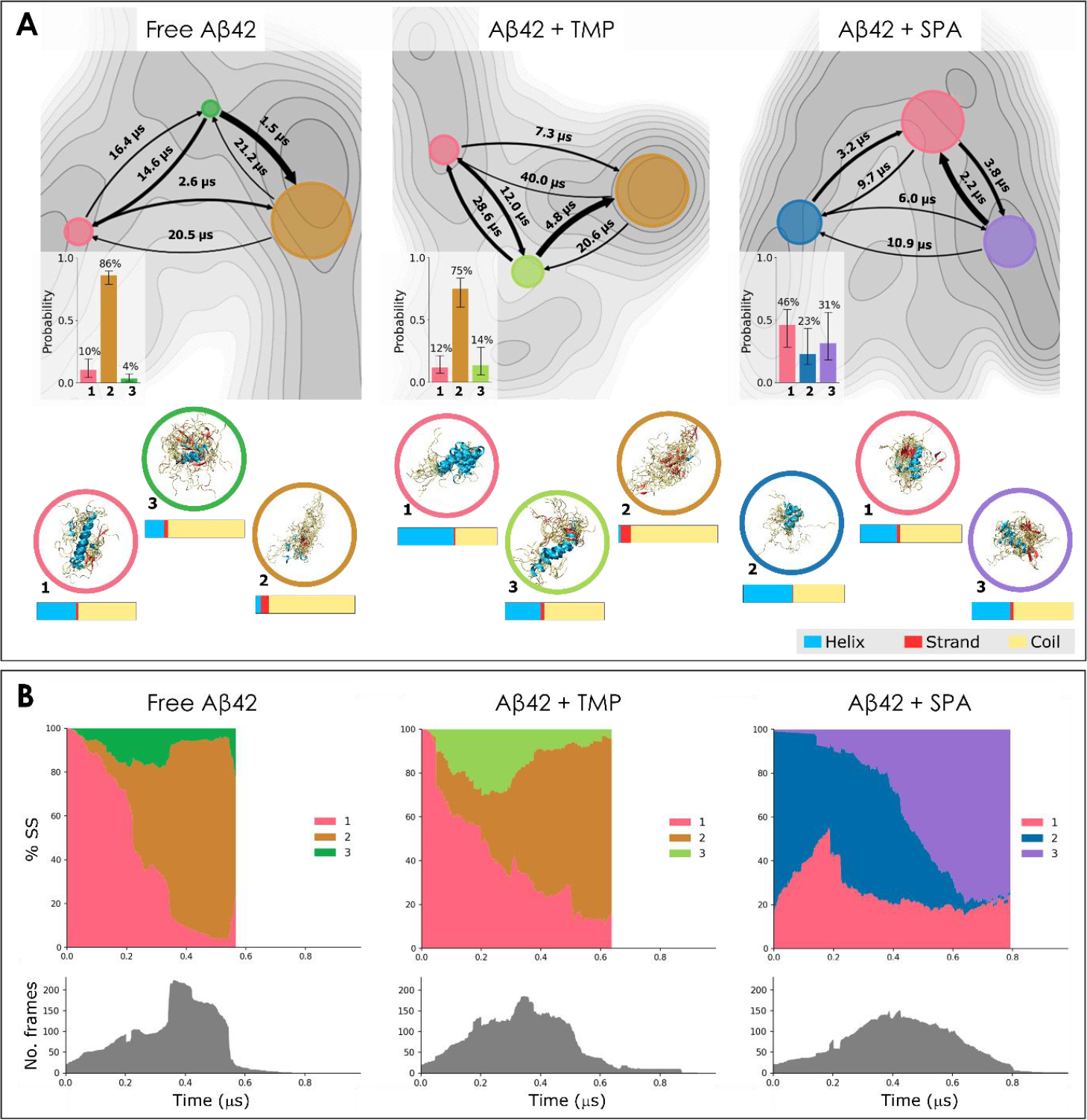
Analysis of conformational states learned using the variational approach to Markov processes on the adaptive simulations and their evolution in time. A) Properties of the states. For each system we report: (i) the free energy surface (FES) projected on the first two tICA dimensions (gray maps), where darker shades correspond to more negative energy regions; (ii) flux diagrams overlapping the FES and projected on the same tICA space, where each state is represented by a colored circle with the area proportional to the state probability, and the arrows indicate the mean first-passage times *T*_*M*_ between the states, with the thickness proportional to the transition probability; (iii) equilibrium distribution of the states (bottom-left corner of FES; the bars represent the 95^th^ percentile of values centered around the median from the ensemble of 20 learned models; see **Supplementary Note 7** for details); (iv) superimposition of twenty representative structures from each state, selected based on the highest assignment probability (below FES, enclosed in colored circles); (v) global mean secondary structure content of each state (below the respective structures). **B) Distribution of the learned states in time (top) and the number of frames available at each time point (bottom)**. The adaptive sampling trajectories were aligned in time and concatenated. The state probability at a given time point was computed as the average soft assignment of all available frames at this time point. From left to right, the state assignments evolve from the beginning to the end of the simulation time. All plots are shown for the free Aβ (left), Aβ + TMP (middle), and Aβ + SPA (right). The states are numbered and color-coded consistently across the entire panel; the same colors across different systems indicate aligned states.

#### Alignment of learned states for comparative analysis

The order of the states on the output of a trained VAMPnet is not well-defined and may thus vary. To construct an MSM from multiple models or compare MSMs of different systems, a correspondence between states across the models had to be established. In this work, we generalized the approach from Löhr et al.^17^ for the alignment of states within a single system to obtain an ensemble of aligned MSMs. Then, we introduced a new method for the alignment of ensembles of MSMs between different systems to compare the systems and further understand the effects of the small molecules on the conformational dynamics of Aβ42.

##### Aligning states within a single system

The states from the 20 models within an ensemble were aligned by a constrained k-means clustering algorithm^52^ using the average inter-residue distance matrices 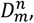, where *n* indexes the models in the ensemble and *m* indexes the states in each model. The cluster centers were initialized by the 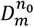 matrices of a randomly selected model *n*_0_ in the ensemble. The clustering iterated in two steps: 1) for each model *n*, its states were sequentially assigned to different clusters in the order of the proximity of the 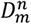 matrix to the closest unassigned cluster center and respecting the constraint that two matrices from the same model cannot be assigned to the same cluster; 2) each cluster center was recomputed as the mean of the 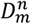 matrices of the corresponding states. These two steps were iterated until the cluster assignment did not change. The states in each model were then re-numbered according to the final assigned cluster. The method by Löhr et al.^17^ is equivalent to performing only one iteration of our method. Our approach is thus less susceptible to incorrect initialization and can lead to a better alignment.

##### Aligning ensembles of Markov state models between different systems

With each system described by an ensemble of *N* mutually aligned MSMs after the single system state alignment (see above), we proposed a novel method for aligning ensembles of MSMs across different systems. In particular, we (i) characterized each state of the given system by a non-parametric distribution over the ensemble, (ii) defined a distance metric to compare such distributions, and finally, (iii) computed an alignment of the ensembles of MSMs between the two systems by solving an optimal matching problem. Details of these steps are given next. The *N* instances of the VAMPnet network learned for a given system *s* output *N* different feature matrices 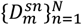 (average inter-residue matrices, see **Supplementary Materials and Methods** for a formal definition of the feature matrix) describing each of the *M* states of the system. Each state *m* was, therefore, characterized by the distribution 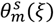 of the features over the different VAMPnet instances as:

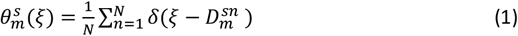

where *δ* is the Dirac delta function defined over the feature space of inter residue distances in which the simulation frames are represented and 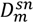 is the inter-residue distance matrix representing state *m* of the learned model *n* for system *s*. 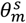 thus represents the state *m* of system *s* with a non-parametric distribution given by the set of Dirac functions centered at the feature matrices 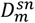 obtained by the instances of the learned ensemble.

To exploit the entire distribution of the features of each state, the distance between two different states was evaluated by comparing their respective distributions. In particular, we employed the Wasserstein distance of two distributions as a distance measure quantifying the cost of aligning two states from different MSMs as:

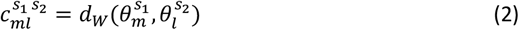

where 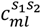 is the cost of aligning state *m* of system *s*_1_ with state *l* of system *s*_2_ and 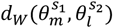 is the Wasserstein-1 distance of the two respective distributions defined as:

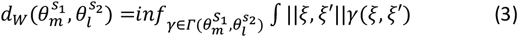

where 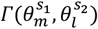 is the set of joint distributions whose left and right marginals are 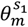 and 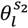 respectively and ‖ξ, ξ′‖ is the Euclidean distance of the two feature vectors ξ, ξ′ distributed according to the joint distribution *γ*(ξ, ξ′). In the case of empirical non-parametric distributions (such as in our case), the problem of Wasserstein-1 distance computation has an equivalent linear program formulation and it was solved using an optimal transport algorithm^53^.

Finally, the alignment of MSM ensembles was formulated as an optimization problem. Without the loss of generality, let us assume that the MSM representing system *s*_1_ does not have more states than the MSM representing system *s*_2_. The problem was defined as:

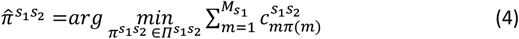

where 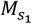 is the number of states of the MSM estimated for system 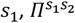 is the set of all bijections from the states of system *s*_1_ into any 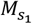 -sized subset of states of system *s*_2_ and the bijection 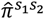 is the optimal mapping of states of system *s*_1_ onto the states of the system *s*_2_. This optimization problem, and thus also the alignment of MSM ensembles, was solved using the Hungarian algorithm^54^.

#### Gradient-based characterization of learned states

The differentiability of the VAMPnet model enables interpretation of the states by investigating the feature importance, which is hard to do using classical Markov state models. This analysis aimed to understand how important the different parts of the protein structure (here represented by the peptide inter-residue distances) are for the definition of different states. While there exist different methods to investigate the importance of features in neural networks^55,56^, they are usually applied to single models for simple tasks, such as the classification of individual images. The challenge of adopting those methods for the current study was in calculating the feature importance for an ensemble of MSMs. We proposed a method to identify which features were important for the classification of the simulation frames into the learned states, building on the gradient-based method proposed for image classification^56^. In our approach, we computed the gradients for each of the models in the MSM ensemble separately and aggregated their results over the ensemble. To this end, the MSMs produced by the models needed to be aligned, which we did by using our state alignment method discussed earlier (see *Aligning states within a single system*). The gradients for individual Markov states were computed as follows:

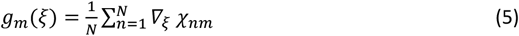

where *g*_*m*_ is a 780-dimensional vector containing the ensemble-averaged gradient of the output probability of state *m* computed with respect to the input features ξ; *N* is the number of models in the ensemble; *∇*_ξ_ is the operator of gradient with respect to the coordinates of the network input features ξ; and χ_*nm*_ represents the output node corresponding to state *m* of *n*^*th*^ VAMPnet model in the ensemble. Here, the 780-dimensional network input vector was obtained by vectorizing the upper triangular inter-residue distance matrix and removing the diagonal and two subdiagonals. The intuition is that the i^th^ entry of vector *g*_*m*_ expresses the change in the probability of the assignment of the given frame of the simulation to state *m* induced by an increase in the distance of the i^th^ pair of residues at the input of the VAMPnet network. The above definition computes the gradient value for an individual frame of the system. To aggregate the gradient value over a representative set of frames from the investigated system, we evaluated the gradient vector 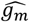 as the average of *g*_*m*_ over 10,000 randomly selected simulation frames ξ. For visualization purposes, we took the 780-dimensional vector of evaluated gradients 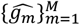 and arranged it back into a 42×42 matrix corresponding to the shape of the inter-residue distance matrix. These gradients evaluated and averaged over randomly selected frames should express the importance of particular residues on average for the classification into a specific state without any particular assumptions about the input frame.

#### Estimation of the free energy landscape

We estimated the free energy landscape of Aβ42 for each of the studied systems, projected on the first 2 time-lagged independent component analysis (tICA) dimensions, by performing Gaussian kernel density estimation on 10% of the simulated frames^17^.

### Experimental validation

Aβ42 in its monomeric form (*N*-methionine-Aβ42, or N-Met-Aβ42) was produced and purified following an adapted version of the protocol by Cohen et al.^57^. Spectroscopic properties of N-Met-Aβ42 alone or in the presence of TMP, SPA, and the membrane-mimicking hexafluoroisopropanol (HFIP) were measured using circular dichroism (CD), Fourier-transformed infrared spectroscopy (FTIR), and nuclear magnetic resonance (NMR). Aggregation kinetics were recorded using thioflavin T (ThT) assays^58^. A 1000-fold molar excess of TMP or SPA with respect to the concentration of N-Met-Aβ42 was used, to replicate the experimental conditions previously reported to exert biological effects from those molecules^33^.

## RESULTS

### Selection of the computational protocol for the simulation of Aβ42

We aimed to query, by molecular dynamics (MD) simulations, the conformational diversity and dynamics of Aβ42 (the most aggregation-prone and the second-most abundant isoform of Aβ^4,5^) and the effect of small molecules on such dynamics. The molar excess of small molecules with respect to Aβ42 was lower in the simulations (100-fold) than in the experiments (1000-fold), but it ensured sufficient interactions with the peptide (see **Supplementary Note 1**). Some of the key parameters to consider in any MD simulation are: (i) the starting conformation, (ii) the MD technique and its length, and (iii) the force field. For the starting conformation, we chose a structure of the full-length peptide obtained from liquid state NMR (PDB IDs 1Z0Q^59^; see **Supplementary Note 2** and **Supplementary Figure S1**). Because of its enhanced ability to sample events occurring in longer time-scales^60–62^, we applied adaptive sampling. This method consists of several MD trajectories simulated in parallel and over multiple consecutive epochs, in an adaptive approach. The MDs from each epoch are iteratively seeded from selected snapshots from previous MDs, according to a predefined criterion. This criterion defines the feature to be sampled (also called *metric* or *collective variable*), and the objective is to maximize the variability of that feature sampled in the overall simulation (in this case, the secondary structures).^63,64^ Based on the literature,^65^ we explored the AMBER ff14SB^66^ (hereafter termed A14SB) and CHARMM36m^42^ (C36m) force fields as the ones likely to provide reasonable ensembles to study Aβ42. Notably, C36m was developed specifically for intrinsically disordered proteins and has already been used with Aβ42^35^. We tested different combinations of parameters in three adaptive sampling protocols, and compared the results to the initial structure, experimental data^59,67^, and previous reports.^8^ The goal was to obtain conformations of Aβ42 diverging from the initial NMR structure (membrane-like environment), and to reach average secondary structure ratios that approximate the experimental ones (aqueous environment). The selected protocol used the C36m force field and A14SB was discarded (protocol C; see **Supplementary Note 2, Supplementary Table S1**, and **Supplementary Figures S2-S4**). The respective MD ensembles seemed to be well converged (**Supplementary Figure S5**).

### Secondary structure content in simulations of free Aβ42 and Aβ42 with ligands

To compare the simulations of Aβ42 alone or in the presence of an excess of TMP and SPA (**Figure 1A**), we first analyzed the global secondary structures content of the peptide in the three systems (**Figure 1B**). In the adaptive simulations of free Aβ42, the peptide showed a larger ratio of coils (77.5%), followed by the α-helices (16.8%) and finally the β-strands (5.7%). In the presence of TMP, the α-helix content of Aβ42 peptide increased by 11.1% to 27.9%, whilst the ratio of β-strands remained unchanged (5.8% vs 5.7%). In the presence of SPA, the differences in the secondary structure were more striking. In this case, the content of α-helices was nearly the same as in the original NMR structure (41.6% vs 42.1%), the ratio of coils was slightly lower (56.1% vs 57.9%), and the β-strands were half of those in free Aβ42 (2.3% vs 5.7%). This striking result suggests a strong effect of SPA in preserving the α-helical structures of Aβ42.

We analyzed the secondary structures in more detail, dissecting the different propensities by the sequence residues (**Figure 1C**). The results showed that Aβ42 could adopt a coiled structure over its entire sequence, with the highest fractions in the N-terminal residues 1-8. Helical structures were most significant for residues 10-20, with α-helical structures near and above 40% and decreasing in further residues. The β-strands were the least frequent element, present at the C-terminal tail of the peptide (residues 30-41) and, to a lesser degree, also around residues 2-8 and 17-20. This is in agreement with Tomaselli et al., who reported the formation of an antiparallel β-sheet made of the two β-strands containing amino acids 18–22 and 37–41.^59^ TMP had little effect on the secondary structure distribution, only slightly increasing the frequency of helical structures in the regions that already had a propensity for it (residues 9-28) and reducing the β-strands in the N-terminal residues 2-8. However, the inclusion of SPA resulted in a substantial reduction of the β-strand content in residues 2-20 and 30-41 and in a significant increase of helical propensity in residues 9-28 and 30-37. Thus, we observed that both studied Aβ modulators (TMP and SPA) could increase the regular structures, specifically protecting the α-helix content of the Aβ42 peptide. The effect was notably stronger with SPA, which also prevented or slowed down the transitions from helices into coils and β-strands.

We further analyzed the different MD ensembles and calculated the radius of gyration (*R*_g_) to assess the compactness of the Aβ42 peptide in the three systems. We found that the free Aβ42 alone had a significantly (with *p value* < 10^−4^ from the *t*-test) broader and more skewed distribution of *R*_g_ (average *R*_g_ = 14.2 ± 4.3 Å) than in the presence of TMP or SPA (*R*_g_ = 13.3 ± 3.1 and 11.8 ± 2.1 Å, respectively; **Figure 1D**). This indicates that the free Aβ42 had a population of extended conformations that was not found in the presence of TMP or SPA. SPA showed a particularly strong effect on shifting Aβ42 towards more compact conformations, compared to the other two systems. Interestingly, Löhr and co-workers recently reported an aggregation inhibitor that presented the opposite effect and stabilized the extended, higher-entropy conformations of Aβ42^68^.

#### Effects of ligands on the evolution of secondary structure elements over time

To understand the evolution of secondary structure elements in the adaptive sampling simulations, we first performed the time-based alignment and concatenation of the MDs (**Supplementary Note 3** and **Supplementary Figure S6**). We computed the evolution of the mean secondary structure content along the continuous simulation time of the aligned and concatenated simulations (**Figure 1E**). We observed that the different secondary structure ratios evolved quickly in the free Aβ42, decreasing for α-helices and increasing for coils and β-strands. In the presence of TMP, those values changed similarly but more slowly, while SPA induced the slowest changes. Classical MDs showed similar trends towards the apparition of coils and strands over time. However, the capacity of the small molecules to preserve helical elements was not as pronounced as in adaptive-sampling MDs (**Supplementary Note 4, Supplementary Figures S7-S9, Supplementary Table S2**). We can speculate that performing longer simulation times might result in a further decrease in the levels of α-helices and an increase of β-strands.

### Conformational analysis of ligand effects using Markov state models

Initially, we tried to construct conventional Markov state models (MSMs) to analyze the adaptive sampling simulations and characterize the conformational states of Aβ42.Different metrics and settings were tested, namely the *RMSD* of the Cα atoms, the *secondary-structure*, the *self-distance* of all Cα atoms, and combinations of those metrics (**Supplementary Materials and Methods**). However, none of these analyses produced reliable models (see example in **Supplementary Figures S10-S12**), so we decided to use the recently published method for MSM construction using artificial neural networks. We further extended that method with new analyses, which proved highly useful for comparing different systems and improving the interpretability of the results.

#### Construction of variational Markov state models

We approached the construction of MSMs with VAMPnet^16^ by testing several lag times (25, 50, 75, and 100 ns) and different numbers of Markov states (2, 3, 4, and 5). Since we are interested in identifying the major differences among the three systems (free Aβ42, Aβ42 + TMP, Aβ42 + SPA), we prioritized the characterization of a few major macrostates rather than many microstates. For this reason, we explored only a relatively small number of states, as done previously by Löhr et al.^17^ According to the implied timescales plots (**Supplementary Figure S13**) and the Chapman-Kolmogorov tests (**Supplementary Figure S14**), we selected τ = 25 ns as the final lag time. By evaluating the impact of the additional states on the change in the frame classification (**Supplementary Figure S15**), together with considering the transition rates for each state, we decided to use the 3-state MSM for all the studied systems. Using the selected parameters, we re-estimated the MSMs for MD simulations generated by protocol C. We first constructed 16 subsets of data by gradual addition of epochs to the training and validation data. From the models, we calculated the exact transition probabilities, mean first-passage times, and transition rates (**Supplementary Figure S16**), as well as the respective structural propensities (**Figure 2A** and **Supplementary Figure S17**). Finally, we verified that additional data did not significantly affect the estimated implied timescales and that the size of our datasets was thus sufficient for VAMPnet training (**Supplementary Figure S18**).

#### Evaluation of the effect of using the soft versus the hard assignment

Interestingly, we found the models to be quite certain about the classification of frames into the learned states, thus diminishing the differences between the hard and the soft assignment. For free Aβ42, Aβ42 + TMP and Aβ42 + SPA we found 99%, 99% and 98% of the frames, respectively, to be classified into one of the states with probability higher than 95%.

#### Alignment of learned states across systems with and without ligands

To automatically detect similar conformational states across different systems and compare the estimated MSMs, we developed and applied a novel alignment method. This method aligns different states, by minimizing the global cost of alignment of MSM ensembles and produces alignment costs for each pair of matched states *T*_*e*_ (see Methods – *Alignment of learned states*). To distinguish truly aligned states from those without a counterpart in the other system, we considered two states as aligned only if their alignment cost was lower than the threshold *T*_*e*_ = 6 (see **Supplementary Note 5**); the threshold was selected empirically by comparing the visualized structures (**Figure 2A**), the secondary structure content, and contact maps (**Supplementary Figure S17**) of the states proposed for mutual alignment. This approach allowed us to find two similar states between free Aβ42 and Aβ42 + TMP (states 1 and 2), and one similar state between free Aβ42 and Aβ42 + SPA (state 1; see **Supplementary Figure S19**).

#### Comparison of learned states across systems with and without ligands

The evolution and kinetics of the constructed MSMs for the studied systems are shown in **Figure 2**, as well as a representative ensemble of structures for every state. The free Aβ42 system (**Figure 2**, left) was characterized by a sparsely populated source state (state 1, pink, 10% equilibrium probability), a dominant sink state (state 2, orange, 86% equilibrium probability), and a meta-stable transition state between them that was the least populated of all (state 3, green, 4% equilibrium probability). The kinetic roles (source and sink) were derived from the transition kinetic rates and the mean first-passage times, and from the secondary structure contents of each state. Hence, the source state (1, pink), with the structural content most resembling the starting NMR structure (ca. 58% coil, 40% α-helices and 2% β-strands), converted fast into the sink state (2, orange; *T*_*M*_ = 2.6 μs), and could be reasonably formed from the transition state (3, green; *T*_*M*_ = 14.6 μs). The sink state was characterized by disorder, with the highest contents of coils and β-strands and the lowest contents of α-helices. The transition state represented a middle point in terms of secondary structure content, and it converted faster into the source or sink states than it was formed. This kinetic ensemble is in good agreement with the results previously described by Löhr et al. for the monomeric Aβ42, namely in terms of microsecond transition times between the states, the presence of one dominant state that was mainly disordered, and the inexistence of long-lived folded states^17^.

According to our alignment method, the Aβ42 + TMP system (**Figure 2**, center) had counterparts in the free Aβ42, namely the disordered sink state (orange) and the helical-rich source state (pink). The equilibrium probability of the sink was slightly reduced (state 2, orange, 75%) and the more helical source was slightly increased (state 1, pink, 12%). A new transition state appeared in this system (lime, 14% equilibrium probability), with intermediate secondary structure propensities and a higher α-helical content compared to the transition state in the free Aβ42. Perhaps for this reason, the cost of their alignment was above the selected threshold (**Supplementary Figure S19**), and the state was thus considered a newly formed state. This was supported by the visualized structures (**Figure 2A**) and the detailed secondary structure and contact maps for the respective states (**Supplementary Figure S17**). Overall, the MSM ensemble for the Aβ42 + TMP system showed higher variability of the equilibrium distribution. Interestingly, the kinetics of this system was rather similar to that of the free Aβ42 but significantly slower, generally with higher transition mean-times. As in the case of the free Aβ42, the formation rates of the disordered sink state 1 were higher than its conversion into the other states.

The simulations of Aβ42 + SPA produced a clearly distinct MSM (**Figure 2**, right), with the equilibrium distribution more uniform than in the other two systems. Furthermore, the confidence intervals of the equilibrium probabilities were even wider, and the free energy landscape appeared more homogeneous, implying that the states in Aβ42 + SPA were less clearly defined compared to the other systems. According to our alignment procedure, only the source state of Aβ42 + SPA (state 1, pink, 46% equilibrium probability) found its counterpart in the free Aβ42 system. The secondary structure content of this state was similar to the corresponding one in the free Aβ42 and the starting NMR structure (61% coil, 36% α-helices, and 3% β-strands). It is noteworthy how the addition of SPA disrupted the kinetic ensemble: the remaining two states differed significantly from those of the free Aβ42, as demonstrated by the high alignment costs (**Supplementary Figure S19**) and the secondary structure contents. Strikingly, in contrast with the previous two systems, the unstructured sink state disappeared as the two new unmatched states with high α-helix contents occurred. This was especially the case of state 2 (blue, 23% equilibrium probability), which contained more α-helices (48.7%) and fewer coils (50.6%) than the initial NMR structure (42.1% and 57.9%, respectively). This state 2 evolved over time into state 3 (purple, 31% equilibrium probability; **Figure 2B**), which had the fastest conversion to the source state, and thus could hardly be considered a “sink” state. All three states interconverted between each other rather quickly, with *T*_*M*_ values in the low microsecond range, suggesting a dynamical meta-stable equilibrium around the source state. All these observations are supported by the study of the time-evolution of the states in the different simulations (**Supplementary Note 6, Supplementary Figure S20**).

We also calculated the radius of gyration (*R*_g_) of the different states (**Supplementary Figure S21**). The free Aβ42 system presented the largest dispersion of *R*_g_ values, with its states showing peaks at higher values, while for Aβ42 + SPA, all the states displayed low *R*_g_ dispersion and peaks at low values (between 10.6-11.0 Å). This observation is in agreement with the *R*_g_ calculations on the global MD ensembles, discussed above, suggesting that the systems differ intrinsically in their degrees of structural order and compactness.

#### Characterization of learned conformational states via network gradients

To better understand the differences between the states in each MSM, we attempted to interpret the molecular features that were determinant to the assignment of each state. For that, we visualized the ensemble-averaged gradients of the state assignment probabilities obtained from the learned neural network models. **Figure 3** shows that the elements near the diagonal were the most important for the classification into the respective states. As our representation does not consider the distances of the residues to their first and second neighbors in the primary sequence, the colored pixels along the empty diagonal in each heatmap correspond to the distances of the residues to their third neighbors in the sequence. Since this roughly corresponds to the length of one turn in an α-helix (ca. 4 residues), the consistently red or blue color of the two subdiagonals closest to the white diagonal to the presence or absence of helices, respectively. This interpretation is also supported by the average secondary structure content per residue and the average contact maps (**Supplementary Figure S17**).

**Figure 3.**
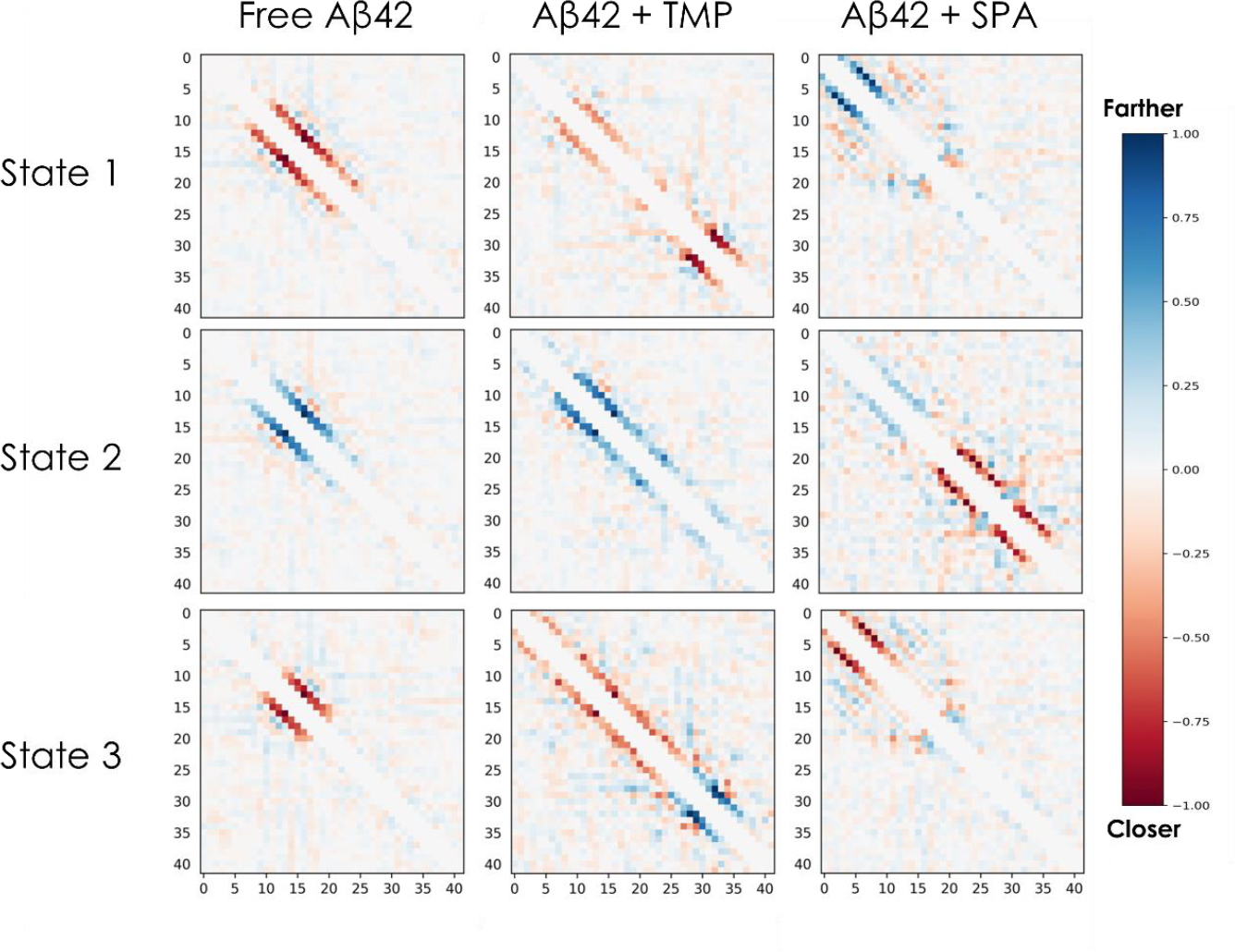
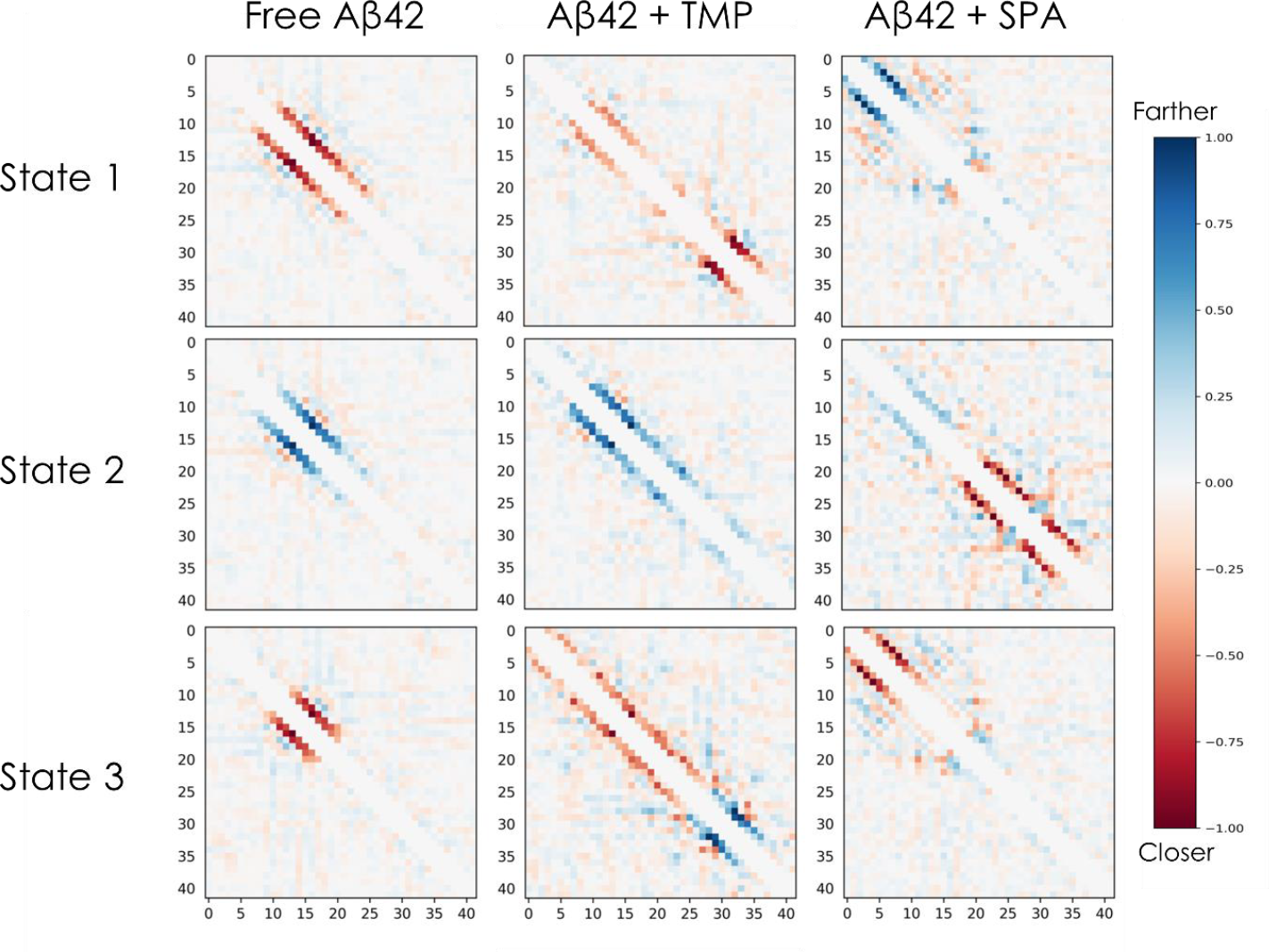
Gradients of the state assignment probabilities of the learned variational Markov state models. Each 42×42 heatmap shows the ensemble-averaged gradients of the model probabilities for the corresponding system and state with respect to the input inter-residue Cα distances. The color indicates how the probability of the particular state would change for an input frame if the distance between the particular pair of residues increased: blue indicates that the probability of the state assignment would increase if the distance between the Cα atoms increased whereas red indicates that the probability would increase if that distance decreased. The presented visualizations correspond to ensemble-averaged gradients evaluated and aggregated over 10,000 randomly selected simulation frames. Columns: MSMs for the free Aβ42 (left), Aβ42 + TMP (middle), and Aβ42 + SPA (right) systems. Rows: states 1 (top), 2 (middle), and 3 (bottom) of each model.

For the free Aβ42 system, the peptide residues around positions 10-25 seem to be crucial for the state classification. The results in the free Aβ42 state 1 heatmap imply that if the red colored residues in this region got closer to their 3^rd^ and 4^th^ sequence neighbors in a particular snapshot, the probability of classifying that snapshot into state 1 (source state) would increase. This means that state 1 prefers a helical conformation in this region. On the contrary, the “state 2” heatmap shows that the probability of classification into state 2 would increase if the blue-colored residues in this region got farther from their 3^rd^ and 4^th^ sequence neighbors, i.e., state 2 (sink state) prefers disorder in this region. The classification into state 3 relies on the same region (residues 10-25) but is split into two parts: residues 13-19 (red) and the rest (gray). This implies that state 3 (transition state) prefers a short helix only in residues 13-19.

For the Aβ42 + TMP system, the corresponding heatmaps show that the presence (red) or lack (blue) of a helix at positions 29-36 are important for distinguishing between states 1 and 3, respectively, while state 2 can be discriminated based on the lack of a helix at positions 10-25. For Aβ42 + SPA, the lack (blue) or presence (red) of a helix at positions 3-12 is relevant for discriminating states 1 and 3, respectively. State 2 differs by the presence of two helices at positions 20-27 and 30-35 (red) as well as by long distances between residues in positions 10-17 (blue pattern).

The states can be compared in more detail by evaluating the gradients on sets of state-specific frames (**Supplementary Figure S22**). Conversely, the gradient matrices can also be aggregated by residue into simpler but still very informative plots (**Supplementary Figure S23**). These can help to readily assess the most influential regions defining the states, compare different systems, and potentially cross-validate the results with other residue-based analyses, e.g., from experimental data (see below).

### Molecular interactions

#### Ligand-peptide interactions

The interactions of TMP and SPA with Aβ42 were assessed by the linear interaction energy (LIE)^45^, and computed for all the 100 ligand molecules with each peptide residue during the adaptive sampling simulations. For this purpose, all the snapshots in the simulations were used. The electrostatic component (Δ*G*_bind_^elec^) dominated the interactions formed by Aβ42 with both TMP and SPA, overshadowing the van der Waals component (**Supplementary Figure S24**). Those interactions were, on average, much stronger with the charged residues (**Figure 4** and **Supplementary Figure S25**). This was expected, considering that both TMP and SPA bear two charges at physiological pH, separated by only a short alkyl chain (a positive and a negative charge in TMP, and two negative charges in SPA). SPA showed both attractive and repulsive interactions (respectively, positive and negative Δ*G*_bind_ ^elec^; **Figure 4**), TMP showed mostly favorable interactions (negative Δ*G*_bind_ ^elec^; **Supplementary Figure S25**). The absolute mean interaction energies were also higher with SPA (from -114 to 71 kcal/mol) than with TMP (from -50 to 0 kcal/mol). Moreover, the interactions were highly variable due to the rapid exchange of the TMP and SPA molecules, which formed unspecific short-lived interactions with Aβ42. This explains the large populations of snapshots with a lower range of interaction energies and the smaller populations of snapshots with strong interactions with the charged residues.

**Figure 4.**
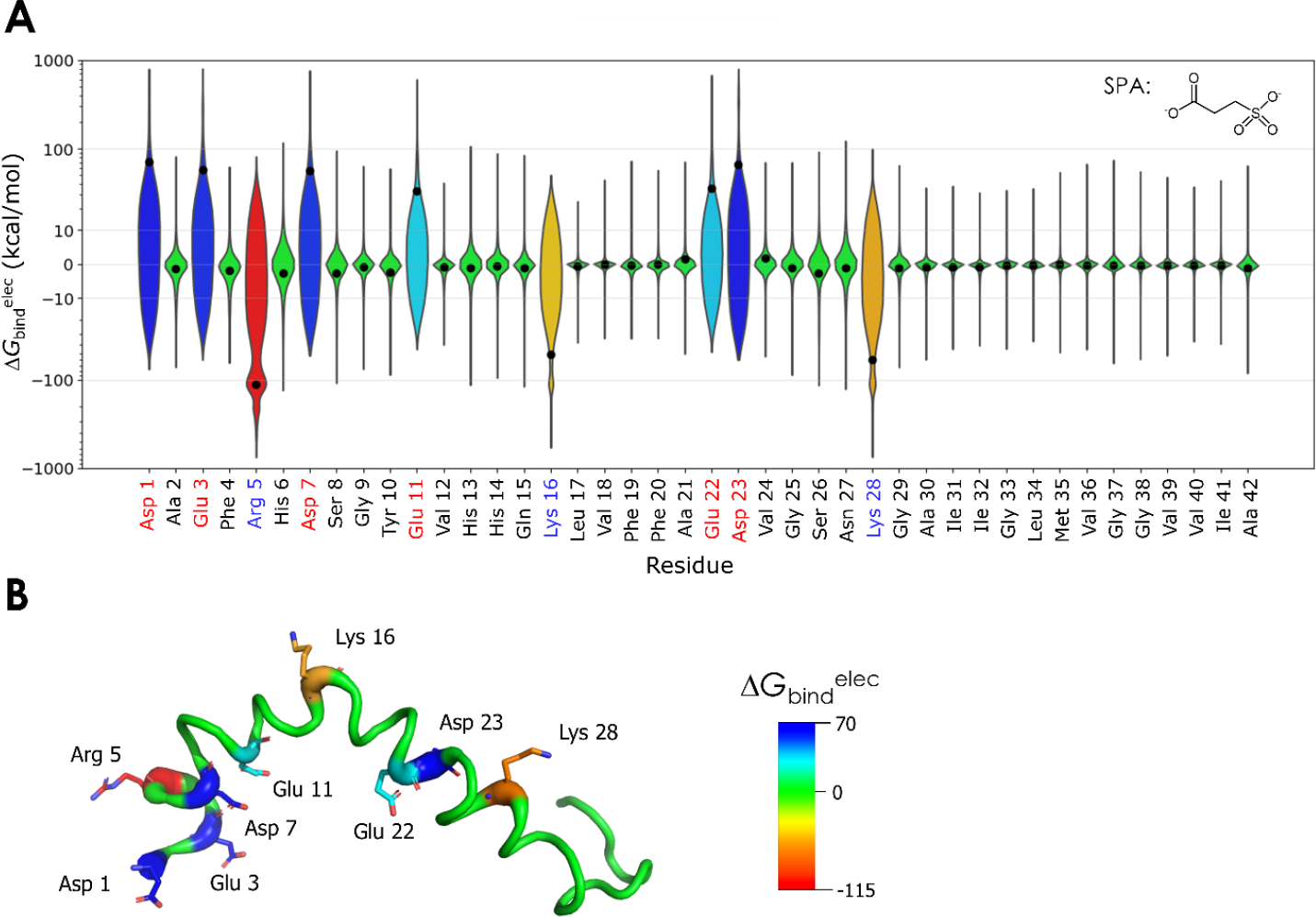
Interactions of SPA with Aβ42 studied by molecular dynamics. A) Violin plot of the binding energy of SPA with each residue of Aβ42. The electrostatic component (Δ*G*_bind_ ^elec^) was calculated for all the 100 molecules in every snapshot of the adaptive simulation of Aβ42 + SPA. The plot shows the distribution of the energy values; the black dots show the mean values; the y-axis uses a quasi-logarithmic scale based on the inverse hyperbolic sine to highlight the higher absolute values. The residue labels are colored by charge: black for neutral, blue for positive, and red for negative. The chemical structure of SPA is shown in the upper-right corner. **B) Structure of Aβ42 with the main interacting residues**. Aβ42 is shown as the putty cartoon and the main interacting residues are represented by sticks (structure from PDB ID 1Z0Q). The colors reflect the mean Δ*G*_bind_ ^elec^ (in kcal/mol) and range from the most positive (blue) to the most negative (red) values obtained for SPA.

Although TMP and SPA have quite similar structures, the global effects of SPA on Aβ42 were more striking than those of TMP. This is probably due to the fact that SPA has a double negative charge, which reverses the charge of positive groups it interacts with. Conversely, TMP is zwitterionic (with a positive and a negative charge) and thus preserves the charge around the interacting residues. A comprehensive comparison of the properties of TMP and SPA and their effects on the simulations of Aβ42 is presented in **Supplementary Table S3**.

#### Intramolecular interactions of Aβ42

The interactions within the Aβ42 peptide were calculated using the molecular mechanics/generalized Born solvent accessible surface area (MM/GBSA) method^47,48^. Interestingly, the electrostatic energy prevailed over the van der Waals, but the polar solvation energy outweighed all the other contributions to the internal free energy of Aβ42 (**Supplementary Table S4** and **Supplementary Note 8**). The peptide was more stable (lower mean total free energy) in the presence of TMP or SPA than alone in solution. This stabilization was mainly due to the solvation energy, which indicates a higher exposure of polar residues to the solvent than the free Aβ42. This effect is concomitant with an increase of the internal hydrophobic contacts in the presence of TMP or SPA, which is consistent with an increase of the compactness of the peptide, according to the *R*_g_ values reported above (**Figure 1D**). Intramolecular salt bridges E22-K28 and D23-K28 have been reported to be important for the conformational transition, oligomerization, and toxicity of Aβ42^69,70^. Analysis of the three ensembles showed that these salt bridges occurred considerably less often in the presence of TMP than in the free Aβ42, and even less with SPA (**Supplementary Figure S26**). This suggests a lower propensity of Aβ42 to form oligomers in the presence of those small molecules. Due to their charged moieties, TMP and SPA induce electrostatic dispersion on the residues involved in the salt bridges, thus weakening those interactions (**Supplementary Table S3**). Similar observations have previously been reported for apolipoprotein E (ApoE) interacting with SPA^71^.

### Experimental validation

To validate our computational findings described above, we experimentally characterized the conformations of N-methionine-Aβ42 (N-Met-Aβ42) alone and in the presence of TMP and SPA. The presence of N-terminal methionine was necessary for the Aβ42 recombinant expression and does not influence its aggregation behavior. This is demonstrated by the routine use of N-Met-Aβ42 aggregation studies^72,73^. Circular dichroism (CD) of N-Met-Aβ42 in aqueous buffer revealed that the peptide was mainly disordered (68% of coils, 29% of β-strands and 3% of α-helices; **Figure 5A** and **Supplementary Figure S27A**). To replicate the NMR structure obtained in 20% (v/v) of hexafluoroisopropanol (HFIP), used herein as the starting conformation for the computational analysis, we titrated the N-Met-Aβ42 with increasing concentrations of HFIP. At 20% HFIP, the secondary structure content of N-Met-Aβ42 was heavily changed in favor of the α-helices, in agreement with the literature^59^ (**Supplementary Figure S28**). We repeated the titrations in the presence of a 1000-fold excess of TMP or SPA. In all cases, no major changes in the CD spectra were induced by the small molecules during the titrations (**Supplementary Figures S27A** and **S28**). N-Met-Aβ42 remained mostly disordered at 0% HFIP and had almost similar helical and strand content at 20% HFIP, independently of the presence of TMP or SPA. This is not in agreement with the computational results, which predicted a significant increase of the helical content of Aβ42 with the small molecules, especially with SPA.

**Figure 5.**
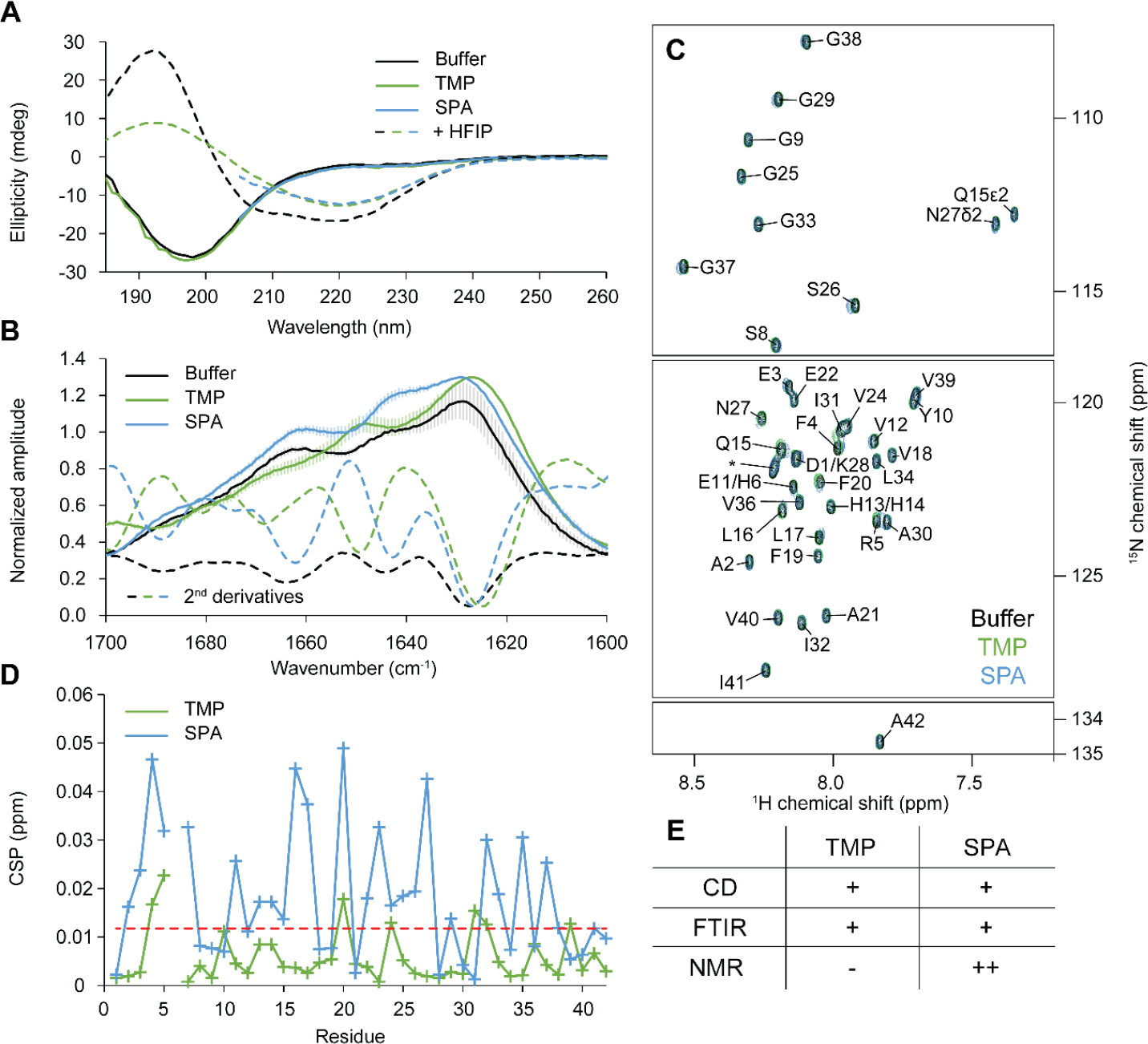
Experimental validation of computational data using biophysical techniques. A) Circular dichroism spectra of Aβ42. N-Met-Aβ42 (37 μM) was studied in the absence (black) or presence of a 1000-fold excess of TMP (green), SPA (blue) or 20% HFIP (dashed curves). The curves for SPA were trimmed below 205 nm to remove the signal from SPA. **B) FTIR spectra of Aβ42**. N-Met-Aβ42 (60 μM) was studied in the absence (black) or presence of a 1000-fold excess of TMP (green) or SPA (blue). The bars represent the standard deviations from successive acquisitions. The second derivatives are drawn as dashed curves. Offset was shifted to improve readability. **C) NMR analysis of Aβ42**. ^1^H-^15^N HMQC NMR spectra of ^15^N-labeled N-Met-Aβ42 were determined alone (black, 69 μM) and in the presence of a 1000-fold excess of TMP (green, 58 μM) or SPA (blue, 55 μM). Assignment is given for free N-Met-Aβ42 (black); the assignment of His6 was ambiguous, thus no CSP was calculated for this residue. **D) NMR chemical shift perturbation (CSP) of Aβ42**. N-Met-Aβ42 in the presence of a 1000-fold excess of TMP (green), or SPA (blue) with respect to the free N-Met-Aβ42. The red dashed line represents the threshold for significance, taken as the standard deviation of all CSPs. **E) Summary of the effects of small molecules on Aβ42 conformations studied by three different biophysical techniques**. - indicates that no significant effect was detected, + indicates a mild effect, and ++ a stronger effect.

To determine whether the molecules induced subtle changes in secondary structure that are below the resolution limit of CD spectroscopy, we analyzed the N-Met-Aβ42 in buffer and in the presence of the molecules using Fourier-transformed infrared spectroscopy (FTIR). Based on the secondary structure deconvolution of the amide I bands^74^, the FTIR spectra of free N-Met-Aβ42 and N-Met-Aβ42 + SPA showed fingerprints from both helical (peak at around 1660 cm^-1^) and strand contributions (peak below 1650 cm^-1^) (**Figure 5B** and **Supplementary Figure S27B** and **S29**). At 1000-fold excess of TMP, a shift of the peak wavenumbers was observed (**Figure 5B**). The spectrum for N-Met-Aβ42 + TMP had one peak centered around 1650 cm^-1^ instead of 1660 cm^-1^, which might suggest more random conformation (coils) of N-Met-Aβ42 in the presence of TMP compared to the free peptide. Nonetheless, the large overlap of the two peaks casts doubts on such interpretations. Further remarks on differences in secondary structure propensities are discussed in **Supplementary Note 9**.

To gain deeper insights into conformational changes of N-Met-Aβ42 upon the addition of the small molecules, we employed nuclear magnetic resonance (NMR). The ^1^H-^15^N HMQC spectral fingerprint of N-Met-Aβ42 revealed a narrow distribution in δ(^1^H) of the backbone amides (from 7.5 ppm to 8.5 ppm), a characteristic of intrinsically disordered peptides (**Figure 5C** and **Supplementary Figure S30**). Using ^1^H-^1^H NOESY and ^1^H-^1^H-^15^N NOESY-HMQC spectra, we assigned the spectral fingerprint and computed the secondary structure propensities using chemical shift indexing^75,76^. This method is based on the published NMR statistics, where each residue is expected to have a chemical shift within a certain region of the spectrum that is a function of its local secondary structure. The resulting global secondary structure propensity was much higher in α-helices than what was previously obtained by CD (29.6% vs 3%, respectively; **Supplementary Figs. S27A** and **S27C**). The secondary structure probabilities of the different residues showed the highest β-strand propensity for the C-terminal tail, and the highest helical propensity of residues 15-25 (**Supplementary Figure S27D**). This is in agreement with the results from our simulations for the free Aβ42 (**Figure 1C**). We titrated N-Met-Aβ42 with increasing concentrations of TMP or SPA, up to a 1000-fold excess (**Figure 5C** and **Supplementary Figure S30**) and measured the chemical shift perturbation (CSP) in the ^1^H-^15^N HMQC spectral fingerprint (**Figure 5D**). The threshold for the CSP significance was taken as the standard deviation of all chemical shifts^77^. Only small CSPs were observed when adding SPA, which were not sufficient to indicate a shift in the global secondary structure (**Supplementary Figure S27C**). This is not unprecedented, as others have also reported minimal changes in the NMR spectrum of Aβ42 upon the binding of small molecules^78^. CSP was observed across most of the peptide sequence in the presence of SPA, namely in the regions 2-7, 11-17, 20, 22-27 and 32-37. Strikingly, these regions correspond to peptide ranges that emerged in the gradient-based analysis of learned conformational states (namely regions 3-12, 10-17, 20-27, 30-35; **Figure 3** and **Supplementary Figure S23**). In the presence of SPA, close distances (structural order) between residues 2-7 are characteristic of the transition between states 1 (pink in **Figure 2A**) to 3 (purple in **Figure 2A**). Similarly, close distances in the residues 22-27 and 32-37 are characteristic hallmarks of state 2 (blue), which is also determined by long distances (disorder) in the range 11-17. It is noteworthy that states 2 and 3 in this system are distinctively different from the other two systems. Thus, gradient-based analysis of learned states was able to pinpoint similar conformational events as the ones captured by NMR. Moreover, region 22-27 is neighboring the salt bridges between 22-28 and 23-28, which are relevant to the conformational transition, oligomerization, and toxicity of Aβ42^69,70^, as pinpointed in the *Intramolecular interactions of Aβ42* section.

Finally, we assessed the fibril formation of N-Met-Aβ42 using the well-known thioflavin T (ThT) fluorometric assay with and without the small molecules. We found that neither TMP nor SPA seemed to significantly reduce the N-Met-Aβ42 fibril formation rates, as observed by other groups^79^. This is in contrast with HFIP, which is a known solubilizing agent of Aβ42 and a crude membrane mimetic^59^ (**Supplementary Figure S31**). In fact, a change in the CD spectrum was observed in the presence of HFIP and either TMP or SPA (**Figures 5A, 5E** and **Supplementary Figure S28**).

## DISCUSSION

Alzheimer’s disease drug candidate TMP and its metabolite SPA are thought to modify the conformational dynamics of the Aβ42 peptide and decrease its propensity to form toxic oligomers^33,34^. The conformational diversity of Aβ42 has been previously explored by exploiting the variational approach to Markov processes in VAMPnets^16^ to construct Markov state models (MSMs), to better capture the slowest processes in MD simulations^17,68^. However, the exact mechanism of action of TMP, and particularly SPA, on Aβ42 was still unclear. To fill this gap, we first applied the variational approach to Markov processes on adaptive sampling MD simulations using VAMPnets^17^, and then ran our newly developed Comparative Markov State Analysis (CoVAMPnet) pipeline to: (1) align the learned conformational states across ensembles of different MSMs, and (2) based on the learned VAMPnet gradients, to characterize these states by the inter-residue distances. The CoVAMPnet alignment method proved a powerful approach to: (i) quantitatively compare the different conformational states of Aβ42, (ii) identify which states were preserved across different systems, and (iii) identify which states were unique. The CoVAMPnet gradient-based characterization of the learned ensembles of Markov states utilizes the end-to-end differentiability of the neural network-based MSMs, i.e., a property that the conventional methods for MSM estimation lack. The analysis of gradients allowed us to reason, at the molecular level, which residues are responsible for the assignment to a specific state obtained from the variational Markov state analysis. We expect these newly developed methods, i.e., (i) the alignment of ensembles of variational Markov state models across different systems, and (ii) the gradient-based characterization of learned states, to become valuable for studying the impact of small molecules on the conformational dynamics of intrinsically disordered proteins and peptides^80,81^.

The newly developed analyses were applied to MD simulations of Aβ42. It is know that the sampling protocol (namely the force field, the length of the simulations, the adaptive metrics, and the simulation method) can highly influence the global results^82,83^. This is largely due to the intrinsically disordered nature of the Aβ42 peptide, which has a rather shallow energy landscape with many energy minima separated by small energy barriers^6,80^. For this reason, the conformational sampling of Aβ42 remains a challenge^82,83^. Starting from a helix-rich Aβ42 structure, biased towards the conformation in the membrane environment^59,84^ (PDB ID 1Z0Q), we identified the most suitable adaptive protocol to simulate Aβ42, according to the secondary structure contents expected in aqueous phase (dominated by coils and β-strands). In this way, we sampled the conformations and transitions occurring immediately after the release of Aβ42 from the transmembrane region to the extracellular fluid. After approximately 64 μs of adaptive MDs, the free Aβ42 diverged substantially from the initial structure, increasing the total amount of random coils and β-strands (as expected) while decreasing the ratio of α-helices, and became closer to experimental values and previous reports^59,67,85^. We identified two regions of Aβ42 that were more prone to form β-strands (mainly residues 2-8, 17-20 and 30-41). The MSMs learned from the variational Markov state analysis revealed that the most populated state of Aβ42 is highly disordered and contains some β-strands. This state is in equilibrium with two other states with higher contents of α-helices, but still bearing mainly coils. These results are in good agreement with recent reports by Löhr et al., obtained from much longer simulation times (315 μs)^17^.

The presence of TMP and SPA shifted Aβ42 towards more structured conformations (less coils and higher content of α-helices) and reduced the propensity of regions 2-8, 17-20 and 30-41 to form β-strands. This behavior is similar to what has previously been reported for some aggregation inhibitors^86–88^, and is in contrast with some others^68^. The variational Markov state analysis showed that TMP and SPA induced a change in the equilibrium distribution and interconversion rates of the Aβ42 conformational states. SPA exerted a much stronger effect, stabilizing new conformational states that were richer in α-helices than in the other systems. Since β-strand structures lead to the formation of β-sheets, the precursors that prompt the oligomerization and fibrillation of Aβ^2,4,5^, these results suggest the potential of TMP and SPA to inhibit or delay both processes. This can be particularly relevant if we consider previous studies suggesting that oligomers may start by the formation of β-hairpins made of β-strands of residues 16-24 and 28-35^89^, and that α-helices in regions 10-21^85^ or 17-21^90^ may prevent the formation of higher oligomers and aggregation. While Aβ42 is preserved in its monomeric form, it should not be harmful until it is cleared from the brain, namely through the binding to apolipoprotein E (ApoE)^91–93^. Our simulations suggest that TMP and SPA may affect the conformational equilibrium of Aβ42 in the brain and prolong its monomeric soluble state, thus allowing to extend the effective time of the clearance mechanisms. Due to their charged terminal moieties, both TMP and SPA formed mainly electrostatic interactions with the charged residues of Aβ42. These interactions were non-specific and short-lived, but they promoted the exposure of polar residues (similar to a “solvation” effect), but they promoted the exposure of polar residues (similar to a “solvation” effect), induced Aβ42 to be more compact, and weakened intramolecular electrostatic interactions (as previously observed^71^). Importantly, some of the intramolecular salt bridges (E22-K28, D23-K28) considered to promote the formation of β-sheets, aggregation and neurotoxicity of Aβ42^69,70,89^ were disrupted by the presence of those small molecules. The difference between TMP and SPA in terms of charge distribution (zwitterionic and doubly negative, respectively) is likely the main factor responsible for the overall stronger effects of SPA (see **Supplementary Table S3**). The reasons for the stronger stabilization of α-helices by SPA are not clear. However, it may be due to competition of the densely charged ligand with the water molecules, which may lead to preventing their destabilizing action on the peptide, as previously described for a series of ions at higher concentrations^94^.

The CoVAMPnet algorithm developed for identification of structural features in the learned variational MSMs based on network gradients proved useful. We were able to identify the peptide regions with preferential order or disorder in the different states and pinpoint major differences across the different systems. Remarkably, this analysis showed good agreement with the CSPs in the NMR spectra, correctly predicting the peptide regions most affected by the presence of SPA. These computational findings were in agreement with previous studies involving Aβ, TMP, and SPA, namely: (i) the unstructured nature of the peptide, (ii) shift of the Aβ42 conformations by those ligands towards more compact structures, (iii) reduction of the β-strand propensity, and (iv) non-specific interactions with charged residues^33,35,95^. Reports also have shown that both small molecules can interact with the soluble Aβ40 or Aβ42, change their dominant conformation, inhibit the formation of oligomers and fibrils, decrease the Aβ-induced neuronal cell death^25,33,34^, and have protective effects *in vivo*^30^.

We applied several experimental biophysical techniques to validate the computational results described above. Although the experimental outcomes showed only a mild influence of both TMP and SPA on N-Met-Aβ42, several relevant effects were observed (**Figure 5E**). FTIR revealed slight changes in secondary structure upon the addition of TMP, suggesting higher coil conformation propensity for the peptide. On the other hand, NMR showed a stronger impact of SPA on the ^1^H-^15^N NMR spectral fingerprint of N-Met-Aβ42, indicating either direct ligand-peptide interactions, subtle changes in secondary structure, or both. Strikingly, these perturbations were observed in the same peptide regions highlighted by our network gradient analysis. TMP did not produce significant CSPs. Altogether, these results suggest a stronger effect of SPA on Aβ42 than TMP. Yet, the fibril formation kinetics of N-Met-Aβ42 seemed unaffected by TMP or SPA.

The experimental results corroborated several computational findings: (i) the intrinsically disordered Aβ42 interacts with TMP or SPA molecules through many weak interactions, (ii) these interactions induce conformational changes on the peptide, (iii) SPA has stronger influence on Aβ42 than TMP, and (iv) the regions affected could be predicted by the gradient analysis of the learned state probabilities. On the other hand, not all the predictions from our molecular modeling were confirmed experimentally: (i) Aβ42 showed higher β-strand content compared to the computational results, and (ii) TMP and SPA did not change significantly the global secondary structure propensities of Aβ42 and did not prevent fibril formation. The differences in the time scales sampled by the simulations (microseconds) and the experiments (minutes/hours) and the peptide concentration effects may have contributed to this discrepancy. Moreover, the membrane mimetic HFIP modulated the impact of TMP and SPA on N-Met-Aβ42, which may deserve further investigation. An extended discussion of these phenomena is provided in **Supplementary Note 10**. In further works, the development of *specific binders* able to stabilize α-helices in the regions of Aβ42 mentioned above could be a better approach for designing drugs targeting the neurotoxic oligomerization of Aβ. The interaction of TMP and SPA with other proteins participating in the amyloid cascade^96^, which has been demonstrated in the case of ApoE (especially ApoE4)^71,97^, should also be considered and evaluated in future studies. Particularly, we have recently shown the strong impact of TMP and SPA on ApoE4, shifting its structure and properties towards those of ApoE3 and significantly reducing its aggregation.^71^ The observation of significant effects of TMP and SPA on ApoE4, but weaker ones on Aβ, is also important in the context of a recently published paper reporting the existence of five subtypes of AD^98^. All subtypes showed a higher prevalence of the APOE e4 genotype, while only selected ones are characterized by modified levels of Aβ. In the future, it will be interesting to relate this information to the data collected within phase 3 of clinical trials, which will reveal the efficacy of TMP on the different subtypes.

In summary, in this work, we introduced CoVAMPnet to compare and interpret learned MSMs across different systems. CoVAMPnet is composed of two methods: (i) the alignment of Markov state models, and (ii) characterization of learned conformational states based on network gradients. The CoVAMPnet approach can be applied to study and compare any related molecular systems and extract valuable information. It can be especially useful to study the impact of small molecules on intrinsically disordered proteins and peptides, whose quantitative analysis can be extremely difficult. Furthermore, we applied CoVAMPnet to study molecular effects of potential anti-Alzheimer’s drugs on hallmark peptide Aβ42. Our computational results suggested that TMP, and particularly SPA, in short dynamic time windows can stabilize structured helical conformations of Aβ42, potentially preventing its oligomerization. *In vitro* validation confirmed the stronger impact of SPA on Aβ42 and the peptide regions affected by this molecule. However, in long time ranges the global secondary structure was not significantly modified, neither was the Aβ42 aggregation propensity under the experimental conditions. This suggests the potential existence of additional mechanisms, such as the suppression of ApoE4 aggregation^82^, contributing to the mode of action and the clinical effects of TMP/SPA in AD besides the conformational shift of Aβ42.

## Supporting information

Supplementary materials and methods, supplementary notes, supplementary figures, supplementary tables, and supplementary references

## Data availability

The sampled stripped trajectories and intermediate data, including the trained neural network weights, are available from https://data.ciirc.cvut.cz/public/projects/2023CoVAMPnet/.

## Code availability

The code and example data are available from https://github.com/KoubaPetr/CoVAMPnet.

## Supporting Information

Detailed materials and methods, complementary to the concise descriptions in this main text. Supplementary discussions (Notes 1-10). Supplementary figures: structure of Aβ42 (Figure S1), comparison of the Amber ff14SB and CHARMM36m force fields (Figure S2-S3), comparison of different adaptive sampling protocols (Figure S4-S5), temporal alignment and concatenation of the adaptive sampling and classical MDs (Figure S6-S9), conventional Markov state model analysis (Figure S10-S12), variational Markov state analysis using VAMPnets (Figure S13-S18), comparative Markov state model analysis (Figure S19), time-based evolution of the states (Figure S20), radius of gyration by state (Figure S21), characterization of learned conformational states via network gradients (Figure S22-S23), interactions of Aβ42 with the small molecules (Figure S24-S26), experimental validation (Figure S27-S31). Supplementary tables: comparison of computational protocols for the simulation of Aβ42 (Table S1), analysis of classical MDs (Table S2), summary of the effects of small molecules (Table S3) (PDF).

## ACKNOWLEDGEMENTS

This work was supported by the Ministry of Education, Youth and Sports of the Czech Republic (grants ESFRI RECETOX RI LM2023069, e-INFRA CZ LM2018140, ESFRI ELIXIR CZ LM2023055, TEAMING CZ CZ.02.1.01/0.0/0.0/17_043/0009632), the European Regional Development Fund under the project IMPACT (reg. no. CZ.02.1.01/0.0/0.0/15 003/0000468), the National Institute for Neurology Research (EXCELES Neuro nr. LX22NPO5107 MEYS): Financed by European Union – Next Generation EU, and the European Union (SinFonia 814418, TEAMING 857560 and ERC FRONTIER 101097822). Views and opinions expressed are those of the author(s) only and do not necessarily reflect those of the European Union or the European Research Council. Neither the European Union nor the granting authority can be held responsible for them. Petr Kouba is a holder of the Brno Ph.D. Talent scholarship funded by the Brno City Municipality and the JCMM.

