## Supplementary materials and methods, supplementary notes, supplementary figures, supplementary tables, and supplementary references for "CoVAMPnet: Comparative Markov State Analysis for Studying Effects of Drug Candidates on Disordered Biomolecules"

### SUPPLEMENTARY INFORMATION

### SUPPLEMENTARY MATERIALS AND METHODS

#### Molecular dynamics (MD) simulations

##### *System preparation*

The structures of tramiprosate (TMP) and 3-sulfopropanoic acid (SPA) were constructed and minimized using Avogadro 2[1]. On the one hand, TMP was built with the protonated amine ($NH_3^+$) and deprotonated sulfonate ($SO_3^-$), which is expected at pH 7.4, with a neutral overall charge. On the other hand, SPA at pH 7.4 is deprotonated on both the sulfonate ($SO_3^-$) and carboxylate ($COO^-$) groups, with a net charge of -2. The minimization step was performed by the Auto Optimization Tool of Avogadro, using the UFF force field[2] with steepest descent algorithm. The resulting structure was then submitted to further optimization and calculation of their partial atomic charges using Gaussian 09[3], with the Hartree-Fock method and 6-31G(d) basis set in vacuum. The *antechamber* module of AmberTools 16[4] was used to extract the RESP charges for the ligands from the Gaussian output files, which were compiled in the PREPI parameters file using the atom types of the AMBER force field. To prepare the PAR and RTF parameter files compatible with the CHARMM force field we used the SwissParam webserver[5]. The three-dimensional structural data of the Aβ42 peptide was obtained from the RCSB Protein Data Bank[6] (PDB entry 1Z0Q). The entry contains 30 structures derived from NMR experiments, and all these structures were extracted, saved separately, and used as starting points for independent adaptive sampling simulations.

The following steps were performed with the High Throughput Molecular Dynamics (HTMD)[7] scripts. Each peptide structure was protonated with PROPKA 2.0 at pH 7.4 [8]. For the systems with ligands, during the preparation protocol of HTMD[7], 100 molecules of TMP or SPA were randomly placed within a radius of 10 Å away from the protein, and at least 5 Å away from each other. A molar stoichiometry of 100 TMP or SPA molecules per molecule of Aβ42 was selected to approximate the experimental excess (1000:1) without compromising the computational costs of the simulations. The concentration of ligand was calculated after the equilibration dynamics, based on the cell volume, and corresponded to approximately 250 mM.

The systems (free Aβ42 or Aβ42 + ligands) were solvated in a cubic water box of TIP3P[9] water molecules with the edges at least 20 Å away from the protein (for free Aβ42) or 5 Å away from the ligands (for Aβ42 + ligands), by the *solvate* module of HTMD. $Cl^-$ and $Na^+$ ions were added to neutralize the charge of the protein and get a final salt concentration of 0.1 M. The topology of the system was built, using the *amberbuild* module of HTMD, with the AMBER ff14SB[10] force field (here termed A14SB) and the previously compiled PREPI parameters file for the ligands. Alternatively, we used the *charmmbuild* module of HTMD, with the modified CHARMM36m[11] force field (here termed C36m) and the parameters for the modified mTIP3P[9] solvent model, with the respective parameters for the ligands. The CHARMM36m/mTIP3P combination is expected to provide more accurate ensembles for intrinsically disordered proteins, which is the case of Aβ42[11].

***Adaptive sampling MD***

The systems were equilibrated prior to the production molecular dynamics (MD) simulations using the *Equilibration_v2* module of HTMD[7]. The system was first minimized using the conjugate-gradient method for 500 steps. Then the system was heated to 310 K and minimized as follows: (i) 500 steps (2 ps) of NVT thermalization with the Berendsen barostat with 1 $kcal \cdot mol^{-1} \cdot Å^{-2}$ constraints on all heavy atoms of the protein, (ii) 1,250,000 steps (5 ns) of NPT equilibration with Langevin thermostat and same constraints, and (iii) 1,250,000 steps (5 ns) of NPT equilibration with the Langevin thermostat without any constraints. During the equilibration simulations, holonomic constraints were applied on all hydrogen-heavy atom bond terms and the mass of the hydrogen atoms was scaled with factor 4, enabling 4 fs time step[12–15]. The simulations employed periodic boundary conditions, using the particle mesh Ewald method for treatment of interactions beyond 9 Å cut-off, electrostatic interactions suppressed for more than 4 bond terms away from each other, and the smoothing and switching van der Waals and electrostatic interaction cut-off at 7.5 Å[13].

HTMD was also used to perform adaptive sampling of the Aβ42 conformations. Production simulations were started after the equilibration cycle and employed the same settings as the last step of the equilibration. The trajectories were saved every 0.1 ns. Three adaptive sampling protocols were used. At first, in protocol A we used adaptive sampling with the *self-distance* metric for all the Cα atoms, and the time-lagged independent component analysis (tICA)[16] projected in 1 dimension. 50 ns MDs were run in 4 parallel replicas, for 10 adaptive epochs, and performed for all of the 30 models of PDB ID 1Z0Q. This corresponded to a cumulative time of 2 μs per starting structure, and 60 μs for the combined simulation. This protocol was applied to the free Aβ42 with both AMBER and CHARMM force fields. For the CHARMM force field, the simulation time was doubled to 20 epochs per starting structure, yielding a total cumulative time of 120 μs. In protocol B we used the same settings as in protocol A, but with the CHARMM force field and the following modifications: i) only the first structure of the original PDB was used; ii) 50 ns MDs were run initially in 10 parallel replicas (for the first 12 epochs) and later extended to 20 replicas, for a total of 80 adaptive epochs, corresponding to a cumulative time of ca. 74 µs (the adaptive metric was the same). Protocol B was applied to the free Aβ42 only. Protocol C was implemented by modifying protocol B as follows: i) the *secondary-structure* was used as the adaptive metric, also with the tICA projected in 1 dimension; ii) 200 ns MDs were run in 20 parallel replicas for 16 adaptive epochs, corresponding to a cumulative time of ca. 64 µs. This protocol was applied to free Aβ42, Aβ42 + TMP and Aβ42 + SPA.

***Classical MDs***

HTMD was also used to perform classical MD simulations, where only the first model of the PDB entry 1Z0Q was used. The free Aβ42, Aβ42 + TMP and Aβ42 + SPA systems were prepared and equilibrated as described above. The endpoint of the equilibration cycle was taken as a starting point for subsequent unrestrained MD simulations. These MDs were performed only with the CHARMM36m force field, with the same settings as the last step of the equilibration dynamics. Each MD was run in sequential batches of 200 ns each, for a total of 5 µs, and 10 independent replicates were performed for each system.

***MD analysis***

The ParmEd program[17] was used to convert the CHARMM topologies to AMBER topologies. The *cpptraj*[18] module of AmberTools 16[4] was used to concatenate the filtered trajectories of each system, center and align them by their backbone atoms, and save the combined trajectory in a single file. The same module was also used to compute several properties in the combined ensembles, such as the root-mean square deviation (RMSD) and radius of gyration ($R_g$). The *do_dssp* module of Gromacs 5.1[19], patched with DSSP 3.0[20], was used to compute the secondary structures for every snapshot of the combined trajectories and the total content. An in-house script that applies DSSP 3.0[20] was used to calculate the secondary elements by residue for an ensemble containing every 100$^{th}$ snapshot of each simulation. The default seven-letter DSSP alphabet was converted to the three main secondary elements: α-helix = 310-helix (G) + alpha-helix (H) + pi-helix (I); β-strand = beta-bridge (B) + beta-sheet (E); and coil = coil (“ ”, null assignment) + bend (S) + turn (T). Counting the number of residues of each secondary element in the peptide for all the snapshots analyzed thus resulted in the total secondary structure content of the ensemble. The analyzed snapshots were also grouped in ten correlative bins, the average secondary structure type content values of each residue were calculated for each bin, and the dispersion of each secondary structure type value was reported as the standard error (as a proxy of the certainty of the content value calculated, for each residue, for that particular secondary structure type).

***Ligand-peptide interactions***

The binding free energy ($\Delta G_{\text{bind}}$) of each individual residue of Aβ42 with the total number of ligands was calculated by the linear interaction energy (LIE)[21]. This was computed with *cpptraj*[18] of AmberTools

16 for every snapshot of the combined MDs, and it was expressed as the respective electrostatic and van der Waals components.

#### *Intra-peptide interactions*

The molecular mechanics/generalized Born solvent accessible surface area (MM/GBSA) method[22,23] was used to calculate the intramolecular interaction energy within the Aβ42 peptide. The a*nte-MMPBSA.py*[22] module of AmberTools 14[24] was used to convert the original topology of the system, remove all the waters, small molecules and ions, specify the Born radii as *mbondi3*, and generate the corresponding topology file to be used in the MM/GBSA calculations. *Cpptraj* was used to remove all the small molecules from the combined MD trajectories. The *MMPBSA.py.MPI*[22] module of AmberTools 14 was used to calculate, in parallel, the free energy of the peptide for every frame of the ensemble. The generalized Born method (*&gb* namelist) was used with implicit generalized Born solvent model (*igb=8*) and 0.1 M ionic strength (*saltcon=0.1*). The solvent accessible surface area was computed with the LCPO algorithm[25]. Decomposition of the pairwise interactions was generated *(&decomp* namelist), with discrimination of all types of energy contributions (*idecomp=4*) for the whole residues (*dec_verbose=0*).

### Comparative Markov State Model Analysis (CoVAMPnet)

#### *Conventional Markov state models*

The simulations were compiled into a simulation list using HTMD[26], the water and ions were filtered out, and the unsuccessful simulations with length less than 50 ns (protocol A) or 200 ns (protocol C) were discarded. Such filtered trajectories were combined, which resulted in cumulative simulation times of 60 μs (protocol A with the AMBER force field), 120 μs (protocol A with the CHARMM force field), ca. 74 μs (protocol B), or ca. 64 μs (protocol C). The conformational dynamics of Aβ42 was studied by the same metric used in the adaptive sampling: the *self-distance* of the Cα atoms of the protein, the *secondary-structure* in the simplified form (3-letter DSSP alphabet), or even a combination of those two. Over the different attempts, the dimensionality of the data was reduced with tICA to a range of 1 to 5 dimensions and lag times from 2 to 10 ns; the data was then clustered using the MiniBatchKmeans algorithm to 1000 clusters. We made several attempts at the classical construction of Markov state models (MSM) from the different data projections. We used lag times ranging from 20 to 30 ns to generate between 3 to 8 states, and the Chapman-Kolmogorov test was performed to assess further the quality of the states.

***Learning Markov state models using neural networks***

The Variational Approach to Markov Processes (VAMP)[27] was used to estimate MSMs by learning neural network (VAMPnets[28]) with physical constraints[29]. In this work, we used the implementation provided by Löhr and co-authors[30], which adopts the self-normalizing set-up[31]. The network architecture consists of 2 parts. The first part ($\chi$-*network*) computes a nonlinear function $\chi$ that maps a representation of the 3D structure from a given MD simulation frame to a lower-dimensional feature space. The elements of the output vector measure the probability of the input frame belonging to the individual states in the estimated Markov model. The second part of the network implements physical constraints to ensure that the learned Markov model is reversible and that the matrix of the governing Koopman operator[28] (a linear operator propagating in time the probabilities of the simulation frames belonging to particular Markov states) has non-negative elements. Details are given next.

*The $\chi$-network*

The VAMPnet architecture consists of two identical, weight-sharing network lobes. During training, the frame $\xi_t$ (representation of the peptide structure at time $t$ of the simulation) is passed on to the first lobe and the frame $\xi_{t+\tau}$ is passed to the second lobe. The value of the lag time separating the frames is fixed ($\tau$ = 5 ns). The frame representation is the vectorized upper triangle of the peptide's inter-residue heavy atom distance matrix without its diagonal and the first two subdiagonals (i.e. the distances of the first and second neighbors). During inference, a single frame is input to the first lobe, to classify it into one of the learned states. The architecture of the lobes contains 5 fully connected (FC) hidden layers with 256 neurons each, and one FC output layer with the number of nodes corresponding to the number of states *M* in the constructed MSM (we experimented with values of *M* between 2 and 5). The first 5 FC layers are activated using scaled exponential linear units. The output layer is activated using softmax, which ensures the output score to be interpretable as a probability of a frame belonging to the state associated with the output node. Batch normalization is used for the input of the network and the weights of the FC layers are initialized using the LeCun initialization[32]. Early stopping is used to stop the training whenever the VAMP score goes down by more than 0.001. The weights are optimized using the Adam optimizer with $\epsilon$ = $10^{-4}$ (constant used to prevent any division by zero), exponential decay rates $\beta_1$ = 0.99 and $\beta_2 = 0.999$ and $clipnorm$ 1.0. L2 regularization with weight $10^{-8}$ is applied to all trainable layers.

*Implementing the physical constraints*

The physical constraints are implemented by two layers called u and S. Both layers ensure that the resulting operator will be physical; they correspond to expressions combining covariance matrices of the input data passed through the χ-network with trainable weights specific to the layers. For more details and precise formulas regarding these constraint layers, see ref. [29].

*Estimating the Koopman operator*

The Koopman operator $K$ performs the temporal propagation between states as

$$E[\chi(\xi_{t+\tau})] \approx K^T E[\chi(\xi_t)] \quad (1)$$

where $\xi_t$ a vector of inter-residue distances representing the structure of the peptide at time $t$ of the simulation, $\xi_{t+\tau}$ represents the structure at simulation time $t + \tau$, $\chi(\cdot)$ is a nonlinear function learned by the χ-network, which maps the simulation frame representations to M-sized vectors of probabilities of the frame belonging to one of the M states, K is a $M \times M$ matrix representing the Koopman operator, and $E$ is the expectation over all possible conformations of the peptide (in practice approximated by the empirical distributions of the conformations observed in the simulations). In other words, the Koopman operator approximately propagates the expected state soft assignments $E[\chi(\xi_t)]$ at time t to time $t + \tau$. The matrix representation K of this operator for a given system is constructed as previously described[29].

*Training the network parameters*

The training of VAMPnet models is performed in an unsupervised fashion, i.e. without any manual or pre-computed labels. The network is trained using loss functions[27], which captures the dynamic requirements for building an MSM. These loss functions are derived from the assumption of Markovianity of the underlying process using VAMP[27].

*Training a single model*

The VAMPnet architecture is trained in three steps. Firstly, the χ-network is trained using the symmetrized VAMP-1 score[27] and learning rate of $5 \cdot 10^{-2}$ followed by training using VAMP-2 score and learning rate of $1 \cdot 10^{-3}$. Secondly, the χ-network weights are frozen and only the constraint layers are trained using the VAMP-E score with learning rate of $5 \cdot 10^{-4}$. Lastly, both the χ-network and the constraint layers are trained together, using the VAMP-E score and learning rate $2.5 \cdot 10^{-4}$.

*Training the model ensemble*

For each of the three systems (1. free Aβ42, 2. Aβ42 + TMP, and 3. Aβ42 + SPA), 60 VAMPnet models are trained from their adaptive sampling simulations (simulated using protocol C). First, 20 random splits of the simulated frames from a given system are generated, with 90% of the frames used for training and the remaining 10% for validation. For each split, 3 VAMPnet models are trained, where only the one with the highest VAMP-E score gets selected for the successive use. The ensemble of the selected 20 models is used to compute the probability of a given simulated frame belonging to each of the 3 states via soft assignment. The state with the highest probability is selected as the hard assignment of the frame. Note that the trained models, and thus also the states, are specific to each system (see below).

#### *Equilibrium distributions, kinetic rates and model validation*

The equilibrium distribution (or population) is the eigenvector of the Koopman matrix $K$ corresponding to its highest eigenvalue (which is equal to 1). The mean first-passage time $T_M^{ij}$ between states $i, j$ can be derived from the Koopman matrix using the transition path theory[33]. The transition rates are computed as the inverse of the mean first-passage times. For model validation, the slowest implied timescales $t_i$ (also known as relaxation times) of the MSMs are studied. They are obtained from the model lag time $\tau$ and the $i^{\text{th}}$-highest eigenvalue $\lambda_i$ of $K$ as

$$t_i = -\frac{\tau}{log(\lambda_i)} \tag{2}$$

This quantity is physical and should not depend on our choice of parameter $\tau$, therefore the value of $\tau$ for the model is selected such that $t_i(\tau)$ is approximately constant around our selected lag time. For the construction of $t_i(\tau)$, the Koopman matrix $K$ is explicitly constructed with different choices of $\tau$. To this end a $\chi$-network trained under a fixed value of lag time $\tau$ = 5 ns is used and only the lag time values in the training of the constraint layers are changed. Koopman matrices for different lag times are constructed using the constraint layers trained with the different values of lag times. To verify that the discretization of the underlying Markov process into an MSM with a particular value of $\tau$ does not significantly disturb the Markovianity, the Chapman-Kolmogorov tests are performed. I.e., it is verified that an $n$ times repeated propagation of a state population $\chi$ by $K(\tau)$ ($K$ estimated for a given $\tau$) is comparable to a propagation by $K(n\tau)$, i.e.

$$K^n(t)\chi \approx K(nt)\chi \tag{3}$$

#### *Alignment of learned states for comparative analysis*

The use of an ensemble of VAMPnet models ($N$ = 20 in our case) for each system and comparison of learned MSMs between different systems motivate a method for finding correspondences between states. In this section, we enhance the approach from Löhr et al. [23] for the alignment of states within a single system and introduce a new method for aligning MSMs between different systems.

##### *Aligning states within a single system*

The $N$ ($N$ = 20 in our case) VAMPnet models trained for a single system provide $N$ different MSMs. While all these MSMs should contain similar states, each MSM is estimated independently, and the numbering of the states between different MSMs may not correspond. Therefore, the states need to be aligned across the $N$ MSMs. Each state $m \in \{1, \dots, M\}$ of an MSM with $M$ states, can be characterized by its average inter-residue distance matrix $D_m^n$ computed as:

$$D_m^n = \sum_{\xi_t \in S} \frac{\chi_m(\xi_t)}{Z} \xi_t \qquad (4)$$

where $n$ $(1 \leq n \leq N)$ identifies the instance of the VAMPnet model used for the estimation of the MSM, $S$ is the set of all simulation frames of a system $s$, $\xi_t$ is the inter-residue distance representation of all residue pairs for a frame at time $t$, which is weighted by term $\frac{\chi_m(\xi_t)}{Z}$, where $\chi_m(\xi_t)$ is the probability of $\xi_t$ belonging to state $m$ and $Z = \sum_{t \in \phi} \chi_m(\xi_t)$ is a normalization constant .

The $N$ MSMs with $M$ states each provide $N \cdot M$ matrices $\{D_m^n\}_{n=1,m=1}^{N,M}$ for each system $s$. The goal is to group the $N \cdot M$ matrices $D_m^n$ into $M$ clusters with a constraint such that matrices corresponding to the same MSM cannot be assigned to the same cluster.

This is implemented by a constrained k-means clustering with M cluster centers. In our approach the cluster centers are initialized by $M$ inter-residue distance matrices $D_m^n$ corresponding to a single randomly selected MSM. Next, the clustering iterates the following two steps. In the first step, matrices $D_m^n$ are assigned to cluster centers respecting the constraint that two matrices from the same model cannot be assigned to the same cluster. This is implemented by greedily assigning matrices $D_m^n$ from the same model in the order of increasing distance to the closest available cluster center, while respecting the above constraint. In the second step, the cluster centers are recomputed by taking the mean of the matrices $D_m^n$ assigned to each cluster. These steps are iterated until there is no change in the cluster assignments between two iterations. The outcome is a set of matrices $D_m^n$ for each of $M$ clusters. These matrices are then considered as representing the same state of the given simulated system. This method is different from previous work[30], which used one of the learned models as a reference, i.e. their approach was equivalent to only performing the first iteration of our method. The proposed approach can obtain potentially a better solution less susceptible to an incorrect initialization.

*Aligning ensembles of Markov state models between different systems*

This method is fully described in the main text, and hence it is omitted here.

***Gradient-based characterization of learned states***

This method is fully described in the main text, and hence it is omitted here.

***Estimation of the free energy landscape***

We estimate the Aβ42 free energy landscape for each of the studied systems by performing Gaussian kernel density estimation on 10% of the simulated frames[30]. We perform this estimation over the tICA space representation of the frames, as this allows for an efficient projection of the free energy landscape to the plane given by the first 2 tICA dimensions, which are relevant for the slowest state transitions.

**Experimental validation**

***Production and purification of monomeric Aβ42***

Aβ42 in its monomeric form was produced and purified following an adapted version of the protocol by Cohen et al.[34]. BL21-DE3 cells were transformed with pET-Sac-Abeta(M1-42) (71875, Addgene) encoding N-methionylated Aβ42 (N-Met-Aβ42) and plated on LB/agar supplemented with 100 µg/mL ampicillin. Following the overnight growth at 37 °C a 10 mL preculture was inoculated with a transformed colony and incubated at 37 °C overnight. Protein production was initiated by transferring 1/10 (v/v) of the preculture into 1 L of fresh LB media supplemented with ampicillin. Cells were incubated at 37 °C and the protein expression induced by 0.5 mM IPTG when optical density reached ca 0.7. All incubations were performed with vigorous shaking (180-220 rpm). Following the overnight (ca. 16 h) expression at 37 °C, cells were harvested at 6,000 x g for 20 min at 4 °C and frozen. $^{15}$N-labeling was performed using M9 minimal medium instead of LB medium and $^{15}NH_4Cl$ as the sole source of nitrogen. Pre-culturing was performed in two steps instead of one: first 10 mL of LB + ampicillin over the day followed by 100 mL of M9 overnight, both at 37 °C.

Cells were resuspended in 10 mM Tris 1 mM EDTA pH 8.5 (purification buffer, PB), sonicated on ice for 15 min (30 s on, 60 s off) at 60% amplitude then centrifuged at 18,000 x g for 30 min at 4 °C. The supernatant was discarded, and the pellet resuspended in 30 mL of fresh PB. This process was repeated two more times. The final pellet was resuspended in PB containing 8 M urea, incubated for 20 min at 8 °C under gentle stirring, sonicated on ice for 6 min (30 s on, 60 s off) at 60% amplitude then centrifuged at 18,000 x g for 30 min at 4 °C. The supernatant was diluted to 2 M urea by the PB, and loaded onto a gravity column containing 5 mL of an anion exchange resin (DEAE Sepharose Fast Flow, Sigma Aldrich) equilibrated in PB + 2 M urea. The flowthrough was loaded to a second column. Protein on both columns was eluted separately using PB + 2 M urea, PB, PB + 10 mM NaCl and PB + 100 mM NaCl. Protein yields were assessed using tris-tricine SDS-PAGE (15% acrylamide). All fractions containing Aβ42 were pooled, and loaded onto a HiLoad 16/600 superdex 75 pg (Cytiva). Purified Aβ42 eluted between 70 and 90 mL. All fractions were pooled and lyophilised in glass vials for long-term storage. Prior experiments, lyophilised Aβ42 was resuspended in 6 M guanidium hydrochloride pH 8 and loaded onto a GL 10/300 Increase 75 pg column (Cytiva) equilibrated in 20 mM sodium phosphate pH = 7.4. Monomeric Aβ42 was collected between 13.6 and 14.6 mL on ice and used within max 4 hours for the downstream experiments.

*Secondary structure determination using circular dichroism (CD)*

CD spectra of 37 μM monomeric Aβ42 were measured at 10 °C using Chirascan CD spectrometer (Applied Photophysics, USA) in a quartz cuvette. The CD spectra were measured in the far-UV range between 185 and 260 nm in 1 nm increments with 1 nm bandwidth, and 1 s integration time. Each spectrum was collected in triplicate and averaged. The buffer-baseline subtracted data were used for spectral deconvolution and secondary structure analysis using BestSel (https://bestsel.elte.hu/)[35]. 37 mM small molecules (1000-fold molar excess) and HFIP were added where indicated.

*Secondary structure determination using Fourier-transformed infrared spectroscopy (FTIR)*

FTIR with attenuated total reflection (ATR) spectra of the samples were collected using a Bio-ATR II unit mounted to the Invenio FTIR spectrometer (Bruker Optics, Germany). Measurements were carried out at 15 °C in 20 μL volumes. The signal from 60 μM Aβ42 without or with 60 mM of small molecules was corrected for the absorption of the buffer and other components (i.e., TMP, SPA) by subtraction of the precisely matched buffer spectrum. Each spectrum was measured at 4-7 repetitions. Gaussian deconvolution of each spectrum was carried out in OriginPro. Second derivative spectra were used for estimation of the number of peaks, which were fixed during the fitting. Samples containing hexafluoroisopropanol (HFIP) could not be measured due to strong absorption of the solvent in the amide regions.

*Nuclear magnetic resonance (NMR)*

NMR spectra were recorded using 850 and 950 MHz Avance NEO NMR spectrometers (Bruker) equipped with 5 mm triple resonance ($^1H$, $^{13}C$, $^{15}N$) inverse cryoprobe and cooled $^1H$ and $^{13}C$ preamplifiers. Samples containing 69 μM (Aβ42 alone), 58 μM (with 58 mM TMP), or 55 μM Aβ42 (with 55 mM SPA), were adjusted to 10% $D_2O$ to perform lock. Pulse sequences from TopSpin (Bruker) library were used. All experiments were performed at 4 °C. Chemical shift perturbation (CSP) was computed[36] as:

$$\mathrm{CSP} = \sqrt{(\Delta\delta(\,^{1}\mathrm{H}))^{2} + (\frac{1}{6.5} \times \Delta\delta(^{15}\mathrm{N}))^{2}} \qquad (5)$$

Secondary structure probabilities were calculated from $^1H$-$^{15}N$ HMQC peak assignment, using the chemical shift indexing (CSI) method with the CSI3.0 web server.[37]

*Aggregation kinetics assay*

10 μM Aβ42 was mixed with pH-adjusted 10 mM TMP or 10 mM SPA, HFIP, and 15 μM of thioflavin T (ThT) on ice. Each mix was loaded in triplicates on a pre-cooled 384-well flat bottom low binding

microplate (Corning, USA) and ThT fluorescence was monitored at excitation and emission wavelengths of 400 nm and 485 nm, respectively, using a Synergy H4 microplate reader, at 37 °C. The curves were normalized and fitted with OriginPro (OriginLab) to the equation[38]:

$$y = y_0 + A/(1 + e^{-k(t-t_{0.5})}) \qquad (6)$$

where *A* is the amplitude, *k* the apparent aggregation rate, *t* is time, and $t_{0.5}$ the aggregation half-time.

### SUPPLEMENTARY NOTES

#### Supplementary Note 1. Effective concentration of small molecules in MD simulations

Kocis and Hey et al. reported that both TMP and SPA could exert their largest effects on Aβ42 only at a high molar excess (1000-fold or higher)[39,40]. These works used in their *in vitro* experiments 22 µM of Aβ42 and up to 22 mM of TMP or SPA. In all our experiments (CD, FTIR, and NMR) the concentrations were only slightly higher (37-69 mM of small molecules for 37-69 µM Aβ42). In our simulations, we had to find a compromise between having a large excess of those ligands and keeping the systems with a manageable size to allow the simulation of reasonable time scales. Therefore, we added 100 molecules of ligands per molecule of peptide (i.e., 100-fold excess) instead of 1000. Nonetheless, the final concentration of those ligands in our *in silico* systems was approximately 250 mM, which is higher than what was used *in vitro*, but not excessively higher (between 4- to 7.5-fold). Thus, we expect that a sufficient number of interactions between the small molecules and the peptide could be observed *in silico* (possibly, helping the equilibrium be reached faster), without exerting an artificial bias due to the concentration differences.

#### Supplementary Note 2. Exploration of different adaptive sampling protocols

To date there are only a few experimental structures of the Aβ42 peptide in its full length obtained from liquid state NMR: PDB IDs 1Z0Q[41] (**Supplementary Figure S1**) and 1IYT[42]. 1Z0Q contains a larger ensemble than 1IYT (30 and 10 structures, respectively), which implies a larger conformational diversity, and that is why it was selected. These structures have been determined in the presence of solubilizing agent hexafluoroisopropanol (HFIP), a known inducer of α-helical structures that prevents aggregation of Aβ42. HFIP creates an apolar environment similar to the lipid phase of membranes so it can be assumed that the conformations of Aβ42 in this PDB are similar to those adapted by the peptide upon its cleavage from the transmembrane domain[41]. In these conditions, the Aβ42 peptide contains solely α-helices (42%) and coils (58%), in contrast to aqueous solution conditions (buffer without HFIP) where Aβ42 contains 67% of coils, 26% β-strands, and only a small fraction of α-helices (6%) (experimental data from this work and refs. [41,43]). Therefore, in our simulations of free Aβ42, we expected to observe the transition from the helical form to coils and β-structures.

In order to assess the ability of the MDs to explore the conformational diversity of Aβ42 we attempted three different protocols, which differed in the starting structure set, the adaptive metric, the number of adaptive epochs and replicas, and the total cumulative MD time (**Supplementary Table S1**). Their performance regarding the exploration of Aβ42 conformational diversity was assessed by comparing the relative abundance of secondary structure elements with that in the starting structure and with experimental results[41,43].

In our first approach (protocol A; see Supplementary Materials and Methods above), we simulated the free Aβ42 in water for 60 µs using both AMBER ff14SB[10] (A14SB) and CHARMM36m[11] (C36m) force fields. These two force fields were selected based on the literature[44], which suggested several force fields to study Aβ, including A14SB and C36m. From these, we tested A14SB, since it was readily available to us, and C36m because it was developed very specifically for IDPs[11] and it had already been used before with Aβ42[45]. We first used the conventional Markov state models (MSMs) to analyze the ensembles obtained in both cases. Although the constructed models did not exhibit Markovian properties, we still could cluster the trajectories and compare the dominant conformations. The results showed that the simulations with A14SB produced well-structured clusters, while the C36m produced much more unstructured and varied clusters (**Supplementary Figures S2-S3**). The Aβ42 peptide is known to be highly flexible and disordered[46], and we expected to observe a large diversity of conformations and orientations of the peptide. For this reason, we concluded that the A14SB force field is not suitable for capturing the conformational ensemble of Aβ42 properly, and hence we decided subsequently to use only the C36m force field. Paul et al.[47] have recently reported an assessment of several force fields for the simulation of the Aβ42 peptide. They have concluded that CHARMM22* is probably the best one to reproduce spectroscopic observables of Aβ42 but also that C36m is reasonably accurate, which validates our choice.

It has to be noted that the resulting ensembles are biased towards the starting structure, which is dominantly helical and coiled. For this reason, the global properties obtained after several microseconds of simulation are still far from the experimental results, which were obtained in very different time scales (minutes/hours). We expect, however, to observe significant differences among the systems, and, more importantly, to describe the influence of the small molecules on the Aβ42 conformations. In order to assess if the experimental results could be approached by simulation, we attempted three different approaches.

First, the protocol  described above using C36m as a force field was extended to a total time of 120 µs (protocol A, running in parallel 30 adaptive MDs with the metric of *self-distance* of all Cα atoms). The assessment of the secondary structures showed a striking similarity of the final ensemble and the initial state, with nearly the same content of α-helix and coil and only 2% of β-strands, in great contrast with the experimental results (**Supplementary Figure S4**). This meant that the simulation did not diverge very much from the membrane-like conformation of Aβ42 present in the original NMR structure (**Supplementary Figure S1**).

Second, we sacrificed the diversity of starting conformations for a longer adaptive exploration. We performed adaptive sampling of the free Aβ42 with one single structure but with a higher number of replicates and epochs, in a total cumulative time of 74 µs (protocol B). This protocol improved the results, achieving a higher ratio of β-strands (6%), and a lower but still high amount of α-helices (27.5%).

Third, in an attempt to enhance the sampling of different secondary elements in Aβ42, we changed the adaptive metric to the *secondary-structures* and increased the length of each individual MD to 200 ns and simulated in total approximately 64 µs (protocol C). This time we obtained an even lower ratio of α-helices (17%) and a higher amount of coils (77.5%). Moreover, the ratios of the secondary structures plateaued after ca. 44 µs of total simulation time (**Supplementary Figure S5**). Compared with the previous two protocols, protocol C proved to be the most efficient in exploring the diversity of secondary structures and diverging from the initial helical-rich conformation found in the NMR structure (**Supplementary Figure S4**). Therefore, the latter adaptive protocol was selected to survey the conformational ensembles of Aβ42 and to study the influence of the small molecules TMP and SPA on such ensembles.

**Supplementary Note 3. Evolution of secondary structure elements over time**

The adaptive simulations consist of many individual trajectories, simulated over several epochs. Before each epoch, the seeds for the new simulations are chosen from selected snapshots from previous simulations, according to the predefined criterion. The objective is to maximize the variability of the simulated trajectories in terms of the specified feature (in this case, the secondary structure).

To assess the evolution and characteristics of the generated data, based on the total continuous simulation time, we set a reference timepoint as the beginning of the first simulation and we align all the other simulations with respect to this timepoint - owing to the fact, that the initial frame of the latter simulations must be present in some previous simulation. As a result, we obtained a direct account of the continuous simulation times performed immediately after the equilibration of the system, and the population distribution of simulation frames in terms of their timestamp (**Supplementary Figure S6**). We found that the longest concatenated trajectories consisted of ca. 750 ns for the free Aβ42, 900 ns for Aβ42 + TMP and 1 µs for Aβ42 + SPA. Although the global dynamic ensembles correspond to ca. 64 µs, this may have implications on the events and transitions sampled here, especially if we consider that the starting conformation of Aβ42 (containing mostly coils and α-helices) is rather distant from the expected end-point state in water (known to be dominated by coils and β-strands; see below). Moreover, we found that sometimes multiple simulations started from the same snapshots, while many simulations were not utilized as seeds in subsequent epochs. This may be due to certain conformational states being rarely sampled, which led the adaptive method to select them multiple times for seeding new trajectories. This is a direct result of the exploration/exploitation tradeoff embedded in the adaptive sampling method[48].

**Supplementary Note 4. Evolution of secondary structure elements in classical simulations**

To gain deeper insights into the time evolution of Aβ42, we performed 5 μs-long MDs (10 replicates) with the three systems (free Aβ42, Aβ42 + TMP and Aβ42 + SPA). We computed the time-evolution of the secondary elements in Aβ42 for those MDs. The results showed that the helical structures tend to change into coils, in a time range of a few microseconds. Then, some of these coils tended to change into β-strands, especially around residues 15-20 and 31-40, but sometimes also around residues 3-6 (**Supplementary Figures S7-S9**). This is in agreement with the results from the adaptive simulations (**Figure 1C**). The main difference among the three systems is how quickly these transitions occur. On average, both TMP and SPA preserved the helical elements of Aβ42 for longer periods than in their absence. However, there was a high variability among the replicates, both in terms of the transition times between secondary elements and the overall propensities (**Supplementary Table S2** and **Supplementary Figures S7-S9**). Moreover, these simulations appear to be still evolving, which does not allow for attaining conclusive statistics. Globally, the Aβ42 + TMP and Aβ42 + SPA showed higher helical propensity and lower content of β-strands than the free Aβ42, although the differences were not statistically significant (*p-values* between 0.3-0.8 from *t* test). These results suggest that, although TMP and SPA may delay and shift the secondary structure of Aβ42 and reduce the formation of β-strands, such effects may not be as dramatic as the adaptive simulations suggested, especially with SPA.

**Supplementary Note 5. Meaning of aligned states across different systems**

The proposed alignment method identifies matching states from different systems including the cost of the alignment for each proposed pair of states. Since the systems differ, and different states may form and vanish in different systems (e.g., due to the addition of small molecules to the environment), the proposed match does not necessarily mean that the two states actually correspond to each other.

To decide whether two matched states correspond to each other, we set a threshold $T_e$ on the alignment cost. The alignment costs were obtained by computing the Wasserstein-1 distance of the distribution of state features over different VAMPnet instances in the model ensemble. Only pairs of states matched with the alignment cost lower than $T_e$ are considered as being aligned (**Supplementary Figure S19**). In this work, the threshold $T_e$ was set empirically to 6. In our case, the alignment cost threshold was defined from a manual comparison of the states. In general, an upper bound for the value of the threshold can be obtained by computing the self-alignment costs $c_{ij}^{s_1 s_1}$ for all states $i, j$ of system $s_1$ and selecting the lowest alignment cost for two non-identical states as the upper bound on the threshold:

$$T_e^{MAX} = \min_{i,j \in \{1,\ldots,M\}, i \neq j} c_{ij}^{s_1 s_1} \qquad (7)$$

where $M$ is the number of states of system $s_1$.

The intuition for this upper bound is that two aligned states between two systems should not have a higher alignment cost than (any) two different states of the same system. For example, if state $A_1$ from system $s_1$ and state $A_2$ from system $s_2$ have an alignment cost of 10, but aligning two different states $A_1$ and $B_1$ of system s1 has a cost of 8, then the states $A_1$ and $A_2$ should not be aligned, since the cost would be higher than the cost of alignment of states that are considered as different in this system. In other words, the cost of the alignment of states within one system provides a reference upper bound value for what should be considered as one state when comparing different systems.

In our case, the upper bound is computed from the 3-state MSM of the reference free Aβ42 system (and has a value of 8.4). Using this upper bound directly as our threshold instead of our manually selected threshold $T_e = 6$ would lead to the same alignment results (see alignment costs in **Supplementary Figure S19**).

**Supplementary Note 6. Time-based evolution of the states**

We assessed how the concatenated adaptive trajectories evolved over time in the context of the state assignment. We found that some states were more prominent in the early trajectories and others in the late ones (**Supplementary Figure S20**). In general, there were not many apparent transitions within the individual simulations. On the other hand, many segments were frequently repeated in the concatenated trajectories, which was due to the criterion applied by the adaptive method for the selection of the seeding frames. We also analyzed the soft assignment of the states along the temporarily aligned trajectories and the development of the state probability over time (**Figure 2B**). For short simulation times, the distribution of states had low confidence due to the low number of snapshots in this time range. For Aβ42 and Aβ42 + TMP, the systems started with state 1 (source, more similar to the initial NMR structure), and the number of states sampled changed quickly. Then the change in the distribution of states decelerated with time, almost reaching an equilibrium. For longer time ranges, however, the assignment of states also decreases its confidence, due to the reduced number of snapshots. Interestingly, for Aβ42 + SPA, state 2 (blue) was found in abundance in short time ranges, and almost disappeared in longer times. This could indicate that state 2 in Aβ42 + SPA (with the highest ratio of α-helices and lowest coils) is metastable and does not survive long time scales. It can also result from the borderlines between states 2 and 3 (purple)being somewhat unclear. This suggests that possibly this system could be described by only two states.

**Supplementary Note 7. Computation of the confidence intervals using ensembles of Markov state models**

Each of the models in the ensemble has learned a separate Markov State Model. We can evaluate each studied quantity by these 20 models separately, giving us 20 different values. These 20 values are used for estimation on the confidence intervals with the *numpy.percentile* function set to compute the 2.5 and 97.5 percentile values. This partly suppresses the effect of potential outliers by replacing the min value with a value interpolated between the lowest and the second lowest values and the max value with a value interpolated between the highest and the second highest values; the area between these two interpolated values is shaded in **Supplementary Figures S13, S14, S17**, and **S18** and denoted by an error bar in **Figure 2.**

**Supplementary Note 8. Intramolecular interactions of Aβ42**

The intramolecular interactions within the Aβ42 peptide were calculated using the molecular mechanics/generalized Born solvent accessible surface area (MM/GBSA) method[22,49]. The electrostatic components prevailed over the van der Waals ($\Delta E_{electrostat}$ and $\Delta E_{vanderWaals}$, respectively; **Supplementary Table S4**) in their contribution to the internal free energy of Aβ42. Moreover, the favorable polar solvation energy ($\Delta G_{polar\ solv}$) outweighed all the other contributions to the total free energy ($\Delta G_{total}$; **Supplementary Table S4**). The peptide was more stable (lower mean total energy) in the presence of TMP or SPA ($\Delta G_{total}$ = -310.4 ± 39.2 and -326.6 ± 27.2 kcal/mol, respectively) than in free solution (-299.0 ± 33.6 kcal/mol). Despite the large variability (standard deviations), these differences are very significant (*p-values* $< 10^{-4}$, from *t* test). Dissecting the different contributions to those energy differences, we found that the polar solvation energy ($\Delta G_{polar\ solv}$) contributes the most. This term is more negative in the presence of the small molecules, indicating that the small molecules prompt the exposure of more polar residues to the solvent than those in free Aβ42. This effect is concomitant with a weakening of the electrostatic interactions (less negative $\Delta E_{electrostat}$) and an strengthening of the van der Waals interactions (more negative $\Delta E_{vanderWaals}$), indicating an increase of the internal hydrophobic contacts in the presence of TMP or SPA. This is consistent with an overall increase of the compactness of the peptide, as we observed by the lower $R_g$ values (see *Secondary structure content in simulations, Comparison of conformational states across the systems*, and **Figure 1D**).

**Supplementary Note 9. Changes in secondary structure in presence of drug candidates and membrane mimetic**

The different experimental techniques produced different secondary structure propensities for N-Met-Aβ42 (**Supplementary Figure S26**). There may be several reasons for such differences.

Aβ42 seems to be adsorbed on the FTIR-ATR crystal surface (**Supplementary Figures S28A-C**), which could explain why FTIR measures a higher β-content than CD and NMR. Aβ42 is known to interact with many surfaces (e.g., quartz) which may introduce errors in the experimental assessment of its secondary structure[50]. Moreover, successive data acquisitions on the same sample revealed an evolution in time, most likely due to non-specific interactions between N-Met-Aβ42 and the ATR crystal surface.

Deconvolution of CD spectra was carried out using the BestSel algorithm[35], which was trained using spectra of folded proteins but not peptides. Therefore, the estimation of the Aβ42 secondary structure may be imprecise[35]. Moreover, CD spectroscopy has, in some cases, a tendency to underestimate helical content compared to NMR[51]. In contrast, our NMR measurements of helical content are in agreement with what was obtained from our simulations (**Figure 1** and **Supplementary Figure S26**).

Nonetheless, SPA induced strong CD spectral changes in the presence of HFIP, slight changes in FTIR spectra, and CSP by NMR (**Figure 5**, **Supplementary Figure S27**). TMP had a similar effect on CD spectra as a function of HFIP concentration, a stronger effect on FTIR spectra but very little CSP. We conclude that: (1) SPA has a stronger effect on the secondary structure of N-Met-Aβ42 than TMP, and (2) their effect is enhanced by HFIP, which was used to obtain the starting structure for simulations[41], indicating that these drugs may be more efficient in the presence of the cell membrane.

**Supplementary Note 10. Discrepancies between computational and experimental results**

Some computational predictions were not confirmed by the experimental methods: 1) Aβ42 showed higher β-strand content from every experimental technique compared to our computational results, 2) TMP or SPA did not change the global secondary structure propensities, and 3) the small molecules did not prevent Aβ42 fibril formation.

To address point (1), the generally higher β-strand content obtained by every experimental technique compared to our computational results might, at least partially, come from the fact that the computational results were obtained at infinite dilution of Aβ42. This is something that cannot be achieved in our experiments, where Aβ42 molecules will interact with each other. Our simulations of the monomeric Aβ42 do not account for peptide-peptide. However, several computational works have already investigated the oligomerization of Aβ42, but such studies are resource-demanding, and they have only scarcely been applied to study the effects of drugs on the Aβ aggregation[44,52–54]. A complementary explanation comes from unexpected unspecific interactions between N-Met-Aβ42 and the ATR crystal surface, as discussed above (**Supplementary Note 9**).

Point (2) can be explained by very different time scales between the simulations and the experiments. Here we simulated each system for ca. 64 μs, which represents a considerable computational effort for current standards. The MD simulations herein performed mimic the effect of the Aβ42 peptide leaving the membrane environment, as it starts in a highly helical conformation[55]. The simulations met several convergence criteria, hence sufficient sampling was assumed. However, our experimental procedures cannot approach such a small time scale. Bench experiments were performed with Aβ42 solvated in an aqueous environment, and took minutes to hours to complete, which corresponds to six or higher orders of magnitude longer times than the simulations. It is possible that, for such long time scales, the structure of Aβ42 could evolve beyond what we simulate within tens or hundreds of microseconds, thus mitigating the initial differences observed for Aβ42 with and without the small molecules. Indeed, **Figure 1E** suggests that the secondary structure of Aβ42 + SPA slowly drifts towards higher β-strand and random coil contents over time. Thus, timescale discrepancies between simulations and experiments can explain the lack of significant change in the structural propensity of Aβ42 in our experiments.

Point (3) could be explained by a combination of the two effects mentioned above. Oligomerization and fibril formation are necessarily dependent on the monomer concentration, and these processes may not be easily extrapolated when we simulate a single Aβ42 peptide for a short time (ca. 64 μs). On the other hand, the concentrations of Aβ42 *in vitro* vs. *in vivo* are very different: between 10-100 μM in our in vitro experiments, as opposed to around 100 pM in the CSF[56,57]. At such low concentrations in vivo, Aβ42 oligomerization and fibril formation is much slower[58], in which case even a faint effect on its secondary structure, as demonstrated here, may be substantial in conferring anti-aggregation properties to TMP and SPA

### SUPPLEMENTARY FIGURES

#### Structure of Aβ42

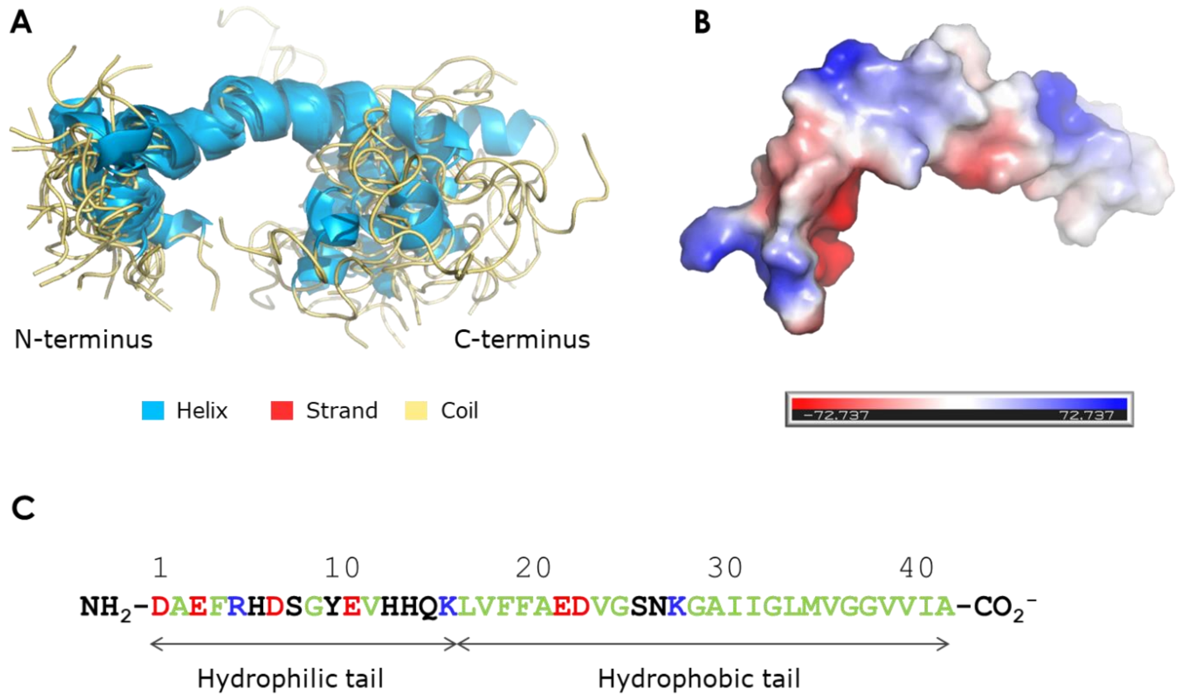

**Supplementary Figure S1. Structures of the NMR structure of Aβ42 (42-residues) used here (PDB ID 1Z0Q).** A) Superimposition of all the 30 models in this ensemble; B) electrostatic potential surface of Aβ42 (for the first model); C) sequence of the Aβ 42 peptide. The secondary structures in A) are highlighted by the colored cartoons; this PDB contains 42.1% of helices (light blue) and 57.9% of coils (pale yellow). The electrostatic potential surface was generated with PyMOL 2[59], and is represented from the exact same viewpoint; it shows the distribution of charges on the peptide surface: blue, positive, red, negative, and while, neutral. The sequence is color-coded as follows: blue, positively charged, red, negatively charged, green, hydrophobic, black, polar neutral residues.

**Comparison of the Amber ff14SB and CHARMM36m force fields**

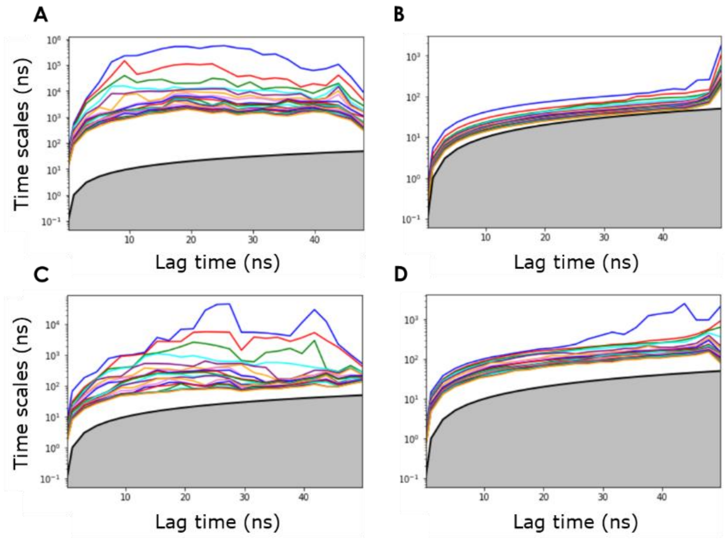

**Supplementary Figure S2. Implied time scales plots for the Markov state model analysis.** The simulations of free Aβ42 with A14SB (A, B) and C36m force fields (C, D), obtained for the projection metric of *self-distance* of all Cα atoms (A, C) and *secondary-structure* (B, D). These results were obtained for adaptive simulation with protocol A and cumulative times of 60 μs. In spite of the lack of Markovianity in most of these models (the implied time scales did not converge with increasing the lag time for most of these models), a lag-time of 30 ns was selected for the subsequent clustering (**Supplementary Figure S3**).

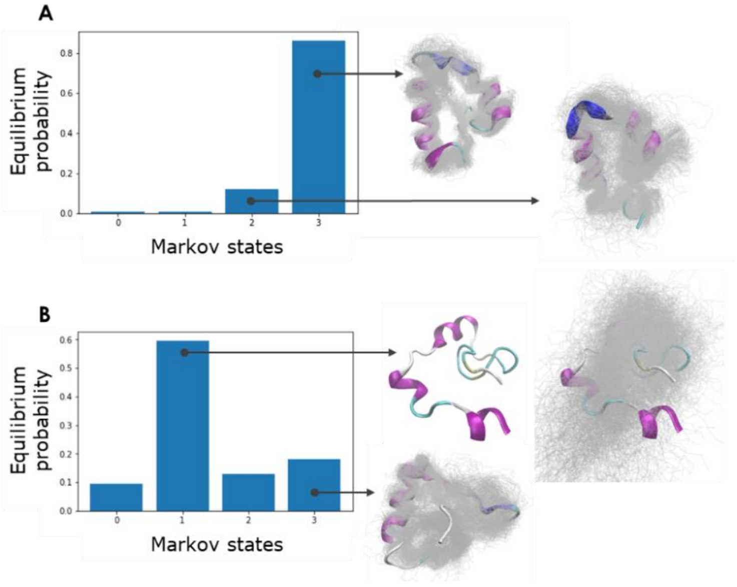

**Supplementary Figure S3. Equilibrium distribution and structures of the most populated states with the two force fields.** Markov models obtained with the *self-distance* of all Cα atoms projection metric, for the simulations obtained from protocol A and a cumulative times of 60 μs with: A) A14SB, and B) the C36m force fields. The structures of each cluster show a representative snapshot in cartoon, superimposed with other structures from the same cluster, represented by the backbone atoms shown as the gray lines. For A14SB (A) the two main states (states 2 and 3) are well clustered and structured, while for C36m the dominant cluster (state 1) is highly unclustered and diverse in orientations.

### Comparison of different adaptive sampling protocols

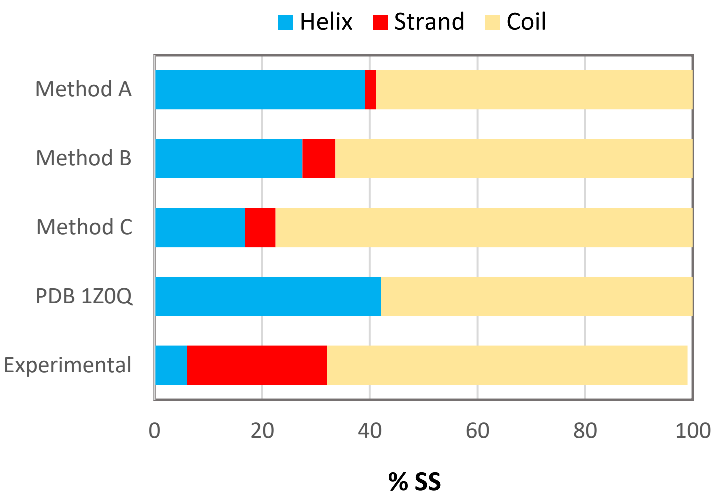

**Supplementary Figure S4. Total secondary structure content (%SS) of free Aβ42 in the simulations with the different adaptive protocols.** Protocol A (120 μs), protocol B (ca. 74 μs), protocol C (ca. 64 μs), the original NMR ensemble (PDB 1Z0Q with 30 structures), and from experimental CD measurements of free Aβ42 in aqueous solution (this work).

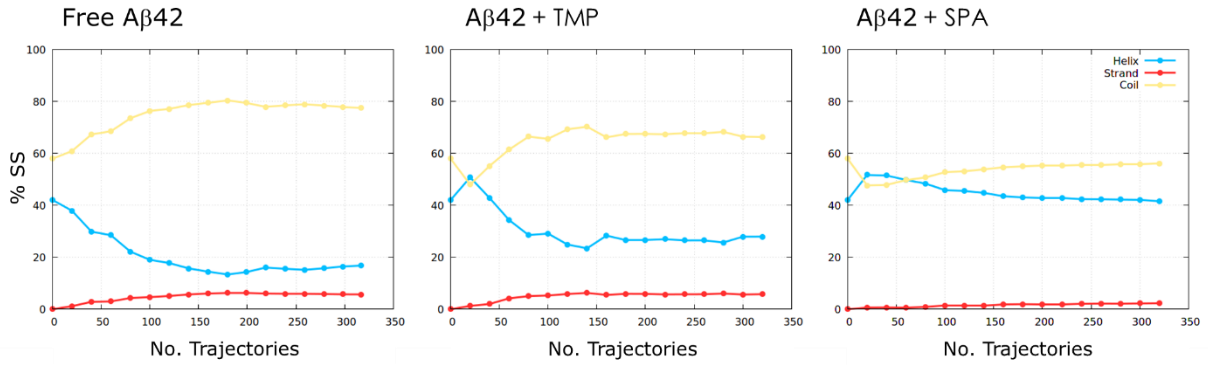

**Supplementary Figure S5. Evolution of the secondary structures with the number of adaptive trajectories.** Cumulative secondary structure content (%SS) of the simulation ensembles with increasing the number of trajectories performed over the 16 epochs of adaptive simulation for the three systems.

**Temporal alignment and concatenation of the adaptive sampling and classical MDs**

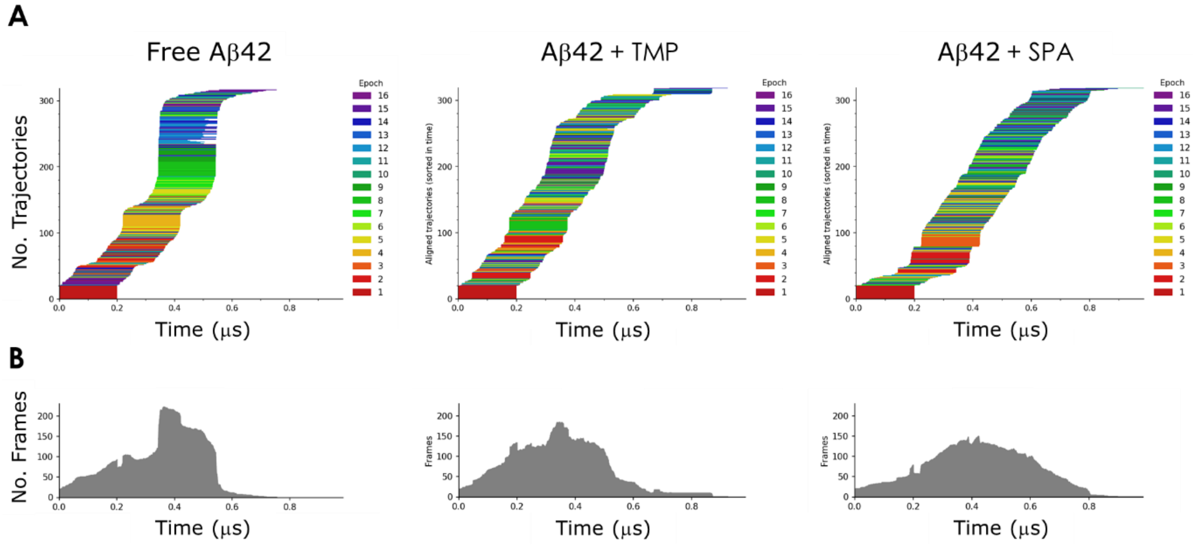

**Supplementary Figure S6. Adaptive sampling simulations aligned in time.** Free Aβ (left), Aβ with TMP (middle), Aβ with SPA (right). The simulations are sorted by the time of their first frame and colored based on their epoch. B) Total number of frames available at each time point over all the adaptive simulation.

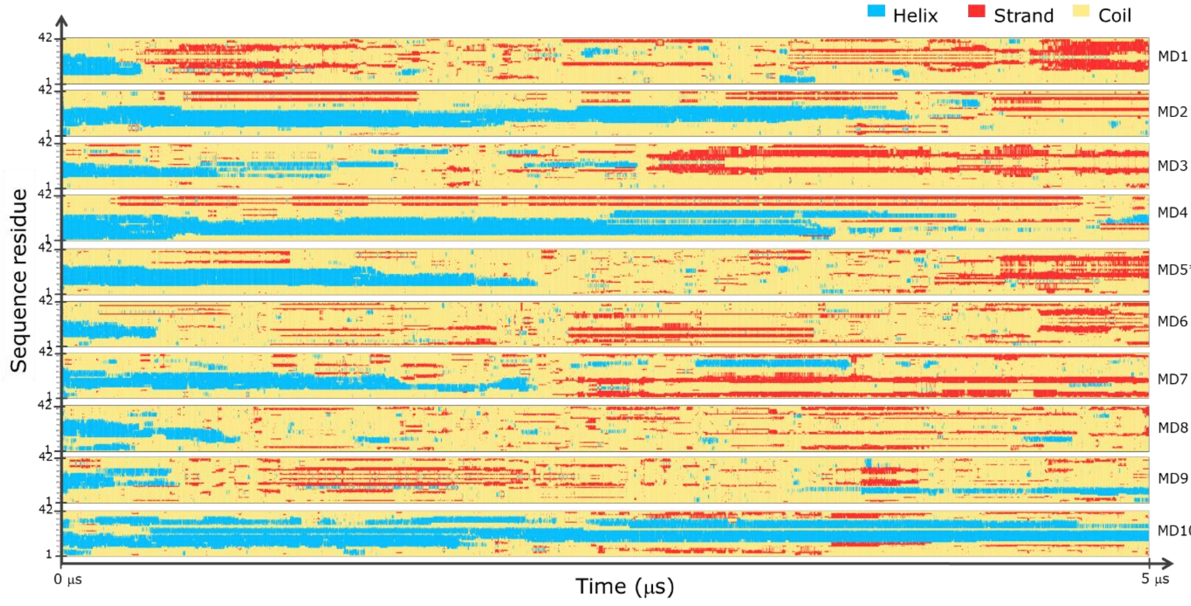

**Supplementary Figure S7. Time evolution of the secondary structures of Aβ42 in all the classical MD simulations for the free Aβ42.** *The MD with secondary propensities closest to the average over all the replicates (with lowest *RMSE*).

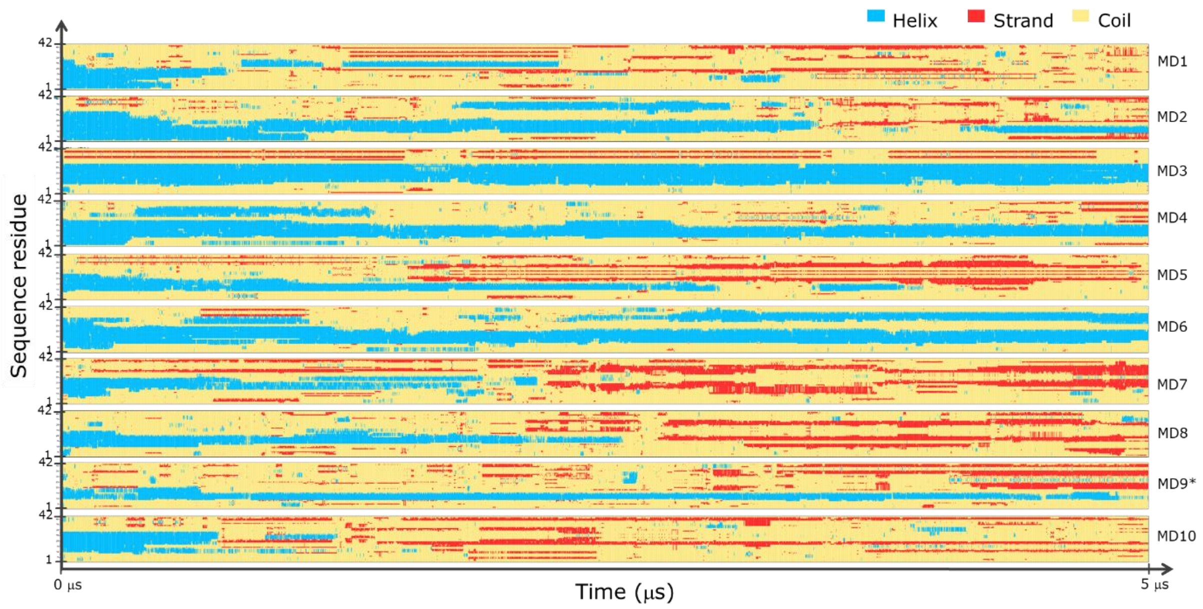

**Supplementary Figure S8. Time evolution of the secondary structures of Aβ42 in all the classical MD simulations for Aβ42 + TMP.** *The MD with secondary propensities closest to the average over all the replicates (with lowest *RMSE*).

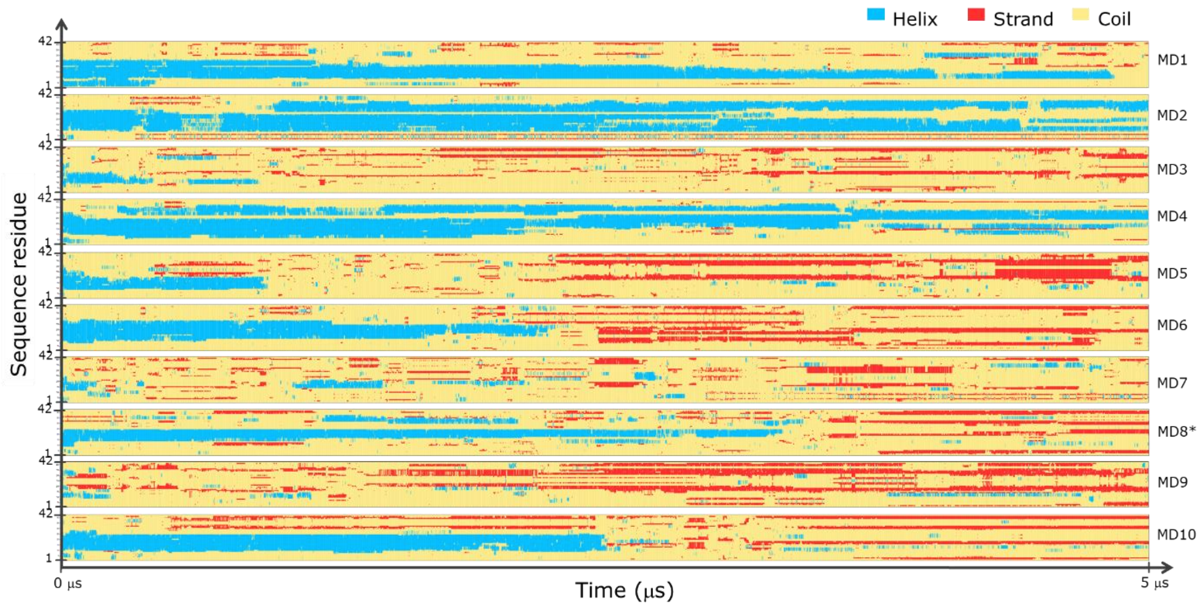

**Supplementary Figure S9. Time evolution of the secondary structures of Aβ42 in all the classical MD simulations for Aβ42 + SPA.** *The MD with secondary propensities closest to the average over all the replicates (with lowest *RMSE*).

**Conventional Markov state model analysis**

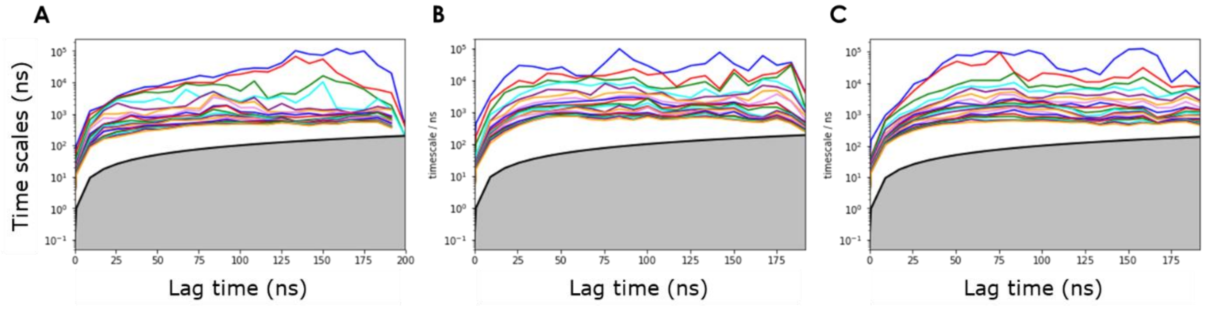

**Supplementary Figure S10. Implied time scales plots for the conventional MSMs**. Data obtained from the adaptive simulations of: A) free Aβ42; B) A$\beta_{42}$ + TMP, C) Aβ42 + 3SPA. These results were obtained for the data projected using the *secondary structure* metric, a dimensional reduction to 5 dimensions using tICA with a lag time of 10 ns, then clustered using the MiniBatchKmeans algorithm to 1000 clusters. The MSMs were constructed with a lag time 25 ns to produce 5 states. The implied time scales do not converge after a certain lag time, which reveals the lack of Markovianity. The respective Chapman-Kolmogorov tests are presented in **Supplementary Figure S11**.

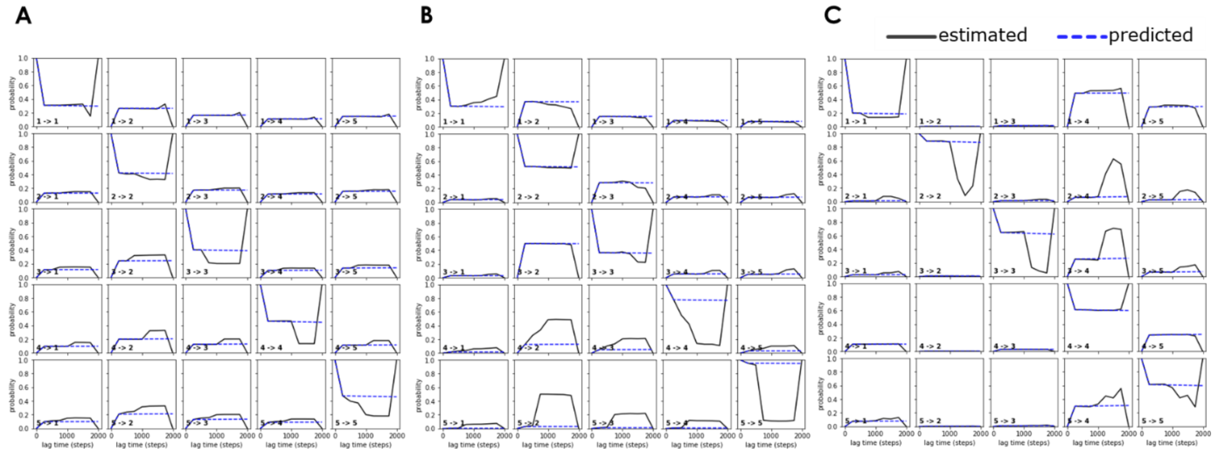

**Supplementary Figure S11. Chapman-Kolmogorov tests for the conventional MSMs**. Data obtained from the adaptive simulations of: A) free Aβ42; B) Aβ42 + TMP, C) Aβ42 + 3SPA. Tests for the analyses with the data projected by the *secondary structure* metric, tICA dimensional reduction, and MSMs constructed with a lag time of 25 ns and 5 states. The state transitions *estimated* from the data (full black lines) do not overlap with the transitions *predicted* from the model (dotted blue lines), showing the non-Markovianity of the models. The respective implied time scales are presented in **Supplementary Figure S10**. The discontinuity of the estimated transitions for higher lag times is likely due to the numerical collapse of the transition matrix to a diagonal matrix: for large lag times, the simulation may not have enough samples of the state transitions to obtain good statistics. As a result, the system will stay in each state with the probability of one. Nevertheless, the discrepancy between the predicted and estimated graphs is visible even for smaller lag times.

*MetricSelfDistance, tica.project(1), tica.lag(10ns):*

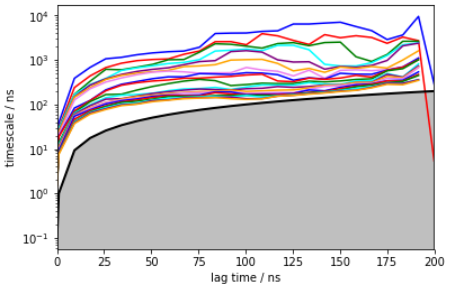

*MetricSelfDistance, tica.project(1), tica.lag(50ns):*

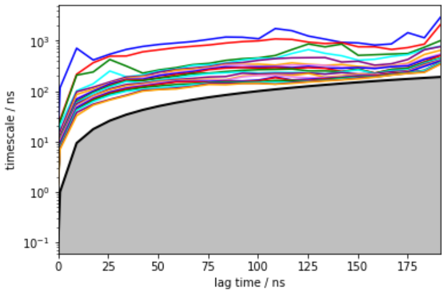

*MetricSelfDistance, tica.project(5), tica.lag(50ns):*

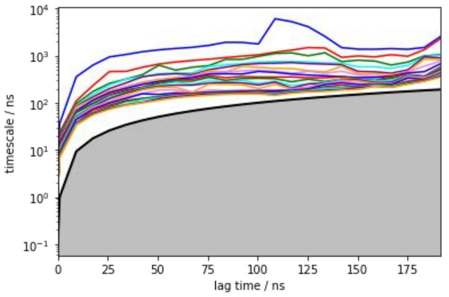

*MetricSecondaryStructure, tica.project(1), tica.lag(10ns):*

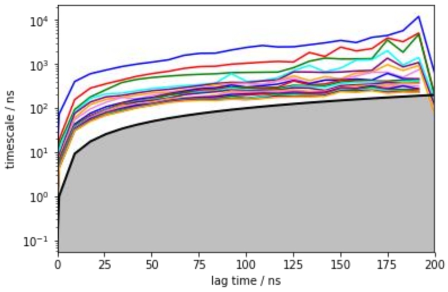

*MetricSecondaryStructure, tica.project(1), tica.lag(25ns):*

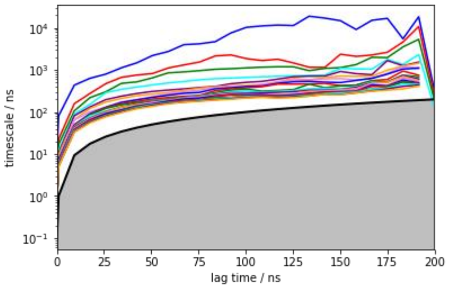

*MetricSecondaryStructure, tica.project(3), tica.lag(10ns):*

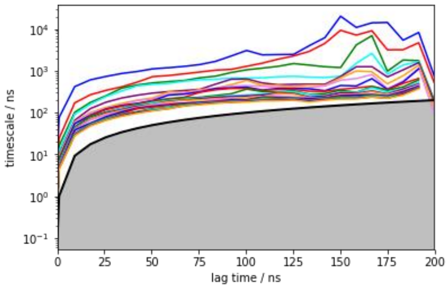

*MetricRmsd+SelfDistance, tica.project(1), tica.lag(10ns):*

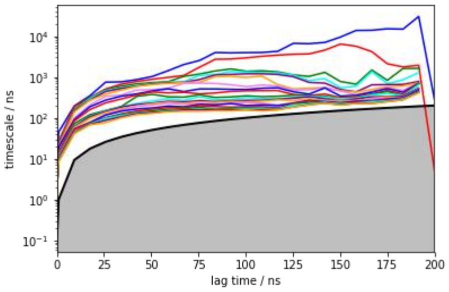

*MetricRmsd+SelfDistance, tica.project(5), tica.lag(10ns):*

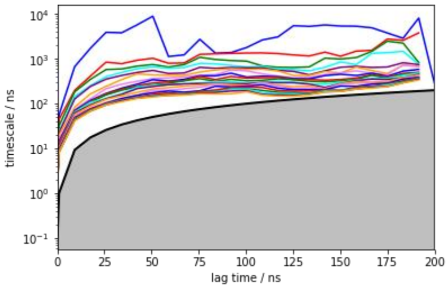

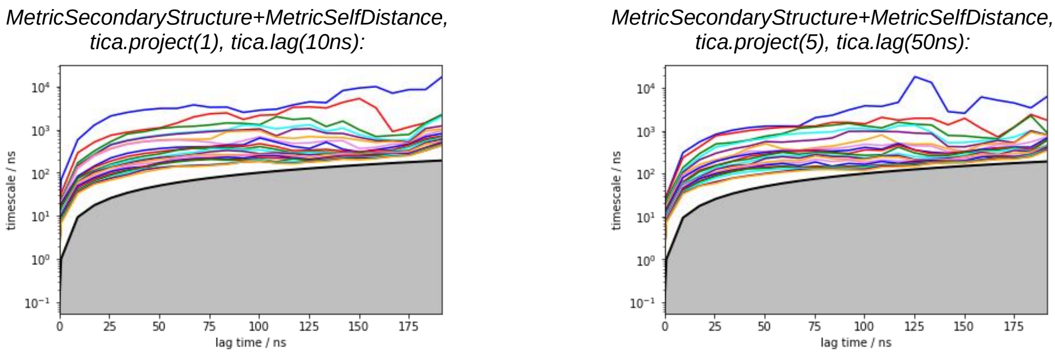

**Supplementary Figure S12.** Implied time scales (ITS) obtained for MSMs constructed for the free Aβ42 system with different settings, by varying: 1) metrics (*MetricRmsd* = RMSD of Cα atoms, *MetricSelfDistance* = self distance of all Cα atoms against all Cα atoms, *MetricSecondaryStructure* = secondary structure simplified for three types; 2) number of components of the tICA projections (*tica.project*); and 3) tICA lag times (*tica.lag*). The respective Chapman-Kolmogorov tests for the state transitions also failed for all of them (estimated ≠ predicted curves; not shown here).

**Variational Markov state analysis using VAMPnets**

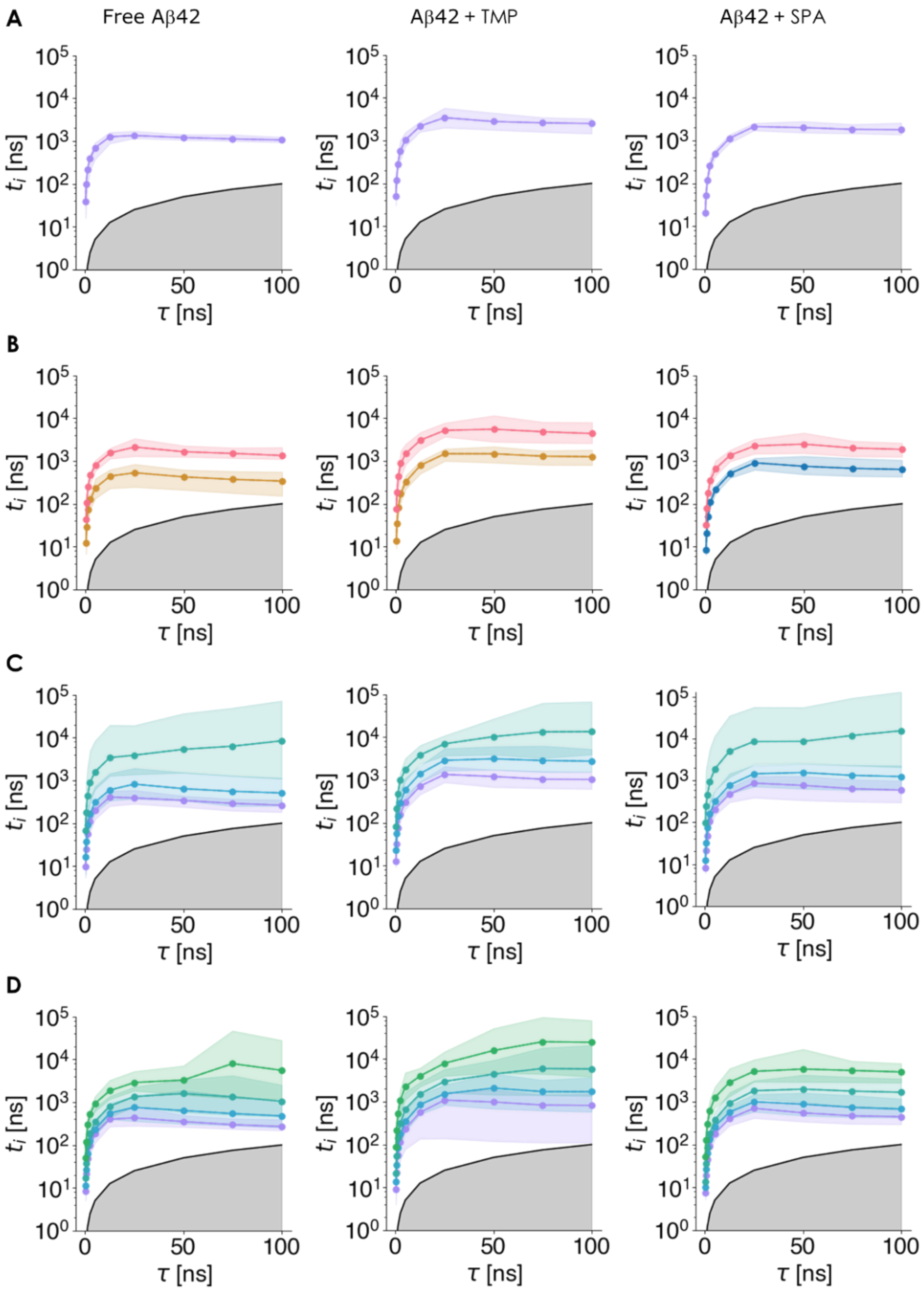

**Supplementary Figure S13.** Implied time scales of the slowest processes for different numbers of states calculated by the variational Markov state analysis using the VAMPnet approach. Implied timescales for: A) 2 state, B) 3 state, C) 4 state and D) 5 state MSMs. Data is presented for all systems: free Aβ42 (left), Aβ42 + TMP (center) and Aβ42+ SPA (right). The vertical axis shows log-scaled implied timescales, the horizontal axis shows the lag times used for the estimation of the Koopman operators from which the implied timescales were computed. The colored shading corresponds to the 2.5 to 97.5 percentile of the values obtained from the respective ensemble of 20 models (see **Supplementary Note 7**). The gray area cannot be resolved by a Koopman model. The implied timescales should not depend on the model lag time, therefore only the models with lag times, for which the implied timescales are approximately constant, should be considered. Panel B) corresponds to the Markov state models reported in the main text.

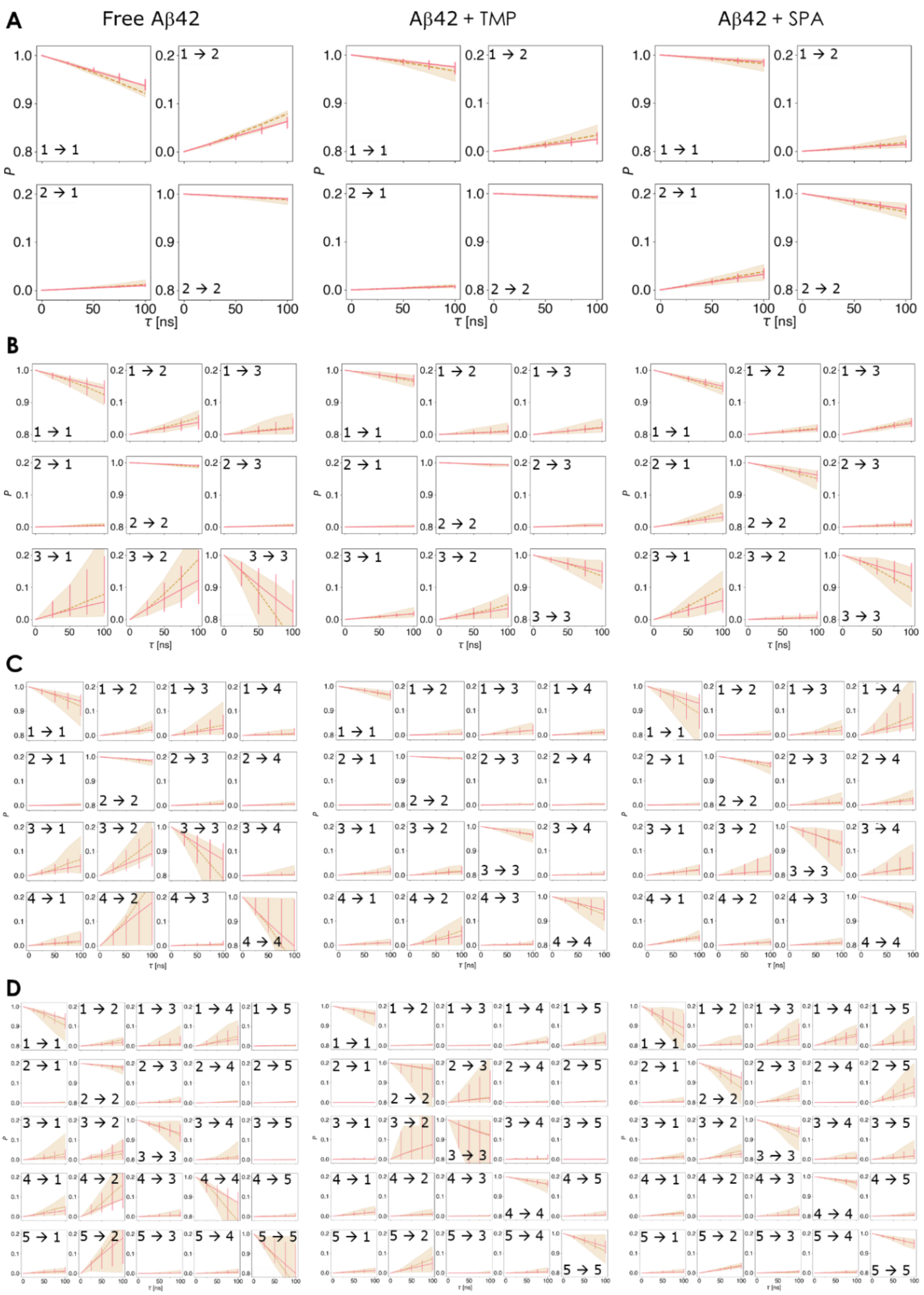

**Supplementary Figure S14. Chapman-Kolmogorov (C-K) tests of Markovianity** for: A) 2 state, B) 3 state, C) 4 state, and D) 5 state MSMs, estimated by the variational Markov state analysis using the VAMPnet approach. Data is presented for all systems: free Aβ42 (left), Aβ42 + TMP (center) and Aβ42+ SPA (right). The elementary plots labeled by "i->j" correspond to the C-K test for a transition from state *i* to state *j* within the estimated MSM: here the red line denotes the time dependency of the propagated proportion of the state population estimated from a single Koopman operator (built under assumption that the lag time $\tau$ = 25 ns) and its corresponding powers and the dashed line represents the estimates obtained from Koopman operators built for particular lag times (i.e. $\tau$ in {25 ns, 50 ns, 75 ns, 100 ns}). Both the red error bars and the shaded areas correspond to 2.5 to 97.5 percentile of the values obtained from the respective ensemble of 20 models (see **Supplementary Note 7**).

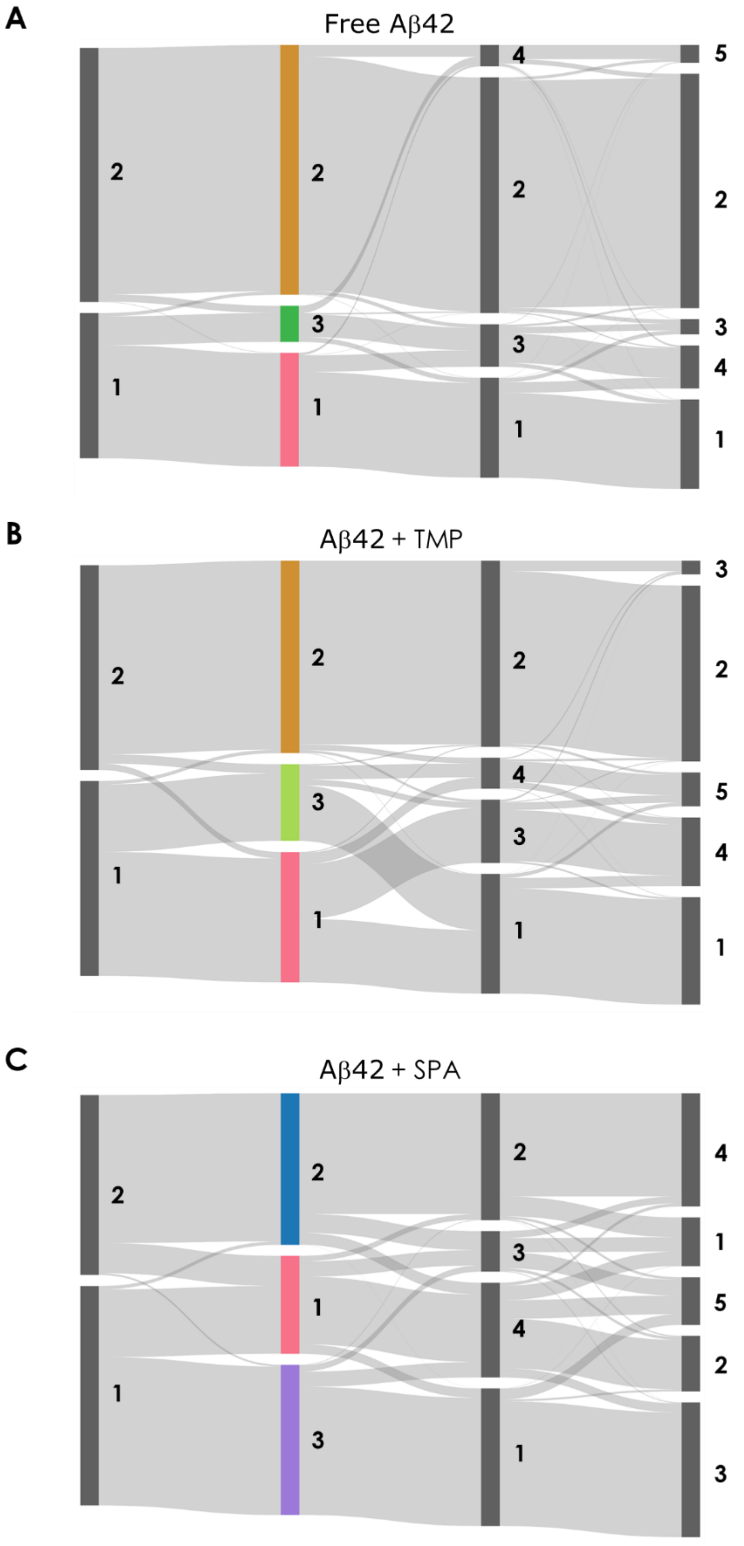

**Supplementary Figure S15. Change in the classification of frames by using VAMPnet estimated MSMs with varying numbers of states.** For A) Free Aβ42, B) Aβ42 + TMP, C) Aβ42 + SPA. The different states are numbered, and for the 3-state MSMs they are color-coded as in the main manuscript. The thickness of the shaded belts connecting two states in the consecutive model-ensembles is proportional to the amount of frames that is classified into the two connected states by the respective ensemble of 20 models.

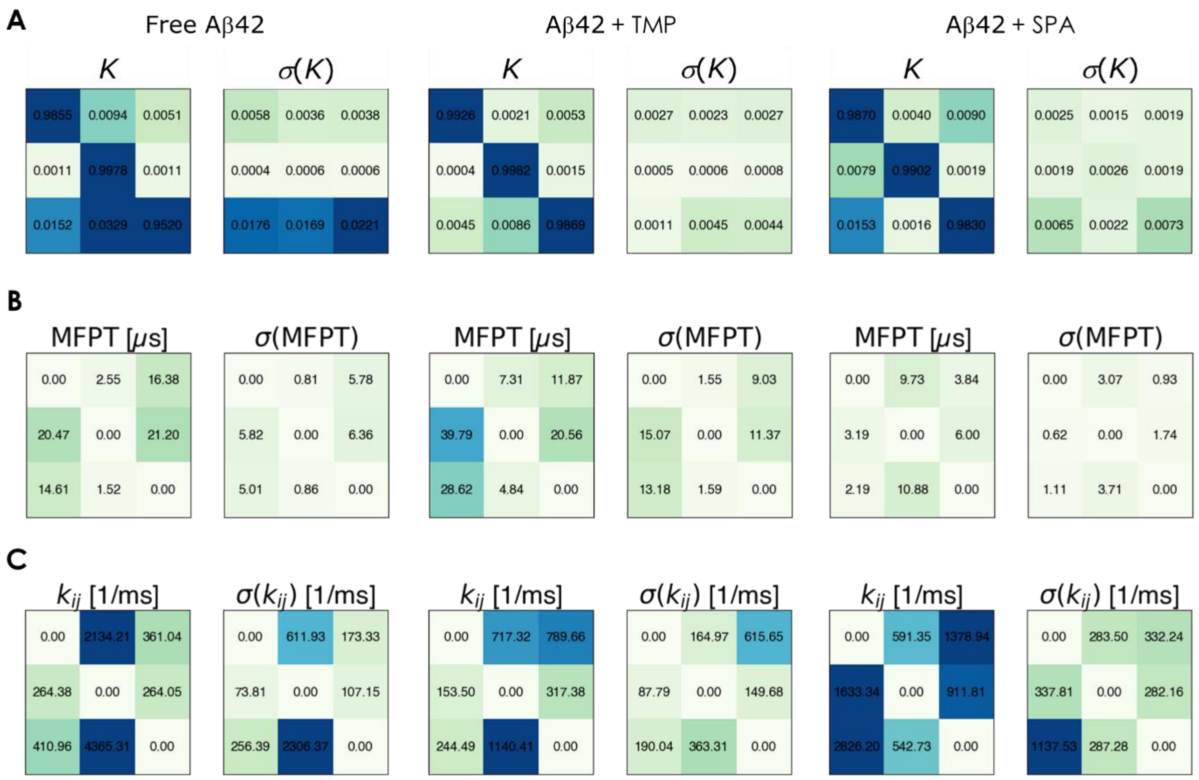

**Supplementary Figure S16. Properties of the MSMs calculated from the variational Markov state analysis using the VAMPnet approach.** A) Matrix representation of the Koopman operator (*K*), B) mean first-passage times (MFPT, also known as $T_M$), and C) transition rates ($k_{ij}$) among the different states. The respective standard deviations ($\sigma$) are presented. The transitions between states *i* → *j* correspond to the values in row *i* and column *j*. The results are presented for free Aβ42 (left), Aβ42 + TMP (middle), Aβ42 + SPA (right). All values are ensemble averages, therefore it does not hold that the rates in subfigure C are the inverse to the MFPTs in subfigure B, because that only holds for values estimated from individual models not for averaged values. For MFPTs and rates the "self-transitions" corresponding to the diagonal elements are not defined. Note that the matrix representation of the Koopman operator K, could be considered a probability transition matrix if $\chi$ were hard state assignments.

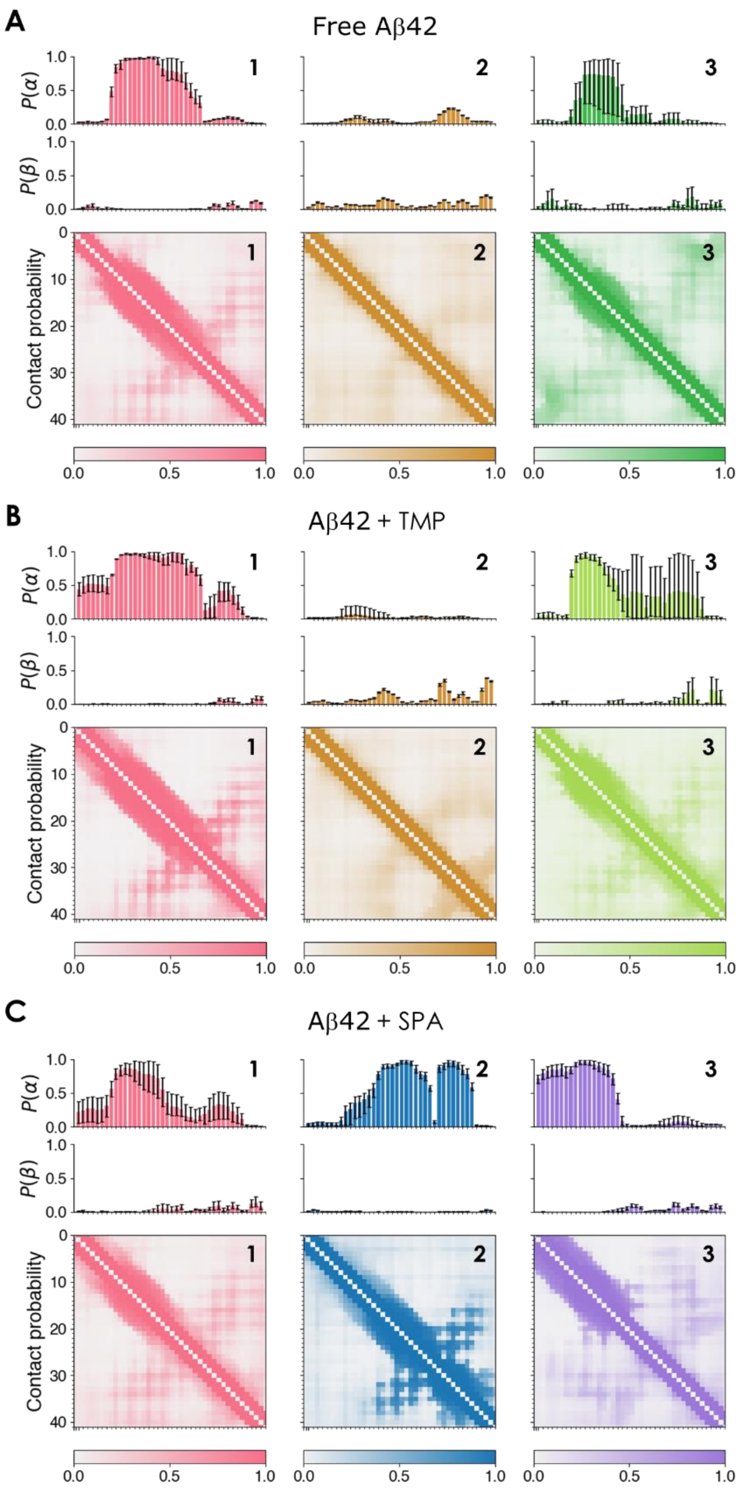

**Supplementary Figure S17. Probabilities of residues being part of the secondary structure element (top of subfigures) and average probabilities of residues being in contact (bottom of subfigures).** A) Free Aβ B) Aβ42 + SPA, and C) Aβ42 + SPA. Data are presented for all states in those systems: state 1 (left), state 2 (center) and state 3 (right). The secondary structure probabilities are shown for α-helices ($P(\alpha)$ histograms) and β-strands ($P(\beta)$ histograms), the error bars in these plots correspond to the 2.5-97.5 percentile of values across the ensemble of 20 models (see **Supplementary Note 7**). The data is colored according to the color coding of the states in **Figure 2** in the main manuscript.

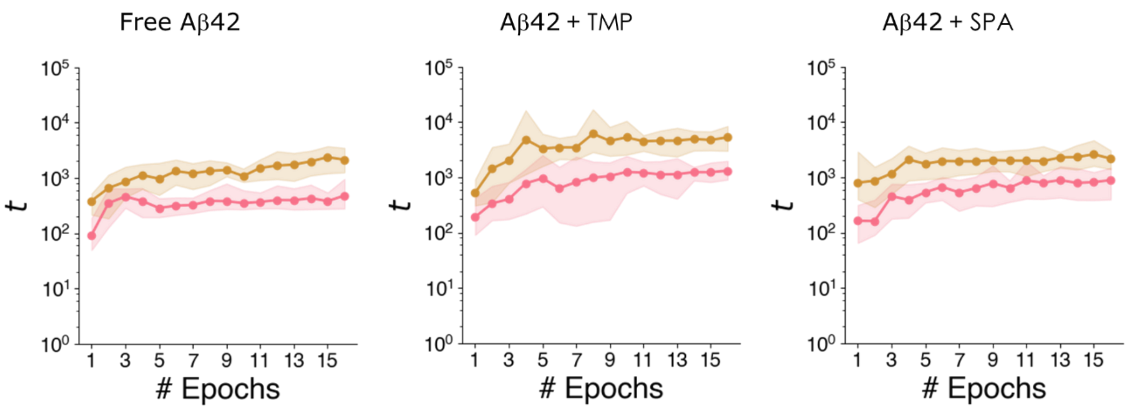

**Supplementary Figure S18. Dependency of estimated implied timescales on the size of the training dataset** for the 3 state MSMs for: free Aβ42 (left), Aβ42 + TMP (center) and Aβ42 + SPA (right). The sizes of the datasets were incrementally increased by adding whole simulation epochs. The shaded area corresponds to the 2.5-97.5 percentile of values obtained from the 20 models in the ensemble (see **Supplementary Note 7**). These plots were used for verification that the sizes of our datasets were sufficient, based on the plotted function being close to constant around our maximum dataset size.

### Comparative Markov state model analysis

| Aβ42 + TMP / Free Aβ42 | State 1 | State 2 | State 3 |
|---|---|---|---|
| State 1 | **4.78** | 9.68 | 5.11 |
| State 2 | 11.27 | **5.66** | 9.26 |
| State 3 | 15.38 | 16.37 | **12.56** |

| Aβ42 + SPA / Free Aβ42 | State 1 | State 2 | State 3 |
|---|---|---|---|
| State 1 | **4.15** | 6.63 | 8.29 |
| State 2 | 8.24 | **10.26** | 10.21 |
| State 3 | 10.66 | 15.16 | **10.44** |

**Supplementary Figure S19. Cost of alignment of the ensembles of states** between free Aβ42 and Aβ42 + TMP (top) and between free Aβ42 and Aβ42 + SPA (bottom). The listed scores correspond to the Wasserstein-1 distances of the distributions of the features of the states (see Equations 1-3 in the main paper), highlighted values correspond to values considered in the solution to the optimization problem of the alignment. Values highlighted in green are below the alignment threshold of 6 and therefore the corresponding states are considered to be aligned. Values highlighted in red are above the alignment threshold and the corresponding states are not considered to be aligned. The states are color-coded the same way as in **Figure 2** in the main manuscript.

#### Time-based evolution of the states

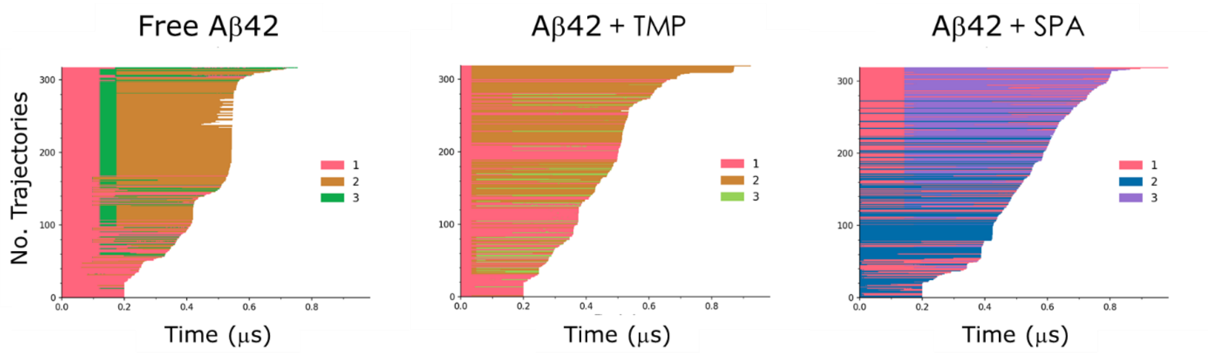

**Supplementary Figure S20. Hard assignment of concatenated adaptive sampling simulations into 3 states by VAMPnets.** Free Aβ (left), Aβ + TMP (middle), Aβ + SPA (right). The concatenated trajectories are ordered by the time of the first frame of their last simulation. White patches denote missing parts of incomplete simulations. We found that some states were more prominent in the early simulations and others in the late ones. In general, there were not many apparent transitions within the individual simulations. On the other hand, many segments were frequently repeated in the concatenated trajectories, which was due to the criterion applied by the adaptive method for the selection of the seeding frames.

#### Radius of gyration by state

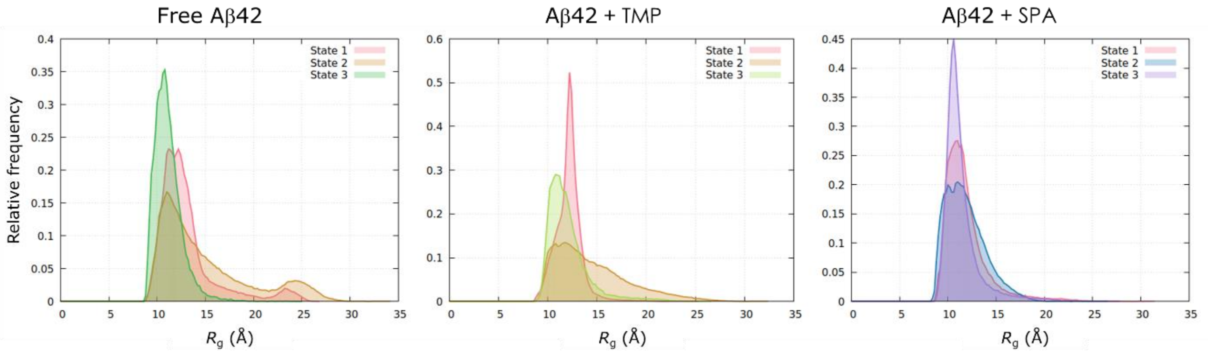

**Supplementary Figure S21. Distribution of the radius of gyration ($R_g$) of the different states from the different systems.** The results are presented for free Aβ (left), Aβ + TMP (middle), Aβ + SPA (right). The different states are numbered and color-coded as in the main manuscript.

### Characterization of learned conformational states via network gradients

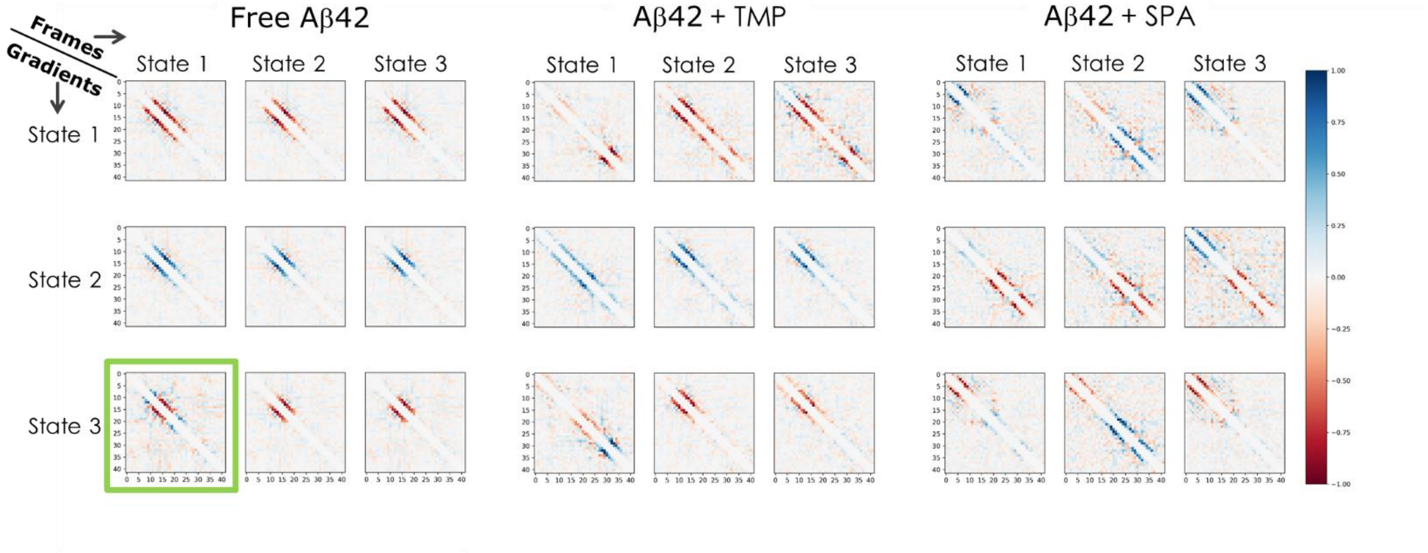

**Supplementary Figure S22. Visualization of the gradient-based analysis of the learned states for the 3-state MSMs with regard to the state assignments of the frames used for evaluation.** MSMs were analyzed for: free Aβ42, Aβ42 + TMP, and Aβ42 + SPA. In each subfigure, the rows show the ensemble-averaged gradients of the probabilities corresponding to state 1 (top), state 2 (center) and state 3 (bottom). Each column then denotes the state assignment of the frames, which were used for the evaluation of the gradients - this is in contrast with **Figure 3** (in the main manuscript) where the frames were used for evaluation regardless of their state assignment. Selecting frames for the evaluation based on their state assignment enables us to directly see, which residue distances should be changed for a frame to change its membership from a particular state into another selected state. This facilitates the pairwise state comparison. See, for example, subfigure for free Aβ42, where the lower left gradient visualization (highlighted by the green box) suggests that while part of the α-helix is shared by both state 1 and 3, it is the extent of this helix which makes the difference.

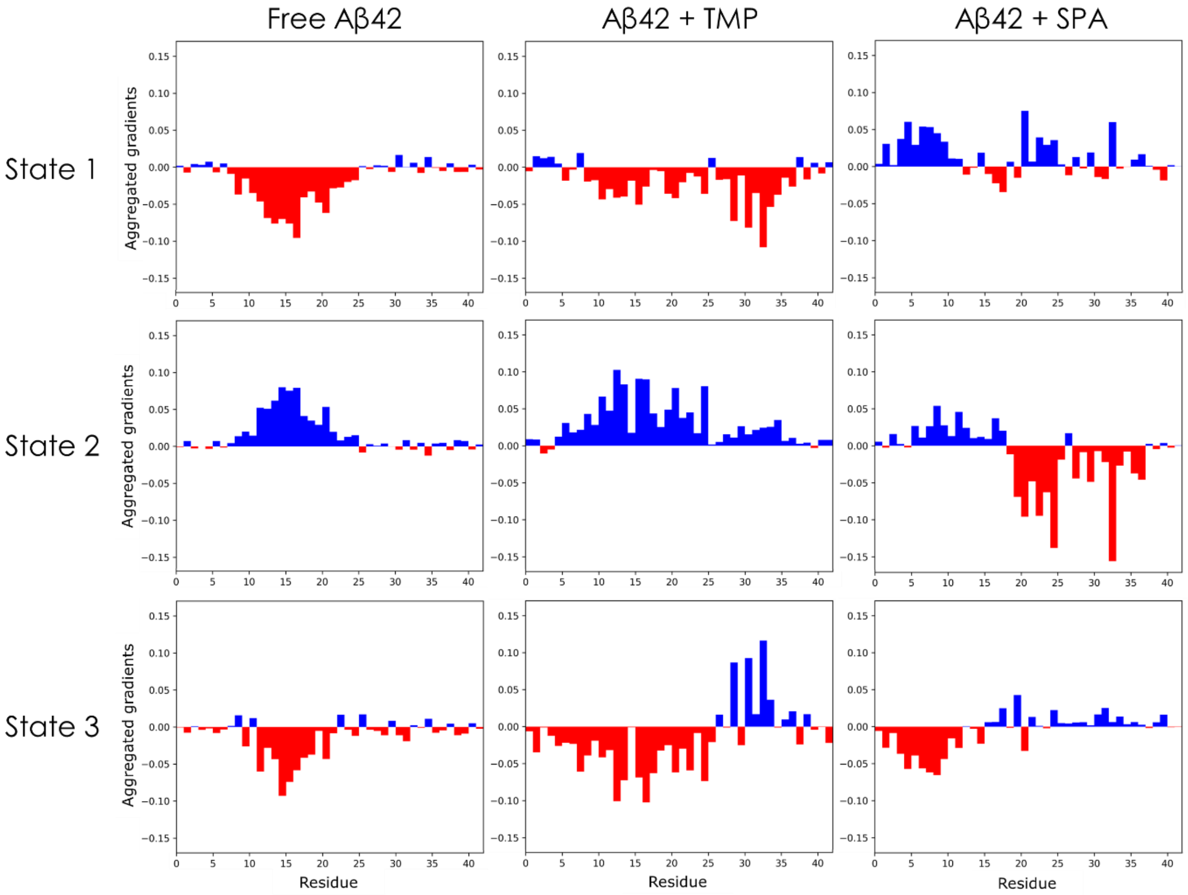

**Supplementary Figure S23. Aggregated per-residue gradients of the state assignment probabilities of the learned variational Markov state models.** Each plot shows the ensemble-averaged gradients of the model probabilities for the corresponding system (column) and state (row) with respect to the input inter-residue Cα distances (see **Figure 3**) aggregated and averaged separately for each residue. A positive value (blue) for a given residue indicates that the probability of the state assignment would increase if the average distance between the Cα atom of this residue and the Cα atoms of the other residues (except the first and second neighbors) increased whereas a negative value (red) indicates that the probability would increase if that average distance decreased; the higher the absolute value, the higher the effect on the probability. The plots were computed from the gradient matrices in **Figure 3** by averaging all non-empty values in each row (i.e., without the diagonal and the first and second subdiagonals). Columns: MSMs for the free Aβ42 (left), Aβ42 + TMP (middle), and Aβ42 + SPA (right) systems. Rows: states 1 (top), 2 (middle), and 3 (bottom) of each model. **Interpretation**: Large positive or negative values mark residues identified by the CoVAMPnet as key for classification into the states of each system. By comparing the sign and magnitude of aggregated gradients for the residues across the systems, one can easily identify the main structural differences between the states, and hence between the global properties of the systems. Insights obtained from this analysis confirm the intuitive understanding of the differences between the states, which relies on comparing the propensity of different secondary structure motifs and their position in the protein sequence. For example, the regions with significant negative values (red) in this plot are consistent with the regions prone to formation of α-helices (reasoning for this is provided in the *Characterization of learned conformational states via network gradients* section in the main manuscript).

**Interactions of Aβ42 with the small molecules**

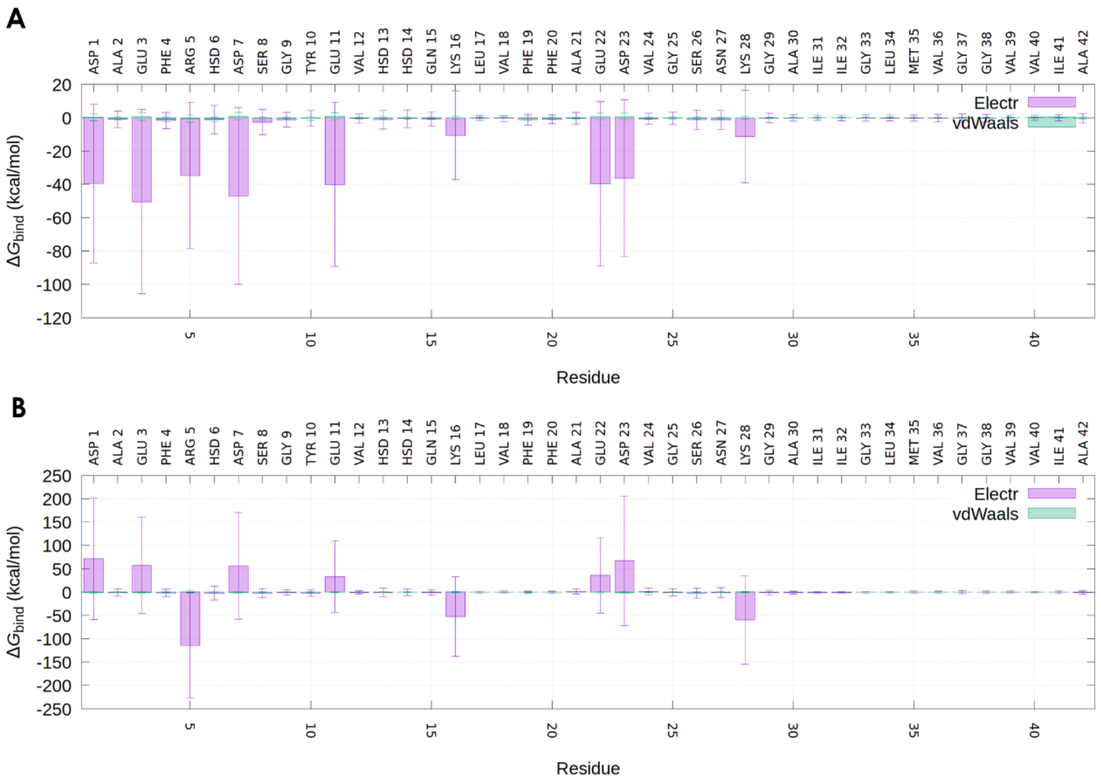

**Supplementary Figure S24. Interactions of TMP (A) and SPA (B) with Aβ42.** Mean value of the binding energy ($\Delta G_{bind}$) of TMP or SPA with each residue of Aβ42, calculated for of all the 100 molecules in every snapshot of the respective adaptive simulations, and dissected by the electrostatic (Electr) and van der Waals (vdWaals) components. The error bars represent the standard deviations from the mean interaction energies on a linear scale.

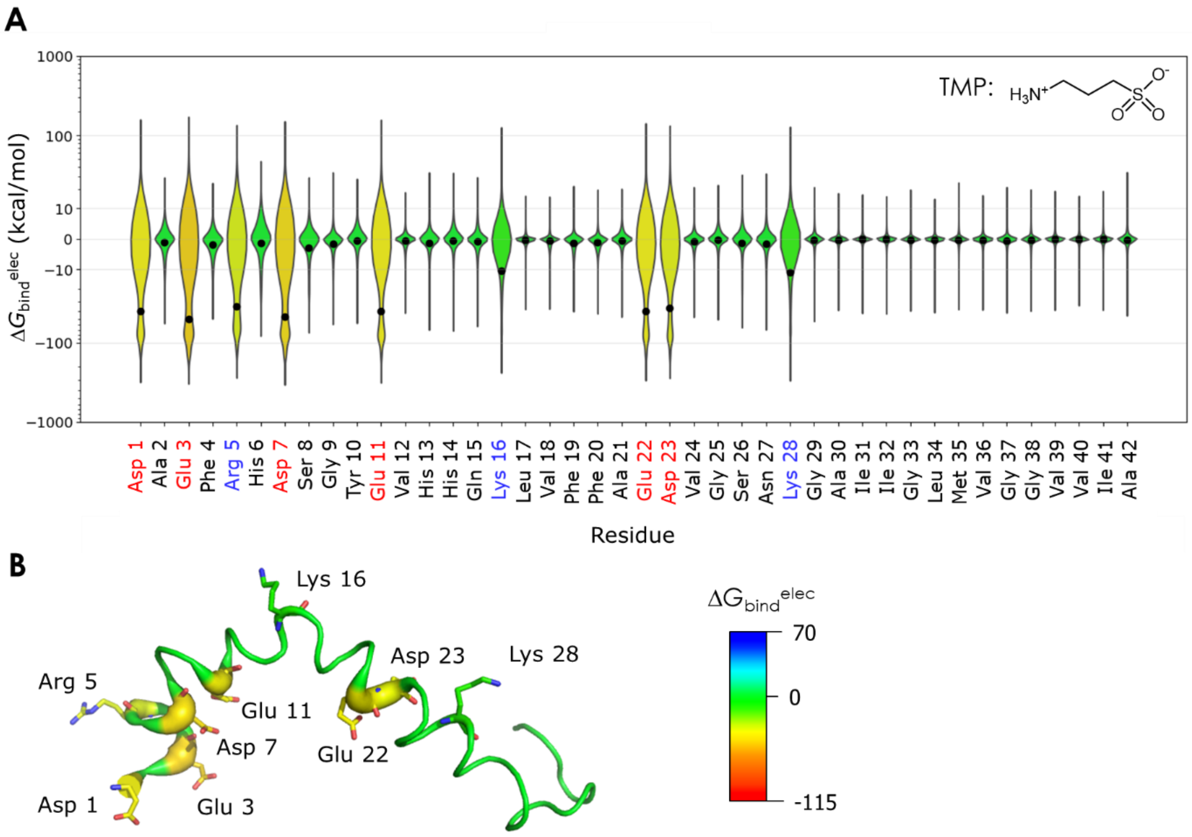

**Supplementary Figure S25. Interactions of TMP with Aβ42 studied by molecular dynamics. A) Violin plot of the binding energy of TMP with each residue of Aβ42**. The electrostatic component ($\Delta G_{\mathrm{bind}}^{\mathrm{elec}}$) was calculated for all the 100 molecules in every snapshot of the adaptive simulation of Aβ42 + TMP. The plot shows the distribution of the energy values; the black dots show the mean values; the y-axis uses a quasi-logarithmic scale based on the inverse hyperbolic sine to highlight the higher absolute values. The residue labels are colored by charge: black for neutral, blue for positive, and red for negative. The chemical structure of TMP is shown in the upper-right corner. **B) Structure of Aβ42 with the main interacting residues**. Aβ42 is shown as the putty cartoon and the main interacting residues are represented by sticks. The colors reflect the mean $\Delta G_{\mathrm{bind}}^{\mathrm{elec}}$ (in kcal/mol), and range from the most positive (blue) to the most negative (red) values obtained for SPA (see **Figure 4**).

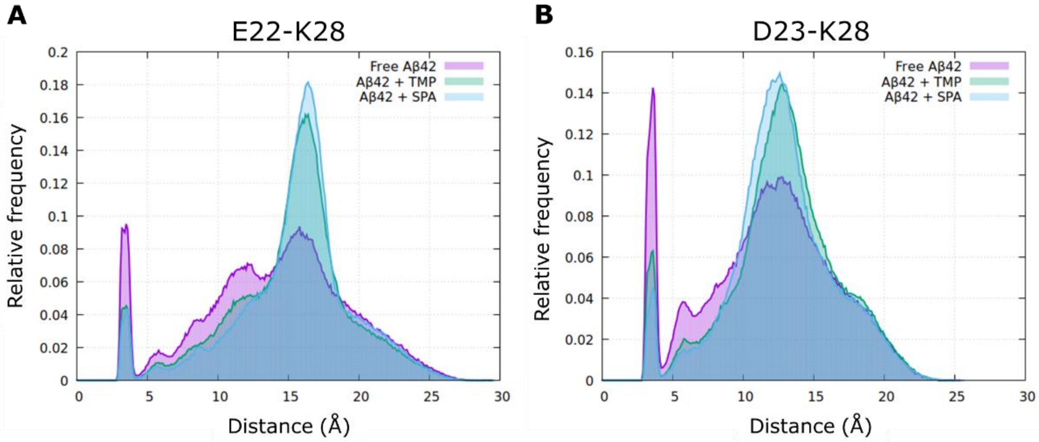

**Supplementary Figure S26. Salt-bridges in Aβ42.** Distribution of the distances between the amine N-atom of Lys28, and A) the carboxyl-C atom of Glu22, or B) the carboxyl-C atom of Asp23 for the different systems. The peaks around 3.6 Å correspond to the formed salt bridges, which were more abundant for the free Aβ42.

**Experimental validation**

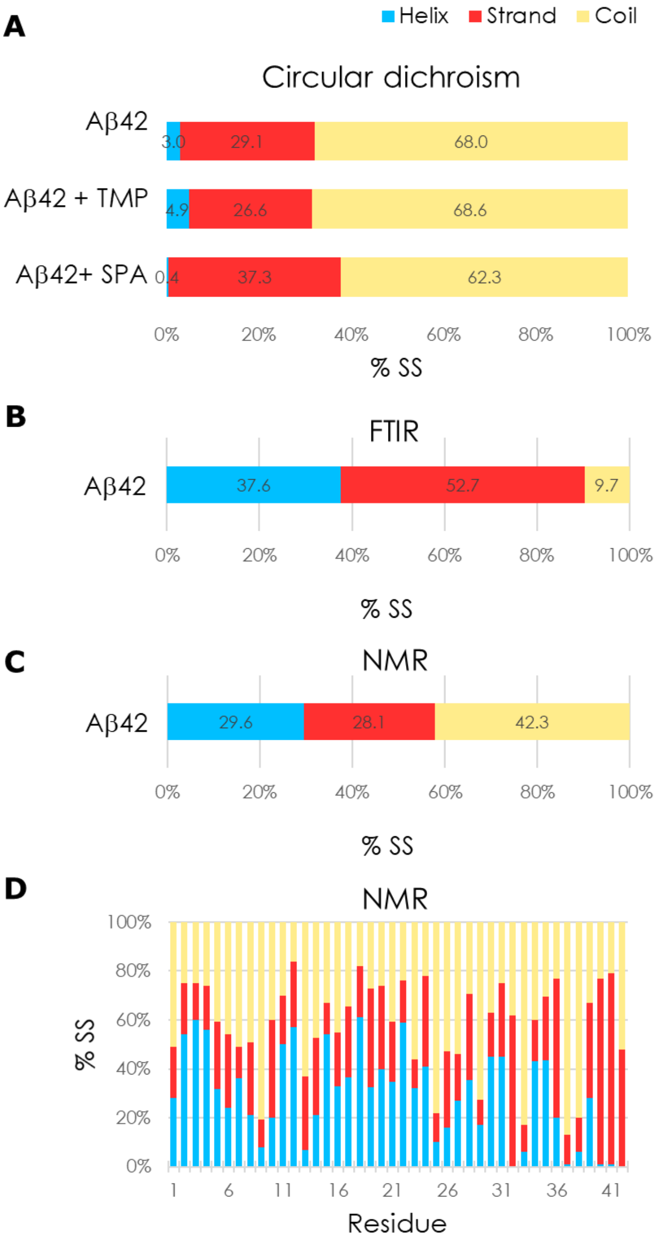

**Supplementary Figure S27. Secondary structure of N-Met-Aβ42 obtained from different experimental techniques, in buffer alone and in the presence of 1000-excess of small molecules.** The total secondary structure content (% SS) from circular dichroism (A), FTIR (B) and from NMR (C), and content by residue from NMR (D).

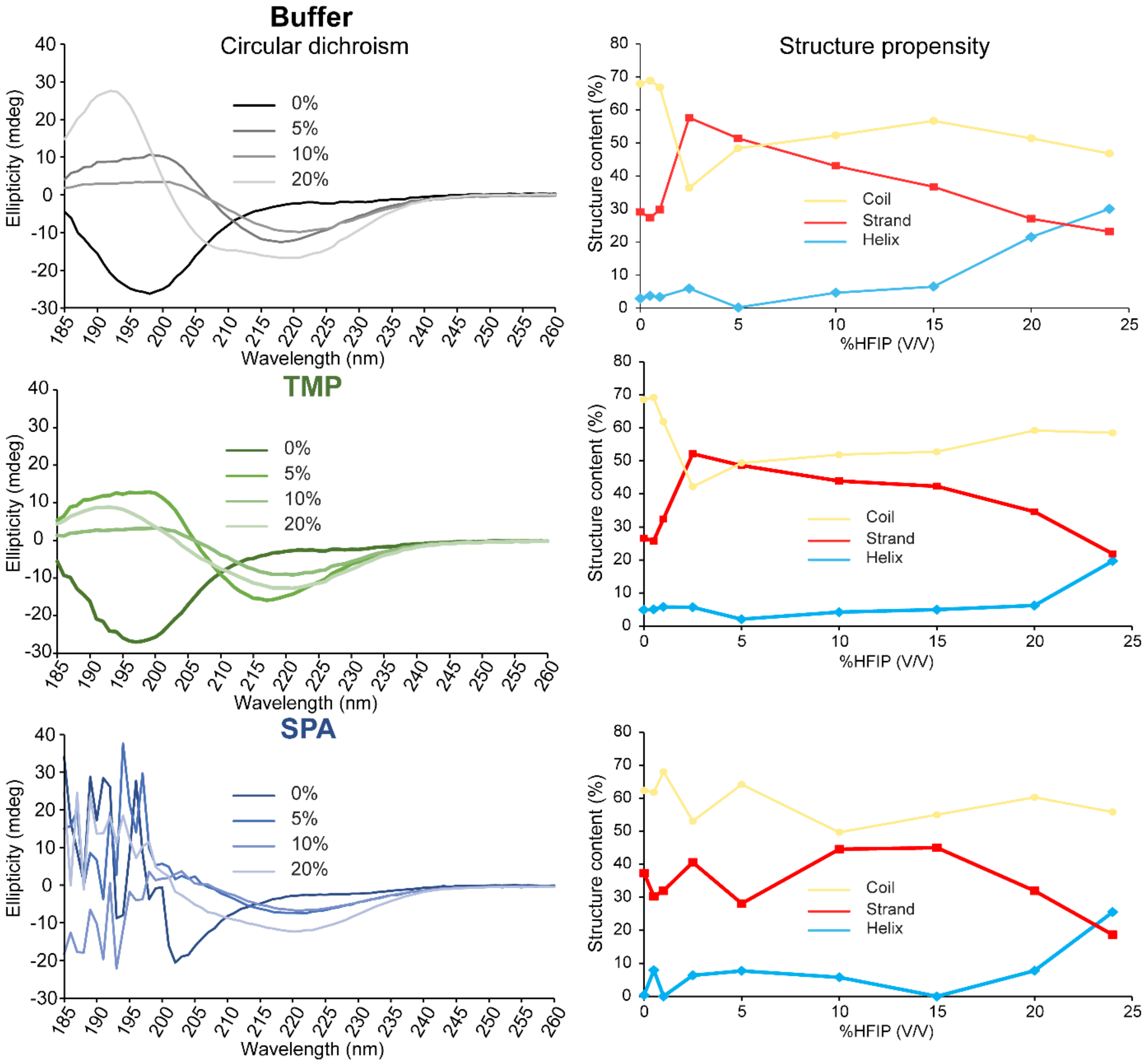

**Supplementary Figure S28. CD spectroscopy of N-Met-Aβ42 in the presence of drug candidates and HFIP.** *Left*: Titration of 37 µM of free N-Met-Aβ42 (top), or with 1000-fold excess of TMP (center) or SPA (bottom) with different ratios of HFIP (volume percentages). *Right*: respective secondary structure content computed by BestSel based on the CD data in the absence of drugs (top), or with 1000-fold excess of TMP (center) or SPA (bottom).

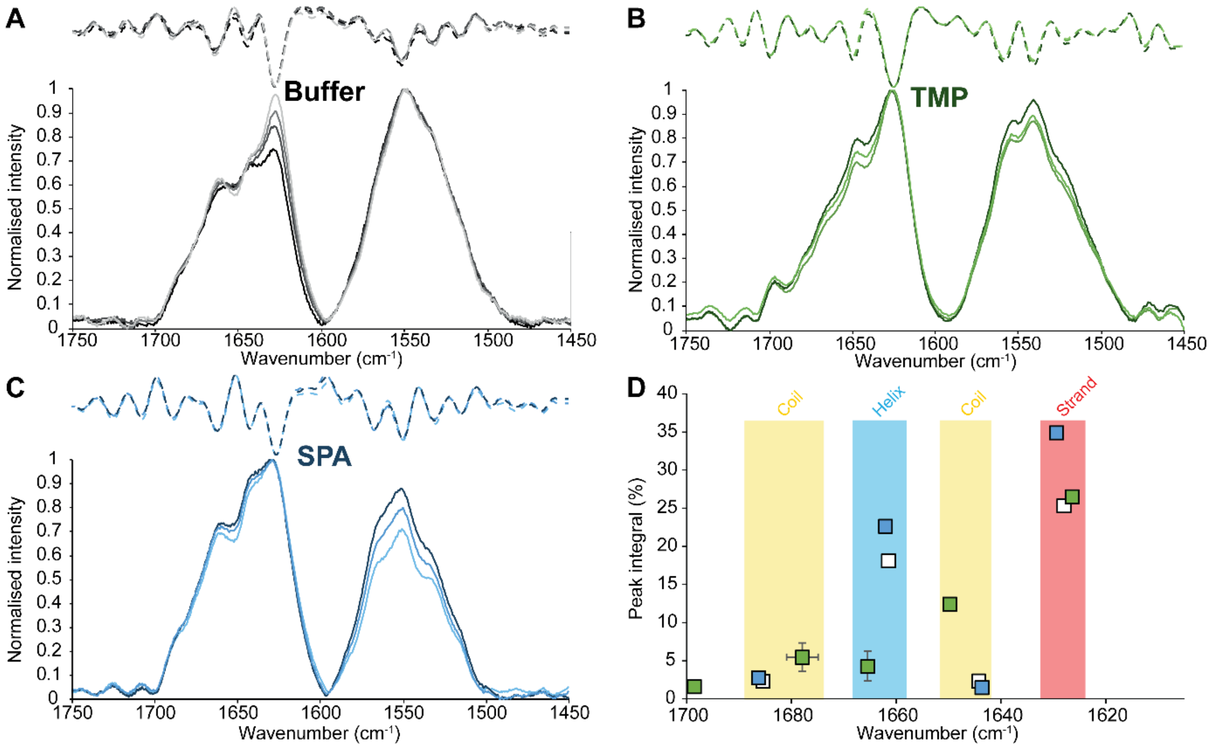

**Supplementary Figure S29. FTIR spectroscopy of N-Met-Aβ42.** FTIR spectra (solid curves) and corresponding secondary derivatives (dashed curves) of the free N-Met-Aβ42 in buffer (A) or with a 1000-fold excess of TMP (B) or SPA (C). The shades represent successive acquisitions on the same sample (darkest: oldest, lightest: latest), indicating the sample evolution after deposition on the ATR crystal. The corresponding secondary derivatives are plotted as dashed curves. (D) Gaussian deconvolution of the FTIR spectra (A-C) and subsequent assignment of the respective total secondary structure content.

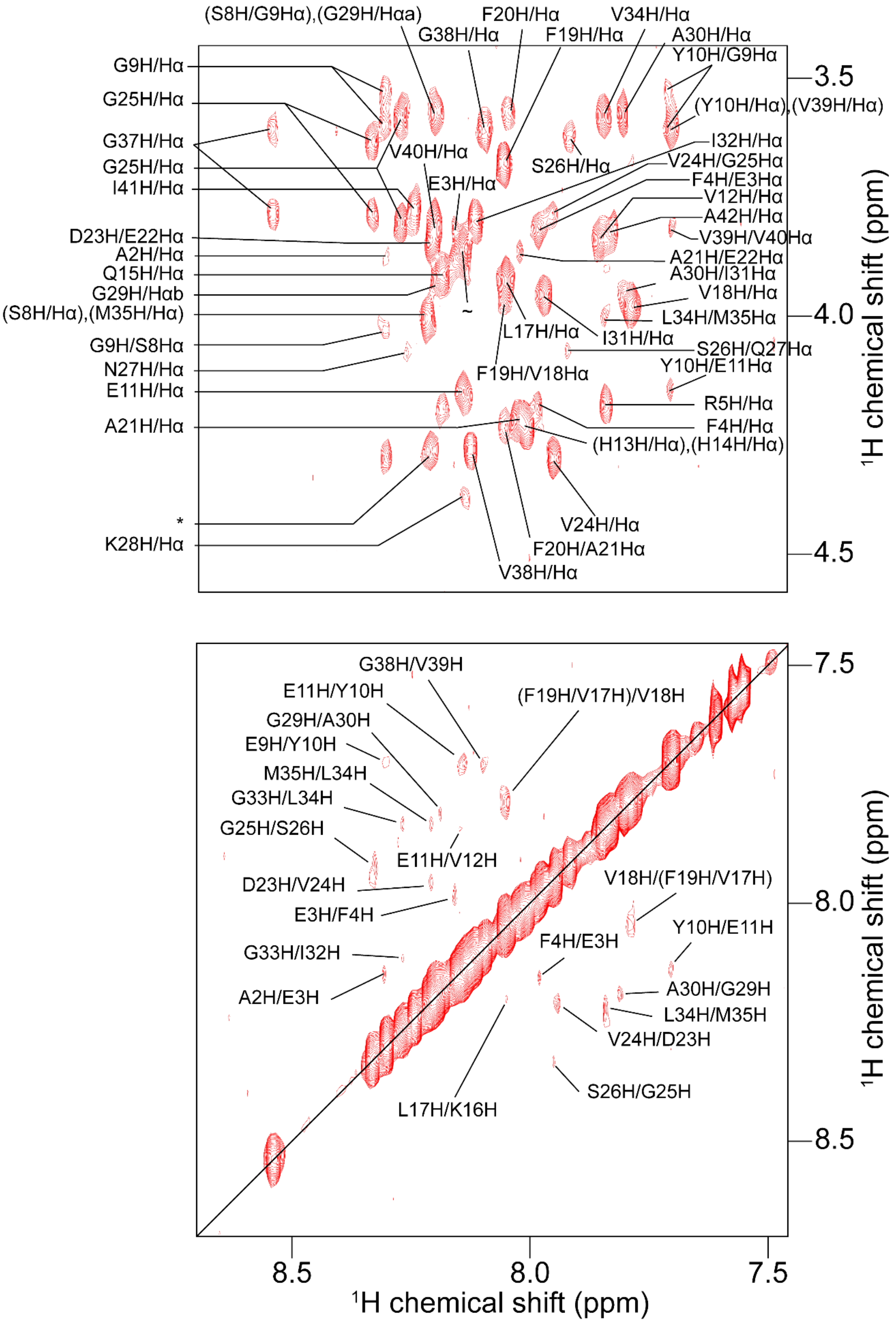

**Supplementary Figure S30. Assigned $^{1}$H-$^{1}$H NOESY spectrum of N-Met-Aβ42 in 20 mM sodium phosphate at pH 7.4 at 4 °C.** Only unambiguous assignments are shown. *: (S8H/S8Hα), (D7H/D7Hα), (D23H/D23Hα). ~: (H6H/H6Hα), (E22H/E22Hα).

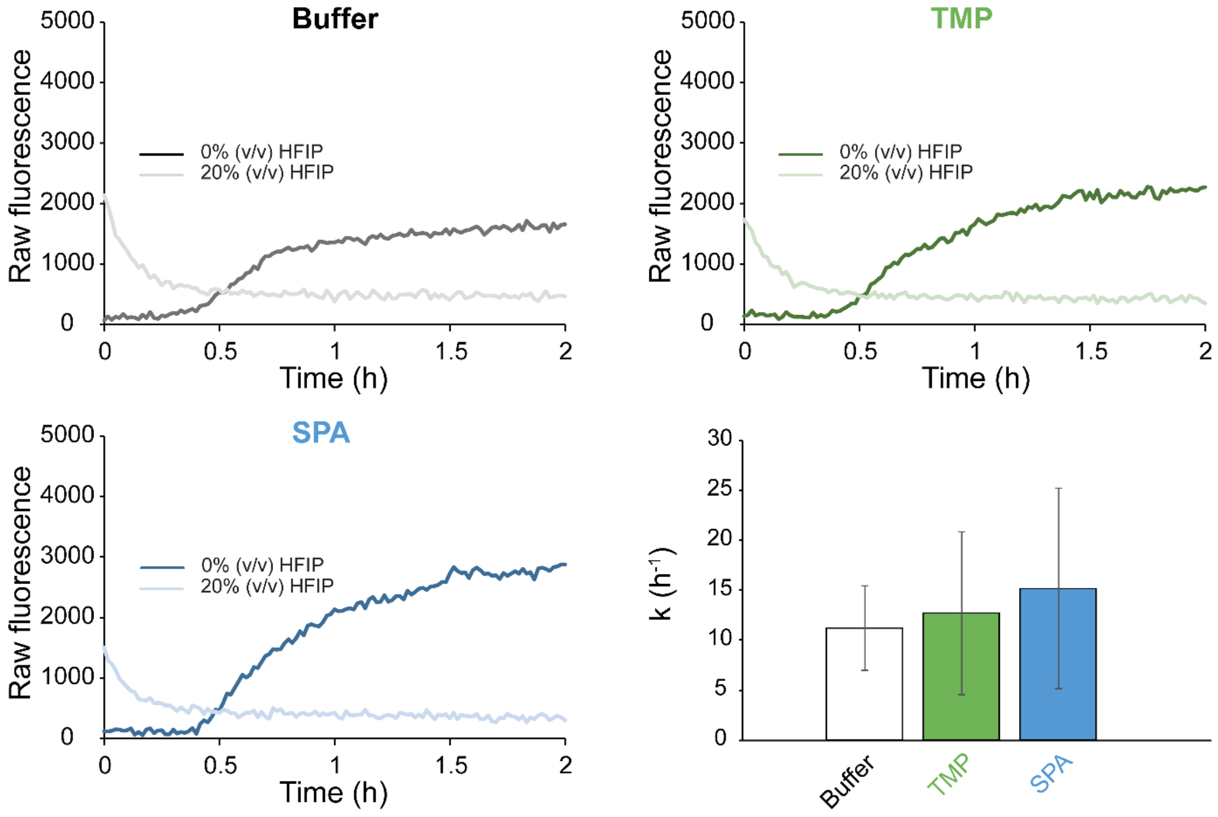

**Supplementary Figure S31. ThT-based fibril formation assay on N-Met-Aβ42.** 10 µM N-Met-Aβ42 were mixed with 15 µM Tht (A) and 10 mM of TMP (B) or SPA (C), in the absence and presence of HFIP (0% and 20% HFIP (v/v)). (D) Apparent fibril formation rate, assuming a logistic growth. The error bars are the standard deviations from triplicates.

### SUPPLEMENTARY TABLES

**Supplementary Table S1. Selection of a computational protocol for the simulation of Aβ42.** Summary of all tested adaptive sampling protocols.[a]

| | Protocol A | | Protocol B | Protocol C |
|---|---|---|---|---|
| **Force Field** | **A14SB** | **C36m** | **C36m** | **C36m** |
| **Single MD length** | 50 ns | 50 ns | 50 ns | 200 ns |
| **# initial structures** | 30 (all) | 30 (all) | 1 (first) | 1 (first) |
| **Sampling metric** | Self-distance (Cα) | Self-distance (Cα) | Self-distance (Cα) | Secondary structure |
| **tICA projection** | 1 dimension | 1 dimension | 1 dimension | 1 dimension |
| **# initial epochs** | 10 | 20 | 12 | 16 |
| **# initial replicas** | 4 | 4 | 10 | 20 |
| **# final epochs** | NA | NA | 68 | NA |
| **# final replicas** | NA | NA | 20 | NA |
| **Total cumulative MD time** | 60 µs | 120 µs | ca. 74 µs | ca. 64 µs |
| **Systems applied** | Free Aβ42 | Free Aβ42 | Free Aβ42 | Free Aβ42, Aβ42 + TMP, Aβ42 + SPA |

[a]The different combinations of adaptive sampling protocol and force field are listed in columns, rows showing the different adaptive sampling parameters used for each combination. NA means “not applicable”.

**Supplementary Table S2. Analysis of classical MDs.** Total secondary structure propensities in the classical MD replicates for the free Aβ42, Aβ42 + TMP and Aβ42 + SPA.[a]

| **Free Aβ42** | **MD1** | **MD2** | **MD3** | **MD4** | **MD5*** | **MD6** | **MD7** | **MD8** | **MD9** | **MD10** | ***AVG ± SEM*** |
|---|---|---|---|---|---|---|---|---|---|---|---|
| **Helix** | 5.0 | 27.7 | 8.8 | 31.8 | **15.5** | 3.8 | 16.4 | 7.2 | 7.6 | 43.3 | *16.7 ± 4.0* |
| **Strand** | 20.0 | 10.0 | 17.0 | 12.7 | **11.3** | 15.8 | 18.6 | 11.1 | 12.7 | 5.8 | *13.5 ± 1.3* |
| **Coil** | 75.0 | 62.3 | 74.2 | 55.5 | **73.2** | 80.4 | 64.9 | 81.7 | 79.6 | 50.9 | *69.8 ± 3.2* |
| ***RMSE*** | *8.3* | *7.9* | *5.6* | *12.0* | ***2.5*** | *9.8* | *4.1* | *8.9* | *7.8* | *19.3* | |

| **Aβ42 + TMP** | **MD1** | **MD2** | **MD3** | **MD4** | **MD5** | **MD6** | **MD7** | **MD8** | **MD9*** | **MD10** | ***AVG ± SEM*** |
|---|---|---|---|---|---|---|---|---|---|---|---|
| **Helix** | 11.0 | 30.4 | 43.9 | 36.6 | 13.9 | 43.7 | 10.4 | 11.7 | **16.1** | 8.1 | *22.6 ± 4.3* |
| **Strand** | 12.3 | 7.4 | 8.1 | 3.2 | 17.2 | 1.2 | 22.1 | 15.2 | **13.0** | 16.2 | *11.6 ± 2.0* |
| **Coil** | 76.7 | 62.2 | 48.0 | 60.1 | 68.9 | 55.1 | 67.5 | 73.1 | **70.9** | 75.7 | *65.8 ± 2.8* |
| ***RMSE*** | *9.2* | *5.6* | *16.2* | *10.0* | *6.2* | *15.0* | *9.4* | *7.8* | ***4.8*** | *10.5* | |

| **Aβ42 + SPA** | **MD1** | **MD2** | **MD3** | **MD4** | **MD5** | **MD6** | **MD7** | **MD8*** | **MD9** | **MD10** | ***AVG ± SEM*** |
|---|---|---|---|---|---|---|---|---|---|---|---|
| **Helix** | 28.6 | 47.3 | 3.3 | 43.2 | 6.6 | 15.8 | 4.9 | **17.0** | 2.7 | 19.3 | *18.9 ± 4.9* |
| **Strand** | 4.8 | 4.0 | 14.3 | 1.7 | 17.8 | 15.9 | 12.3 | **11.4** | 24.3 | 14.5 | *12.1 ± 2.1* |
| **Coil** | 66.6 | 48.7 | 82.3 | 55.1 | 75.6 | 68.4 | 82.8 | **71.6** | 73.0 | 66.2 | *69.0 ± 3.2* |
| ***RMSE*** | *7.1* | *20.7* | *11.9* | *17.2* | *8.7* | *2.9* | *11.4* | ***1.9*** | *12.0* | *2.2* | |

[a]Overall contents for all the 5 μs MDs (MD1-MD10), with the respective average and standard error of the mean (*AVG ± SEM*); *RMSE* is the root-mean-square error of the secondary content in each MD with respect to the average values. *The MDs with secondary propensities closest to the average over all the replicates (with lowest *RMSE*).

**Supplementary Table S3. Summary of the effects of small molecules.** Comparison of the physicochemical properties and the effects of TMP and SPA on the simulations with Aβ42.

| | **TMP** | **SPA** |
|---|---|---|
| **Structures** | 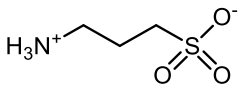 | 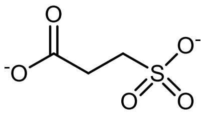 |
| **Charges at pH 7.4** | 1 ⊕, 1 ⊖<br>Total charge: 0 | 2 ⊖<br>Total charge: -2 |
| **Main interactions** | Attractions with all charged residues | Attractions with positively charged residues;<br>repulsions with negatively charged residues |
| **Polarity effects on residues** | Preserves polarity around charged residues | Inverts polarity around positively charged residues |
| **Shift in the overall secondary structure distribution of Aβ42 in adaptive sampling simulations[1)]** | +11.8% α-helices<br>~ same β-strands<br>-11.2% coils | +24.8% α-helices<br>-3.4% β-strands<br>-21.4% coils |
| **Change in the salt bridge formation rates in Aβ42[2)]** | -52% E22-K28<br>-56% D23-K28 | -55% E22-K28<br>-68% D23-K28 |
| **Radius of gyration[3)]** | Narrower distribution and smaller average than for free Aβ42;<br>$R_g$ = 13.3 ± 3.1 Å | Narrower distribution and smaller average than for free Aβ42 and Aβ42 + TMP;<br>$R_g$ = 11.8 ± 2.1 Å |
| **Effects on Markov state model ensembles and kinetics** | Ensemble distribution and kinetics more similar to free Aβ42;<br>predominant sink state as in the free Aβ42 (a disordered state) | Ensemble distribution and kinetics more divergent from free Aβ42;<br>predominant initial state (a helical-rich state);<br>more homogeneous FES |

[1)]Compared to the total %SS found for free Aβ42 (16.8% α-helices, 5.7% β-strands, 77.5% coils); [2)]with respect to the ratios found in the free Aβ42, which was referenced as 100%, and were computed from the height of the respective peaks near 3.5 Å in the distance distribution between the respective residues (**Supplementary Figure S26**); [3)]for free Aβ42 we obtained $R_g$ = 14.2 ± 4.3 Å.

**Supplementary Table S4. Intramolecular interactions of Aβ42.** Free energy (MM/GBSA) calculated for the Aβ42 peptide in the three systems, in the respective adaptive ensembles, and respective energy decomposition.[a]

| Energy Component | Free Aβ42 | | | Aβ42 + TMP | | | | Aβ42 + SPA | | | |
|---|---|---|---|---|---|---|---|---|---|---|---|
| | Average | SD | SEM | Average | SD | SEM | $\Delta\Delta E^b$ | Average | SD | SEM | $\Delta\Delta E^b$ |
| $\Delta E_{bond}$ | 155.5 | 11.2 | 0.02 | 156.2 | 11.3 | 0.02 | *0.7* | 156.4 | 11.3 | 0.02 | *1.0* |
| $\Delta E_{angle}$ | 331.8 | 15.6 | 0.02 | 333.5 | 15.6 | 0.02 | *1.7* | 335.5 | 15.4 | 0.02 | *3.7* |
| $\Delta E_{dihed}$ | 366.3 | 10.5 | 0.01 | 364.4 | 10.9 | 0.02 | *-1.9* | 363.8 | 9.7 | 0.01 | *-2.4* |
| $\Delta E_{UB}$ | 39.4 | 2.8 | 0.00 | 39.6 | 2.8 | 0.00 | *0.2* | 39.7 | 2.8 | 0.00 | *0.3* |
| $\Delta E_{IMP}$ | 24.4 | 3.7 | 0.01 | 24.6 | 3.7 | 0.01 | *0.2* | 24.8 | 3.7 | 0.01 | *0.4* |
| $\Delta E_{CMAP}$ | -50.3 | 10.1 | 0.01 | -45.7 | 12.8 | 0.02 | *4.6* | -41.1 | 9.0 | 0.01 | *9.3* |
| $\Delta E_{vanderWaals}$ | -166.3 | 26.3 | 0.04 | -177.4 | 26.9 | 0.04 | *-11.1* | -192.8 | 16.9 | 0.02 | *-26.5* |
| $\Delta E_{electrostat}$ | -2598.6 | 87.5 | 0.12 | -2565.3 | 84.6 | 0.12 | *33.2* | -2572.2 | 91.3 | 0.13 | *26.4* |
| $\Delta E_{1\text{-}4\ vanderWaals}$ | 114.0 | 6.4 | 0.01 | 116.4 | 7.6 | 0.01 | *2.4* | 119.0 | 6.0 | 0.01 | *5.0* |
| $\Delta E_{1\text{-}4\ electrost}$ | 2333.2 | 26.6 | 0.04 | 2328.4 | 27.5 | 0.04 | *-4.8* | 2321.4 | 25.4 | 0.04 | *-11.8* |
| $\Delta G_{polar\ solv}$ | -880.8 | 81.2 | 0.11 | -916.6 | 76.1 | 0.11 | *-35.8* | -911.3 | 84.1 | 0.12 | *-30.5* |
| $\Delta G_{non\text{-}polar\ solv}$ | 32.4 | 3.5 | 0.01 | 31.6 | 3.1 | 0.00 | *-0.8* | 30.0 | 2.2 | 0.00 | *-2.4* |
| $\Delta G_{total\ gas}$ | 549.4 | 97.5 | 0.14 | 574.6 | 95.0 | 0.13 | *25.3* | 554.6 | 94.0 | 0.13 | *5.2* |
| $\Delta G_{total\ solv}$ | -848.3 | 79.1 | 0.11 | -885.0 | 74.7 | 0.11 | *-36.7* | -881.2 | 83.5 | 0.12 | *-32.9* |
| $\Delta G_{total}$ | **-299.0** | **33.6** | **0.05** | **-310.4** | **39.2** | **0.06** | ***-11.4*** | **-326.6** | **27.2** | **0.04** | ***-27.7*** |

[a]Average energy contributions for: bonds ($\Delta E_{bond}$), angles ($\Delta E_{angle}$), dihedral ($\Delta E_{dihed}$), Urey-Bradley ($\Delta E_{UB}$), improper dihedrals ($\Delta E_{IMP}$), correction map ($\Delta E_{CMAP}$), van der Waals ($\Delta E_{vanderWaals}$), electrostatics ($\Delta E_{electrostat}$), 1-4 van der Waals ($\Delta E_{1\text{-}4\ vanderWaals}$), 1-4 electrostatics ($\Delta E_{1\text{-}4\ electrost}$), polar solvation ($\Delta G_{polar\ solv}$), non-polar solvation ($\Delta G_{non\text{-}polar\ solv}$), total gas phase free energy ($\Delta G_{total\ gas}$), total solvation free energy ($\Delta G_{total\ solv}$), and total free energy ($\Delta G_{total}$); [b]$\Delta\Delta E$ refers to the difference of the mean values of each energy component with those for the free Aβ42. SD is the standard deviation and SEM is the standard error of the mean. All the units are in kcal/mol.
